# Effects of point density on interpretability of lidar-derived forest structure metrics in two temperate forests

**DOI:** 10.1101/2024.01.11.575266

**Authors:** A. Christine Swanson, Trina Merrick, Andrei Abelev, Robert Liang, Michael Vermillion, Willibroad Buma, Rong-Rong Li

**Author notes:** Corresponding Author: A. Christine Swanson.

## Abstract

Three-dimensional forest structure plays an important role in processes such as biomass accumulation and fire spread and provides wildlife with habitat and foraging spaces. Advances in lidar mapping have improved forest structure quantification at local to global scales. However, point cloud density may have effects on estimates of forest structure variables that are not well understood and may vary by forest structural type (e.g. closed vs open canopy). In this study we investigated the effects of lidar point cloud density on forest structure parametrization in an open canopy pine-dominated forest at Assateague Island National Seashore (AINS) and a closed-canopy mixed hardwood temperate forest at the Keweenaw Research Center (KRC) using uncrewed aerial system (UAS)-based lidar. We decimated high point density (> 1000 points m^-2^) lidar data to between 1 and 175 points m^-2^ and analyzed 26 forest structure metrics using Tukey’s method, reliability ratio, and correlation analyses.

Effects of point density on forest structure parameters were often site-dependent, as anticipated. At AINS, maximum (*zmax*) and mode (*zmode*) height significantly differed for point densities less than 10 pts m^-2^ and 25 points m^-2^, respectively, while at KRC, the thresholds were 75 points m^-2^ for *zmax* and 50 points m^-2^ for *zmode*. Reliability ratio of *zmax*, height skewness, height quantiles, and the coefficient of variation of mean leaf area density (LAD) also varied dependent on point density at AINS. At both sites, metrics related LAD varied significantly (p < 0.001) at all but the highest point densities, and the reliability ratio for *zmode*, kurtosis of height distribution and mean horizontal coefficient of variation of LAD varied across point densities without any clear pattern. Point density mainly affected correlations between LAD-derived structural metrics and other metrics (e.g., as point density increased, Shannon diversity of LAD changed from being positively to negatively correlated to *zmax*). This study demonstrates how point density differentially affects lidar-derived forest structure parameters in diverse forest types. Scientists must understand these effects to interpret and compare forest structure attributes derived from different lidars.

## 1. Introduction

Three-dimensional forest structure plays an important role in processes such as biomass accumulation and fire spread and provides wildlife with habitat and foraging spaces. Trees accumulate biomass by converting carbon into living matter, including stems, branches, and foliage. Aboveground forest biomass can be derived through destructive or nondestructive sampling [1]. Destructive sampling requires removal and weighing of all live and dead organic matter (e.g., stems, branches, leaves, flowers, fruits, snags, litter) within a plot [1,2]. Nondestructive sampling is conducted by measuring attributes such as canopy height and area, stem diameter at breast height, and wood specific gravity and using allometric equations to relate these measurements to aboveground biomass [1,3]. How and where a fire will spread is dependent on fuel loading within a forest. Ladder fuels are related to the distance between the forest surface and live canopy fuel layers [4] and allow fires to climb up a forest canopy reaching into the mid-and upper levels, increasing fire severity [5]. The smaller the distance between the surface and live canopy, the more likely a fire is to spread into a forest canopy. As habitats become more structurally complex, the species diversity within the habitat will increase [6,7].

This pattern has been seen in multiple species including birds [8,9], primates [10], macropods [11], amphibians [12], spiders [13], and other arthropods [14]. Habitat complexity has multiple components, including maximum, minimum, and mean canopy height, vertical diversity (often referred to as entropy), and canopy cover. Unfortunately, characterizing forest structure in the field at the scales needed to estimate biomass accumulation, fuel loading, and habitat complexity is both time and cost prohibitive.

The advent of lidar (light detection and ranging) has made it possible to quickly and accurately measure forest structure at plot, landscape, and even global scales [15,16]. Lidar has been used successfully to estimate biomass in both temperate [17] and tropical forests [18,19], which has been essential for measurement reporting and verification of programs such as REDD+ [20]. Additionally, land managers have used lidar data to estimate fuel loadings [21–23], including ladder fuels [24,25], in forests, allowing them to better maintain forests through prescribed burns and respond to wildfires. Researchers have used lidar to understand how forest structure changes in response to disturbance [26], which allows forest managers to create burn regimes that maximize diversity. Lidar-derived forest structure has also been used to better understand habitat structure requirements for various species [27] including birds [28,29], bats [30,31], primates [32], spiders [33], and other arthropods [34,35]. Given the relatively low cost to area ratio of lidar collection, it is not surprising that its use for forest structure quantification has exploded in the last several decades.

Historically, researchers have used lidar to characterize forests in terms of height statistics and distributions using metrics such as maximum and mean height and the skew and kurtosis of height distribution as structural measures [36]. Within the past decade, there has been a pivot towards using metrics derived from volumetric pixels (voxels) [37]. To calculate voxel-based metrics, researchers first divide the point cloud into vertical and horizontal partitions, forming three-dimensional voxels, then calculate metrics based on the lidar points that occur within each voxel [38]. Use of voxel-based metrics may improve estimates of lidar-derived forest structure parameters such as basal area and stem density [38], which could improve biomass [39] and fuel loading [22] estimates and increase our understanding of habitat requirements for different species [29].

Accuracy of lidar-derived forest measurements may be dependent on collection parameters, including point density (i.e., the number of points that return to the lidar sensor over a given area). Several previous studies have measured effects of point density on parameters derived directly from lidar (e.g., maximum height, height quantiles [38,40–42]) or modeled from lidar-derived parameters (e.g., biomass, Lorey’s height [43–45]). These studies have generally focused on parameters measured from the full point cloud [41,43–47], though some recent studies have also investigated the effects of point density on voxel-based parameter estimation [38,42,48,49]. While some studies show that point density has little effect on forest structure metrics at densities > 1 point m^-2^ [45,46], these relationships are site [41] and metric [42] specific. Additionally, most of these studies used lidar with low (≤ 3 points m^-2^; [40,41,45]) or moderate (>3 & ≤ 30 points m^-2^; [42–44,46,47,49]) point densities. Only [38] used high density (280 points m^-2^) lidar, collected from a helicopter, which can fly at a lower altitude than a plane, for their study. The advent of low-flying unmanned aerial systems (UAS) for lidar collection has increased the collection of high density lidar (> 100 points m^-2^) across a variety of forest types (e.g.,[50,51]). To accurately measure forest structure using lidar, one must understand the effects of lidar point density on forest structure metrics derived from the full point cloud, three-dimensional voxels, and rasterized canopy height models at multiple sites with varying forest structure.

In this study, we measure the effects of point density over a broad suite of lidar-derived forest structure parameters computed from the full point cloud, three-dimensional voxels, and canopy height models (CHM). We selected parameters that are important for biomass estimation, measuring fuel loading within forests, and describing wildlife habitat (see Table 1). We investigated the effects of point density on these lidar metrics by collecting UAS lidar with an average point density > 1000 points m^-2^ in two forest types – an open canopy pine forest and a closed canopy temperate deciduous forest to explore whether any effects of point density might be related to forest types. We hypothesize that most forest structure metrics derived from the full point cloud will not be affected by point density while metrics derived from voxels and CHM will be more sensitive to changes in point density. Additionally, we predict that effects of point density on forest structure metrics derived from voxels will be more pronounced in forests that are more structurally diverse or closed-canopy forests.

**Table 1.** Lidar collection parameters for the lidar flights over AINS and KRC.

| Site | Altitude (m) | Sensor | Spot size (mm) | Beam divergence (mrad) | Scan angle (°) | Average point density (points/m<sup>2</sup>) |
| --- | --- | --- | --- | --- | --- | --- |
| AINS | 80 | HDL-32E | 244.7 (x) | 3.0 (x) | +10.67 to - | 1070 |
|  |  |  | 121.5 (y) | 1.2 (y) | 30.67 |  |
| KRC | 80 | LiDAR PUCK(VLP-16) | 228.7 (x) | 3.0 (x) | ±15 | 1464 |
|  |  |  | 129.5 (y) | 1.5 (y) |  |  |

## 2. Methods

### 2.1 Study Sites

We conducted this study at two study sites: Assateague Island National Seashore (AINS) and Keweenaw Research Center (KRC). These sites have been part of ongoing studies into properties of various remote sensing technologies, and as such, had high-density lidar collections conducted via uncrewed aerial systems (UAS). The sites were also chosen because of their differing forest structure – a protected coastal mixed pine and shrub forest (AINS) and a disturbed, secondary growth temperate mixed hardwood forest (KRC).

Assateague Island National Seashore is located on the eastern coast of the United States in Maryland and Virginia (Figure 1). The barrier island has an open canopy forest dominated by loblolly pine (*Pinus taeda*) and interspersed with hardwood species including red maple (*Acer rubrum*) and oak species (*Quercus spp.*). Tree growth is limited to areas where there is shelter from overwash and salt winds in the center portions of the islands. Forest patches transition into shrubby communities dominated by wax-myrtle (*Myrica cerifera*) behind the coastal sand dunes where there is still protection from salt winds. AINS was established as a national seashore in September of 1965 and has been under the protection and management of the United States National Park Service since. Prior to its establishment as a national seashore, AINS had ephemeral human communities dating back to the 1850s with a push for major development in the 1950s halted by a strong storm in 1962 that destroyed structures and roads and lead to the development of the area as a national seashore [52].

**Figure 1.**
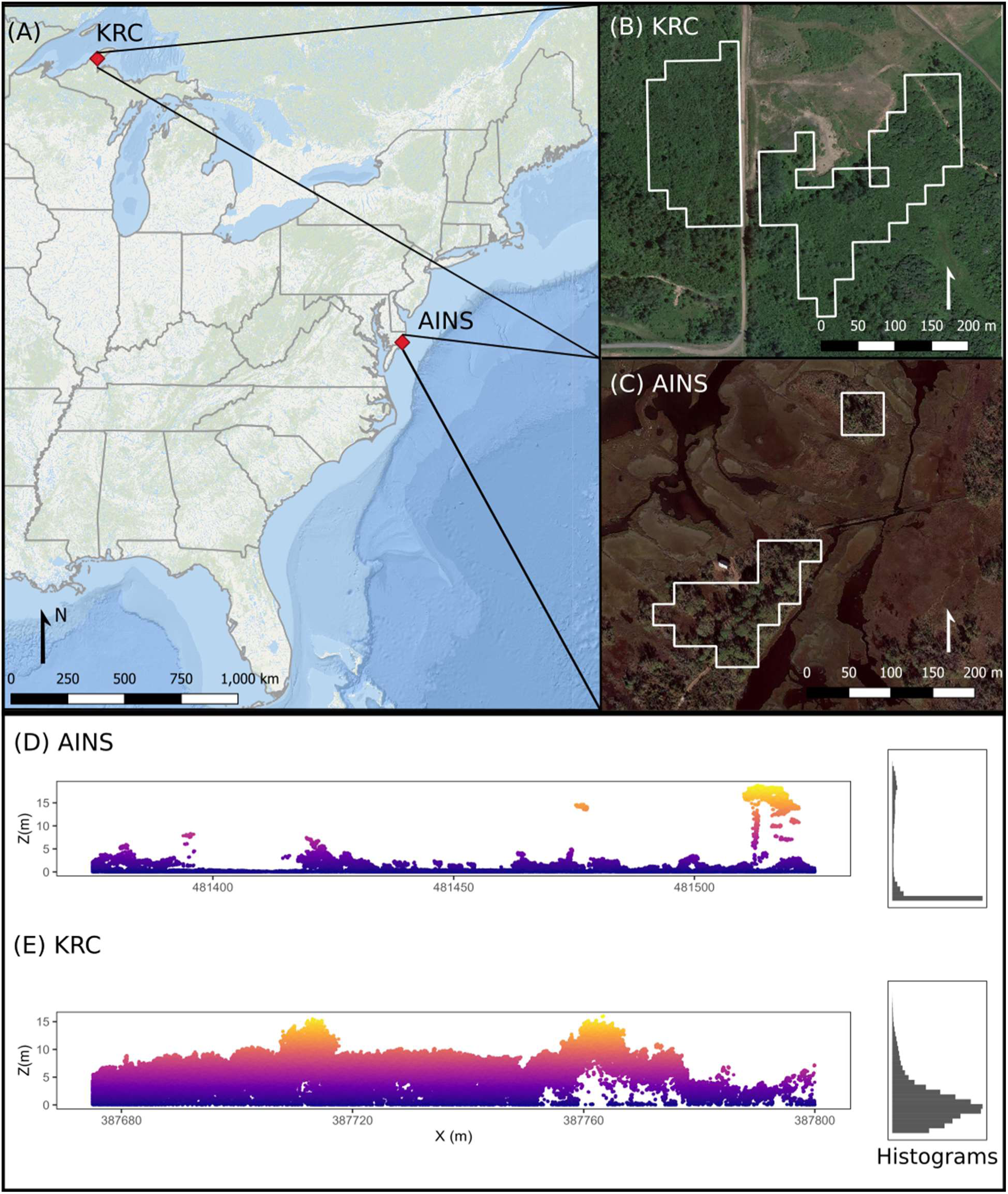
Map (A), satellite imagery with plot outlines (B, C), and lidar cross-sections with histograms of lidar return distribution across canopy layers (D,E) for study locations. AINS stands for Assateague Island National Seashore and KRC stands for Keweenaw Research Center.

Keweenaw Research Center started as a Snow, Ice, and Permafrost Research Establishment (SPIRE) and later transitioned into a vehicle research center operated by Michigan Tech. It is located in Michigan’s upper peninsula near Lake Superior (Figure 1). KRC is a disturbed secondary-growth temperate mixed hardwood forest interspersed with some pine species. The dominant species within our study area is tag alder (*Alnus serrulata*) interspersed with eastern white pine (*Pinus strobus*), jack pine (*P. banksiana*), red maple (*Acer rubrum*), and oaks (*Quercus spp.*). Some quaking aspen *(Populus tremuloides*) are present, particularly east of the north south road (Figure 1).

Lidar height return histograms show the differences in canopy structure between AINS and KRC (Figure 1D, E). AINS has a sparse open canopy. Most lidar height returns came from ground or low vegetation (Figure 1D) with a small secondary peak at approximately 15 m in height. KRC had a right long-tailed distribution with a peak in height returns at approximately 5 m and more vegetation in the midstory than AINS (Figure 1E).

### 2.2 Lidar collection parameters

We collected lidar at AINS on May 6, 2021 and at KRC on June 29, 2022. We flew a dual return Velodyne lidar (Velodyne Lidar, Inc., San Jose, CA.) mounted on a VAPOR 55 (AeroVironment, Arlington, VA) helicopter UAS. We used a Velodyne HDL-32E to collect the data over AINS and a Velodyne PUCK (VLP-16) in the KRC flight. Velodyne lidars have an oval aperture, meaning the beam is not radially symmetric, therefore we present spot size and beam divergence in both the horizontal and vertical directions. Full lidar collection parameters for the two flights are detailed in Table 1.

### 2.3 Lidar pre-processing

We processed all lidar data in R [53] using the lidR package [54]. After reading in the full point cloud for each location, we removed all points that had duplicate x, y, and z coordinates. For AINS, we subtracted the minimum z value from all points, z, to increase the minimum z value to 0 instead of a negative value (due to referencing relative to a geoid) for ease of analysis and clarity of results presentation. We then classified ground points using the cloth simulation filter (CSF) algorithm [55] in lidR’s *classify_ground* function. CSF simulates repeatedly draping a piece of cloth over an inverted point cloud. Points that intersect with the simulated cloth are classified as ground points. Because our lidar point cloud was extremely dense (AINS: 1070 points m^-2^, KRC: 1464 points m^-2^; Table 1) and the areas of interest were generally flat, we changed the default *cloth_resolution* and *rigidness* parameters to 0.01 and 2, respectively. The *cloth_resolution* parameter controls the distance between particles in the simulated cloth and is usually set to the average distance between points, and the *rigidness* parameter controls the malleability of the cloth where 1 is very soft (used for rugged terrain), 2 is medium, and 3 is hard (flat terrain) [54].

From the classified ground points, we created digital terrain models (DTM) using the triangular inverted network (TIN) algorithm. TIN is a method for generating DTM from classified ground points that works by creating a network of interconnected triangles using the classified ground points as vertices then iteratively adjusting the elevation of each vertex ultil the network satisfies a set of constraints, such as minimizing the total surface area of ensuring that drainage patterns are consistent with observed data. Though differing point densities may have had an effect on the DTM creation, a previous study showed that those effects do not create significant errors for point densities of > 0.5 points m^-2^ [46]. The terrain at both study sites also had low relief, so it was unlikely that reduced point density would have a large effect on DTM generation. Therefore, we chose to use a consistent DTM generated from the full density lidar point cloud for all plots. We then we normalized point cloud height (z) by subtracting the DTM from the point cloud. From this normalized point cloud, we subtracted outliers, defined as points where 𝑧 < 0 or 𝑧 > *z̅* + 6𝜎_z_ where *z̅* is the mean height of the point cloud and 𝜎_z_ is the standard deviation of all heights, 𝑧.

### 2.4 Plot selection

From the normalized point cloud, we created a point density raster for each site at a 1 m^2^ resolution. We overlaid a 25 m x 25 m grid over the point density raster and eliminated any plots that did not have full lidar coverage within the grid. We then extracted the point density for each plot and eliminated any plots that did not have a minimum point density of 175 points m^-2^, resulting in a total of 125 plots (27 at AINS and 98 at KRC).

### 2.5 Lidar decimation

### 2.5.1 Sensitivity Analysis

We performed a sensitivity analysis to determine the minimum number of random decimation iterations needed to minimize the variance resulting from randomly removing points from the point cloud. We randomly selected a plot from each site and decimated the lidar point clouds to achieve point densities of 175, 150, 125, 100, 75, 50, 40, 25, 20, 15, 10, 5, 4, 3, 2, and 1 points m^-2^. We repeated the random point decimations 10, 25, 50, 100, 250, 500, 750, and 1000 times and calculated the lidar metrics for each iteration at each density. From these data, we calculated the standard error of mean (SEM) for each lidar-derived metric at each density (Eq. 1):

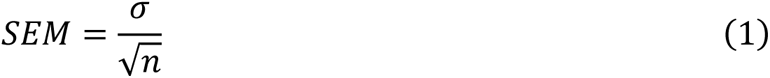

where 𝜎 is the standard deviation of the lidar-derived metric over a given number of iterations (e.g., 10, 25, 50, etc.) and 𝑛 is the number of iterations [56]. We then chose the lowest number of iterations where the SEM became asymptotic across all metrics and point densities (e.g., if SEM of maximum canopy height became asymptotic at 10 iterations for 150 points m^-2^ and 50 iterations at 1 point m^-2^, we used 50 iterations for all densities).

### 2.5.2 Random Point Decimation

After performing the sensitivity analysis and determining the number of iterations at which the SEM stabilized, we randomly decimated each point cloud for each plot over the total number of iterations. We used the *decimate_points* function from R package lidR [54] to randomly remove lidar points from each plot to achieve point densities of 175, 150, 125, 100, 75, 50, 40, 25, 20, 15, 10, 5, 4, 3, 2, and 1 points m^-2^. We repeated this procedure 250 times, as we determined from the sensitivity analysis that this was the lowest number of iteration necessary to stabilize SEM across all metrics and point densities. Figure 2 displays point clouds generated at different points densities for a plot at AINS.

**Figure 2.**
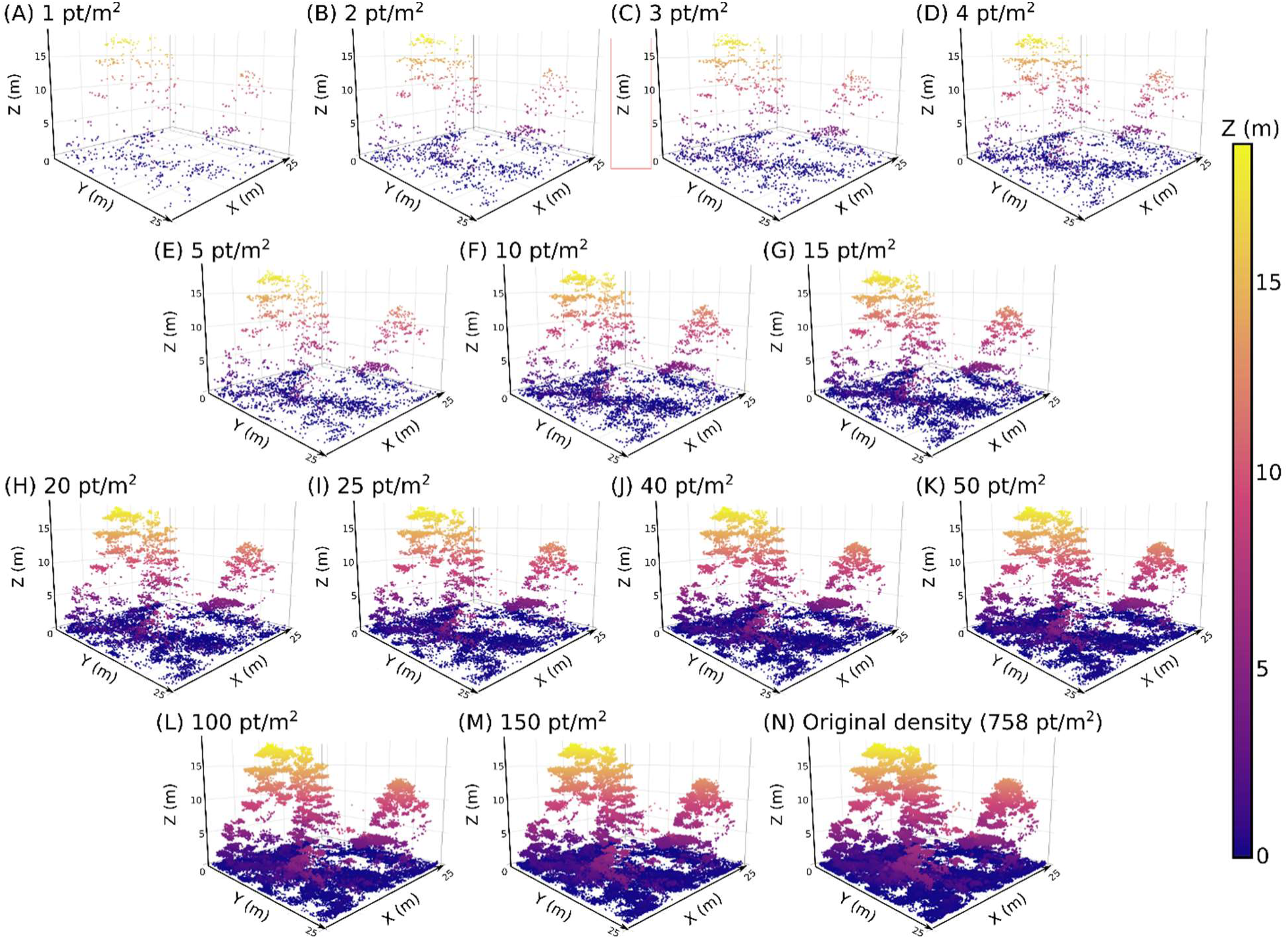
Lidar point cloud from one 25 m x 25 m buffer at AINS displayed at varying point densities: (A) 1 pt m^-2^, (B) 2 pts m^-2^, (C) 3 pts m^-2^, (D) 4 pts m^-2^, (E) 5 pts m^-2^, (F) 10 pts m^-2^, (G) 15 pts m^-2^, (H) 20 pts m^-2^, (I) 25 pts m^-2^, (J) 40 pts m^-2^, (K) 50 pts m^-2^, (L) 75 pts m^-2^, (M) 100 pts m^-2^, (N) 125 pts m^-2^, (O) 150 pts m^-2^, (P) 175 pts m^-2^, and (Q) 758 pts m^-2^, which is the original point density.

### 2.6 Variable quantification

We quantified lidar metrics for all plots at both sites by customizing functions from R packages lidR [54] and leafR [57]. We chose 18 canopy metrics calculated from the full point cloud, seven metrics calculated at the voxel level, and one metric calculated from the canopy height model (Table 2). All lidar-derived metrics were chosen based on their usage in the literature for applications in biomass estimation, biodiversity, and fire management studies (Table 2). With the exception of vertical diversity, voxel metrics were calculated for 1 m x 1 m x 1 m voxels, then averaged over the 25 m x 25 m plot. Vertical diversity was calculated at a scale of 25 m x 25 m x 1 m.

**Table 2.**
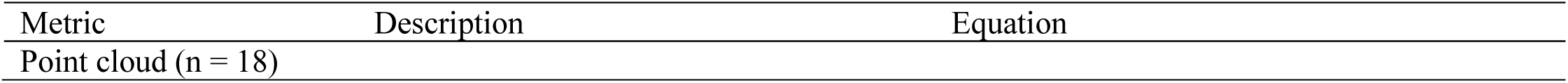
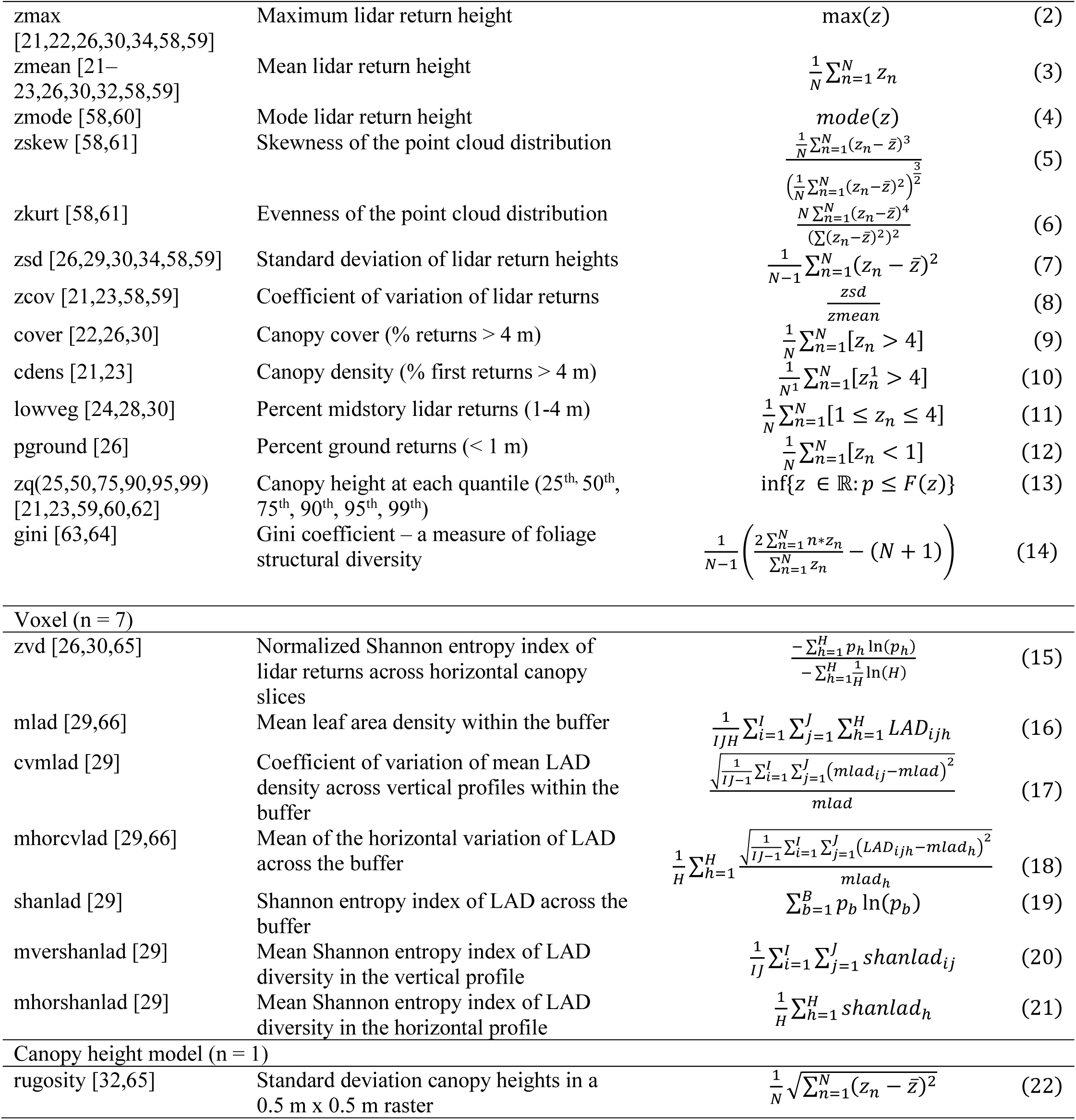
Description of lidar-derived variables and equations for calculating these variables from the lidar point cloud. For point cloud metrics, 𝑧_n_ is the height value of a lidar return, 𝑛, while 𝑁 is the total number of returns and *z̅* is the mean of all height values; 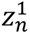 is the height value of the 𝑛_th_ first return, and 𝑁^1^ is the total number of first returns; 𝑝 is the of sampling a random 𝑧 from the point cloud that is less than or equal to a given height value of the cumulative density function, 𝐹(𝑧). For the voxel metrics, 𝑖, 𝑗, and ℎ are the voxel number, and 𝐼, 𝐽, and 𝐻 are the total number of voxels in the x-, y-, and z-directions, respectively; 𝑝_h_ is the proportion of lidar returns within the ℎ^th^ voxel, 𝑝_b_ is the proportion of LAD values within the 𝑏^th^ evenly-sized LAD bin, and 𝐵 is the total number of LAD bins. For the canopy height model (CHM) derived metric, 𝑧_n_ is the height value of the 𝑛^th^ raster cell, and 𝑁 is the total number of raster cells in the CHM.

#### 2.6.1 Cloud metrics

We derived a total of 18 canopy metrics from the point cloud for each buffer area: maximum height (*zmax*), mean height (*zmean*), mode height (*zmode*), skewness and kurtosis of the height distribution (*zskew*, *zkurt*), standard deviation of height returns (*zsd*), coefficient of variation of height returns (*zcov*), canopy cover (*cover*) and density (*cdens*), percent low vegetation (*lowveg*), percent ground (*pground*), height distribution quantiles (*zq25*, *zq50*, *zq75*, *zq90*, *zq95*, *zq99*), and the Gini coefficient (g*ini*) (Table 2; Eqs. 2-14). We set a height threshold of 4 m for canopy cover and canopy density to capture the upper canopy and considered low vegetation as returns between the upper canopy (𝑧 < 4 m) and ground (𝑧 > 1 m). Canopy height quantiles are the infimum height values, 𝑧, such that the cumulative probability, 𝑝, is less than or equal to the probability of interest (e.g., 0.25, 0.50, etc.) and are derived from the cumulative density function of lidar return heights (Eq. 13). The Gini coefficient is a measure of foliage structural diversity [67], calculated with Eq. 14, where the 𝑁 lidar returns in the point cloud are sorted from lowest to highest (i.e, 𝑧_1_ = min(𝑧) and 𝑧_N_ = max(𝑧)).

#### 2.6.2 Voxel metrics

We derived seven metrics from the voxelated point cloud: vertical diversity (*zvd*), mean leaf area density (*mlad*), coefficient of variation of mean leaf area density in the vertical profile (*cvmlad*), mean coefficient of variation of horizontal leaf area density (*mhorcvlad*), Shannon entropy index of leaf area density (*shanlad*), mean Shannon entropy index of leaf area density in the vertical profile (*mvershanlad*), and mean Shannon entropy index of leaf area density in the horizontal profile (*mhorshanlad*) (Table 2; Eqs. 15-21). For all voxel metrics, 𝑖 refers to the voxel in the x direction, 𝑗 refers to the voxel number in the y direction, and ℎ is the voxel number in the z direction. 𝐼 is the total number of voxels in the x direction, 𝐽 is the total number of voxels in the y direction, and 𝐻 is the total number of voxels in the z direction. To calculate vertical diversity, we subdivided the point cloud into 1 m horizontal bins (canopy slices) so that we had voxels of dimensions 25 m x 25 m x 1 m. We the used the normalized vertical complexity index [68] to measure the spread and evenness of lidar returns within the canopy slices (Eq. 15).

The other six voxel metrics were calculated by first deriving the leaf area density (LAD) from 1 m x 1 m x 1 m voxels. LAD is a measure of the vertical and horizontal leaf distribution within a forest canopy. To calculate LAD (Eq. 23), we used a method based on [69] where we counted the number of lidar pulses entering (𝑆_e_) and exiting (𝑆_t_) each voxel. For each voxel,

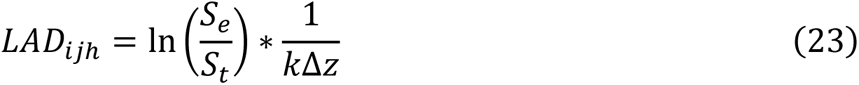

where 𝑘 is an extinction coefficient and 𝑧 is the voxel height. In a forest canopy, the Beer-Lambert extinction coefficient, 𝑘, corrects for non-random foliage distribution and orientation, as well as leaf thickness, all of which affect light dispersion. We set 𝑘 = 1 based on results from [70] who found an extinction coefficient value of 𝑘 = 1.01 was most accurate for estimating LAD at a 1 m^3^ resolution using an airborne lidar sensor (Goddard’s LiDAR, Hyperspectral, and Thermal Imager, G-LiHT) with an average point density of 15.86 pts m^-2^. While the choice of extinction coefficient for our study sites may provide inaccurate LAD estimates, the goal of this study was to compare how metrics derived from lidar with differing point densities varied so we did not further calibrate the extinction coefficient.

We used the methods described in [29] to derive *mlad*, *cvmlad*, *mhorcvlad*, *shanlad*, *mvershanlad*, and *mhorshanlad*. We chose these metrics because they were all ranked within the top seven most important forest structure variables related to bird species richness in Carrasco et al.’s models. Mean LAD (*mlad*) is the mean leaf area density of all voxels within the plot (Table 2, Eq. 16). We also calculated vertical and horizontal variation of LAD within the plot (Figure 3). For each grid cell, ij, in the plot, we calculated mean LAD across all heights, h (*mlad*ij, Figure 3a; Appendix A, Eq. A1). We then calculated the coefficient of variation of *mlad*ij (*cvmlad*) for all grid cells across the buffer (Table 2, Eq. 17; Figure 3b). To measure horizontal variation of LAD in the plot, we first calculated the mean LAD across all grid cells, ij, for a single height, h (*mlad*h, Figure 3c; Appendix A, Eq. A2). We then calculated the mean coefficient of variation of horizontal LAD (*mhorcvlad*; Table 2, Eq. 18; Figure 3d).

**Figure 3.**
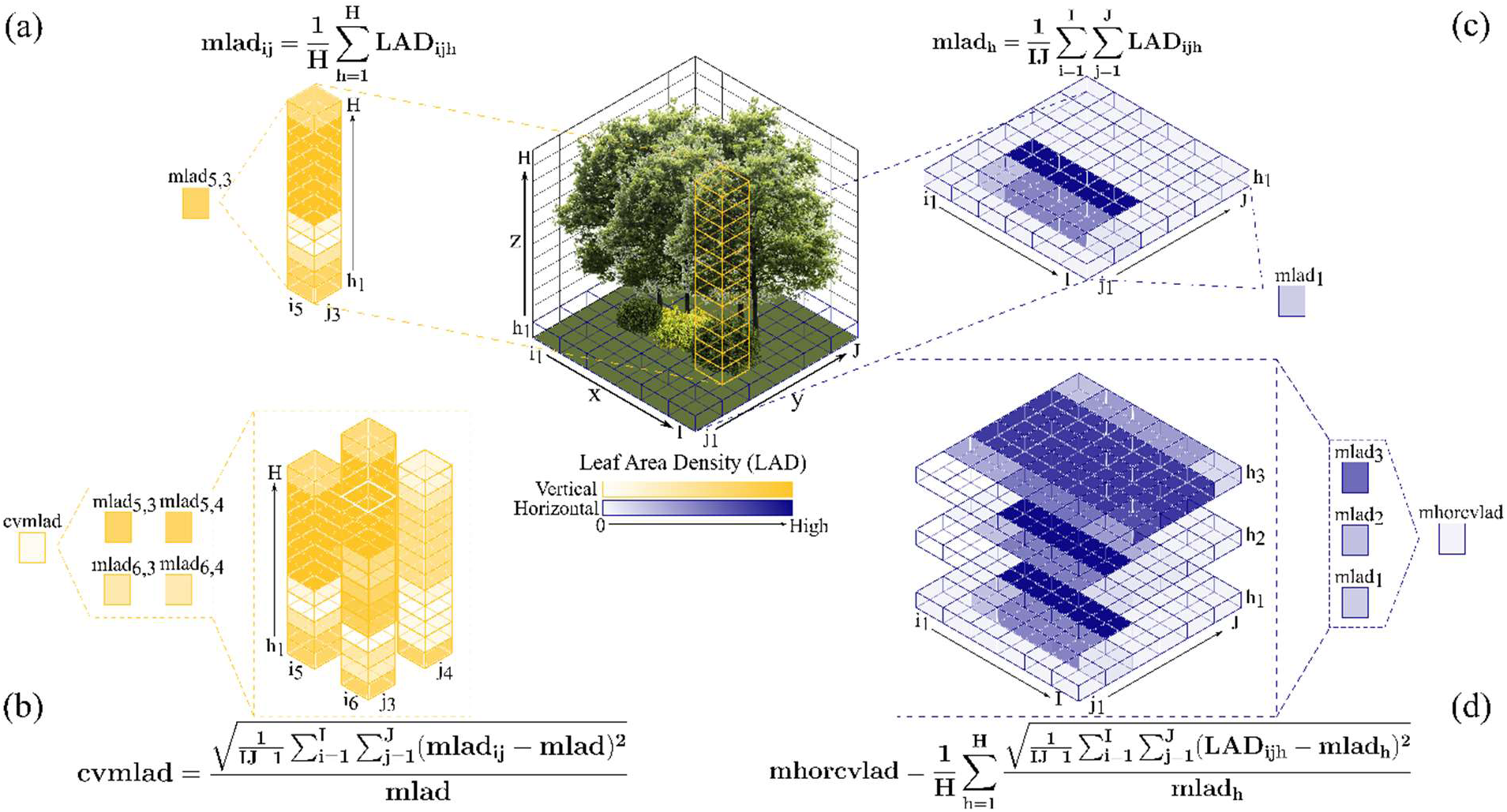
Schematic of vertical and horizontal variation of leaf area density (LAD) within a forested buffer: (a) mean LAD for a single grid cell (*mlad*ij) at heights h=1 to H; (b) coefficient of variation of *mlad*ij across four grid cells (*cvmlad*); (c) mean LAD for all grid cells at a single height (*mlad*h); (d) mean coefficient of variation of *mlad*h across three heights (*mhorcvlad*).

We calculated three different LAD diversity measures for each buffer: overall LAD diversity for the plot (*shanlad*; Table 2, Eq. 19), mean LAD diversity for the vertical profile (*mvershanlad*; Table 2, Eq. 20), and mean LAD diversity for the horizontal profile (*mhorshanlad*; Table 2; Eq. 21). To calculate the proportion of LAD values within even-sized bins, we first divided the range of LAD for the entire buffer by 50 [29]. For each voxel, 𝑖𝑗ℎ, we assigned 𝐿𝐴𝐷_ijh_ to its corresponding bin. We then calculated the proportion of voxels within a bin, 𝑏, dividing number of voxels within that bin by the total number of voxels across all bins, 𝐵. Equations for calculating *shanlad*ij and *shanlad*h are included in Appendix A (Eqs. A3 and A4, respectively).

#### 2.6.3 Canopy height metrics

We derived a single metric, *rugosity*, from the canopy height model. We created the canopy height model 0.5 m x 0.5 m resolution with the *rasterize_canopy* function in lidR. We then calculated *rugosity* by taking the standard deviation of canopy heights across the raster grid (Table 2, Eq. 22).

### 2.7 Statistical Analyses

We performed multiple analyses (described further below) on the variables derived from lidar point clouds of varying point density to explore how changes in point density affected interpretability of metrics from different perspectives. To determine whether metrics were significantly different when derived from higher versus lower density lidar, we performed analysis of variance (ANOVA) and Tukey’s post hoc tests. We used the reliability ratio [45] to quantify replication effects for different point densities. We also conducted correlation analyses to understand how metrics derived from differing point densities related to each other and whether these correlations changed as point densities changed. Analyses were performed separately for each site.

#### 2.7.1 Analysis of Variance

To assess whether there were significant differences in lidar metrics derived from different point densities, we performed an ANOVA. We first checked for homoscedasticity of variance for all metrics with Bartlett’s test [71] using the *bartlett.test* function in R. For those variables that met the criteria of homoscedasticity of variance, we performed a standard ANOVA. For those metrics that violated the assumption of homoscedastic variance, we used a white-adjusted ANOVA as this test uses a coefficient covariance matrix that is adjusted for heteroscedasticity of variance [72]. We performed these tests with the *anova* function in the car package in R [73]. For all metrics that differed significantly based lidar point density, we performed Tukey’s post hoc test with the *TukeyHSD* function in R to make pairwise comparisons between point densities [74].

#### 2.7.2 Reliability Ratio

Metrics derived from lidar are considered to be random variables due to the stochastic processes in which photons return to a lidar receiver, which are related to vegetation and leaf shape, fight altitude, flight speed, and other parameters [45]. To measure the impact of the random nature of lidar returns on lidar metric quantification, [45] used the reliability ratio (Eq. 24), which relates the variance of within plot variances across all plots to the among plot variance and is expressed as

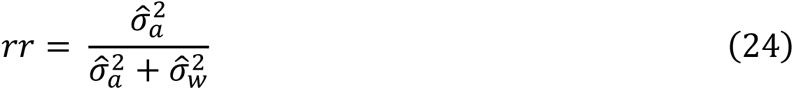

Where 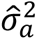 is the among plot variance and 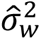 is the variance of within plot variances. We calculated 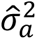 for all point density decimation iterations across all plots at each site (Appendix A, Eq. A5). To calculate 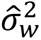, we first calculated the variance of each lidar metric derived from the 250 decimation iterations for each plot (Appendix A; Eq. A6). Then, we calculated the variance of those variances across all plots at each site (Appendix A; Eq. A7). We calculated the reliability ratio in R using the *summaryBy* function from the *doBy* package [75]. Because

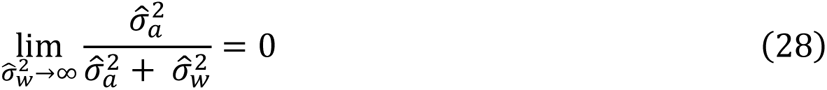

and

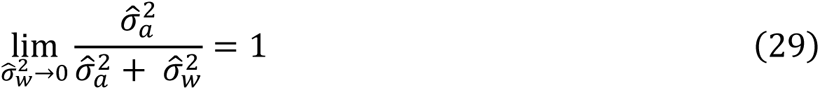

a metric is considered more reliable when the variance of within plot variances is lower than the among plot variance.

#### 2.7.3 Correlation Analysis

Using the *rcorr* function in the *Hmisc* package [36], we calculated Pearson’s r to determine whether changes in point density affected correlation between lidar-derived metrics. At each site, we calculated the correlation between metric pairs for each point density. We then examined whether there were any changes in correlations between metrics as point density changed.

## 3. Results

### 3.1 Sensitivity Analysis

The standard error of mean (SEM) was generally low (SEM < 0.1) for most metrics at both AINS and KRC (Figure 4). At AINS, five metrics had a SEM higher than 0.1 for at least one point density: *zmax*, *pground*, *zq90*, *zq99*, and *cvmlad* (Figure 4A). At KRC, only three metrics had a SEM higher than 0.1 for at least one point density: *zmode*, *pground*, and *mhorcvlad* (Figure 4B). SEM was asymptotic for most metrics and point densities at approximately 100 random decimation iterations at both sites. However, when point density was 1 point m^-2^, 18 of the 26 metrics at AINS and 20 of the 26 metrics at KRC did not reach an asymptote until 250 random decimation iterations (Figure 4). Therefore, we chose to repeat random point decimations 250 times to ensure that SEM was minimized for all metrics and all point densities.

**Figure 4.**
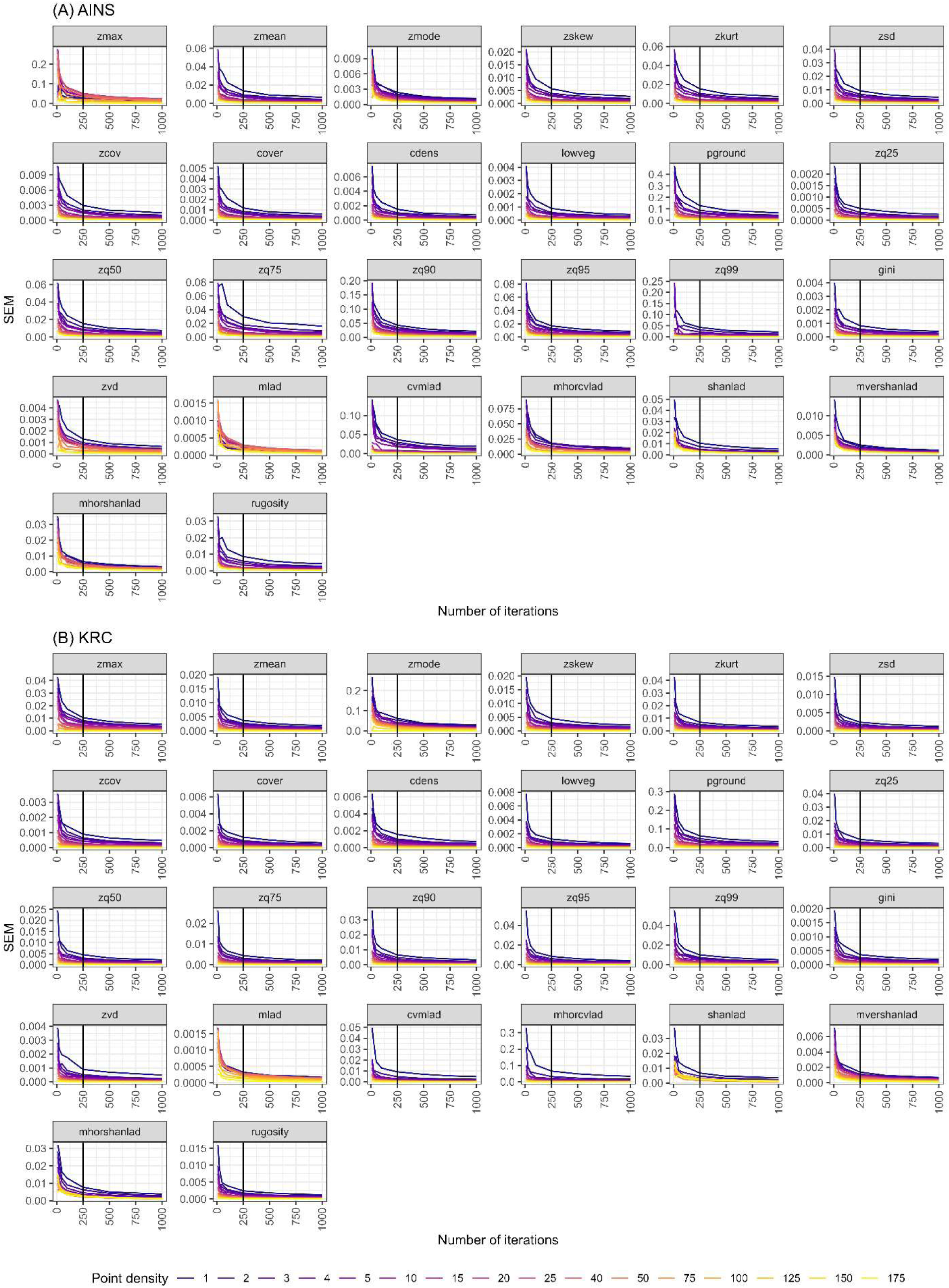
Standard error of mean (SEM) for each metric across 1-1000 iterations for each point density at (A) AINS and (B) KRC.

### 3.2 ANOVA

*3.2.1 Cloud metrics*

Of the 18 metrics derived from the full lidar point cloud, only *zmax* and *zmode* were significantly different at different point densities (Appendix B, Table B1-1). Mean *zmax* increased with increasing point density. There was a 0.78 m (AINS) and 1.36 m (KRC) difference in mean *zmax* calculated from lidar clouds with point densities of 1 point m^-2^ and 175 points m^-2^ (Appendix B, Figure B1-1). The standard deviation of *zmax* was higher at AINS than at KRC. At AINS, standard deviation of *zmax* was 5.54 m at 1 point m^-2^ and 4.96 m at 175 points m^-2^. At KRC, standard deviations of *zmax* were 3.74 m and 3.64 m at 1 point m^-2^ and 175 points m^-2^, respective. Mean *zmode*, decreased as point density increased. The difference in mean *zmode* between the highest (175 points m^-2^) and lowest (1 point m^-2^) density point clouds was greater at KRC (0.33 m) than AINS (0.07; Figure B2). Variance of *zmax* (Appendix B, Figure B1) and *zmode* (Appendix B, Figure B1-2) also decreased with increasing point density.

At AINS, there were no significant differences in *zmax* at point densities of at least 10 points m^2^ while at KRC, the threshold was 75 points m^-2^ (Figure 5). For *zmode*, significant differences between point densities were evident for fewer than 25 points m^-2^ at AINS and 50 points m^-2^ at KRC (Figure 5). However, at KRC, *zmode* was not significantly different when calculated from lidar with point densities of 1 to 5 points m^-2^.

**Figure 5.**
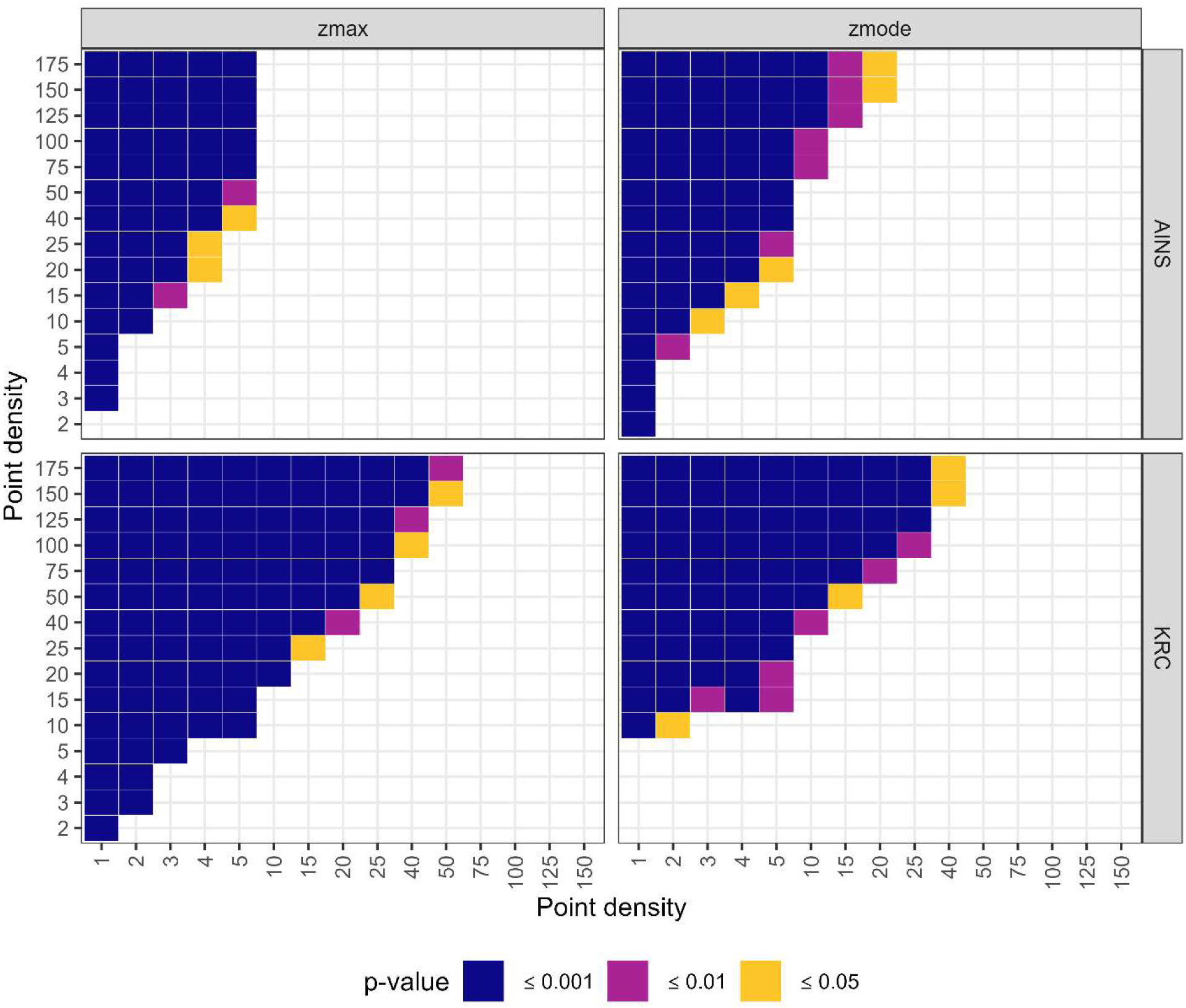
Results from Tukey’s HSD for metrics calculated from the full point cloud. Colors indicate significance level. Missing values indicate metrics did not significantly differ between the paired point densities.

#### 3.2.2 Voxel metrics

All voxel metrics except *zvd* differed significantly based on point density at both sites (Table 4). Vertical diversity (*zvd*) was only significantly different at KRC for point densities of up to 100 points m^-2^ for all buffer sizes (Figure 6). Mean leaf area density (*mlad*) was significantly different for point densities up to 150 points m^-2^ at AINS and all point densities at KRC (Figure 6). The coefficient of variation of *mlad* (*cvmlad*) differed significantly for point densities less than 125 points m^-2^ at AINS and 75 points m^-2^ at KRC (Figure 6). At AINS, the mean horizontal coefficient of LAD varied significantly for point densities less than 75 points m^-2^ (Figure 6). Differences related to lidar point density for *mhorcvlad* were more dynamic at KRC. There, *mhorcvlad* varied significantly for point densities less than 125 points m^-2^ (Figure 6), but when for point densities greater than 3 points m^-2^, there was only a significant difference in *mhorcvlad* when compared to 150 points m^-2^ or more (Figure 6).

**Figure 6.**
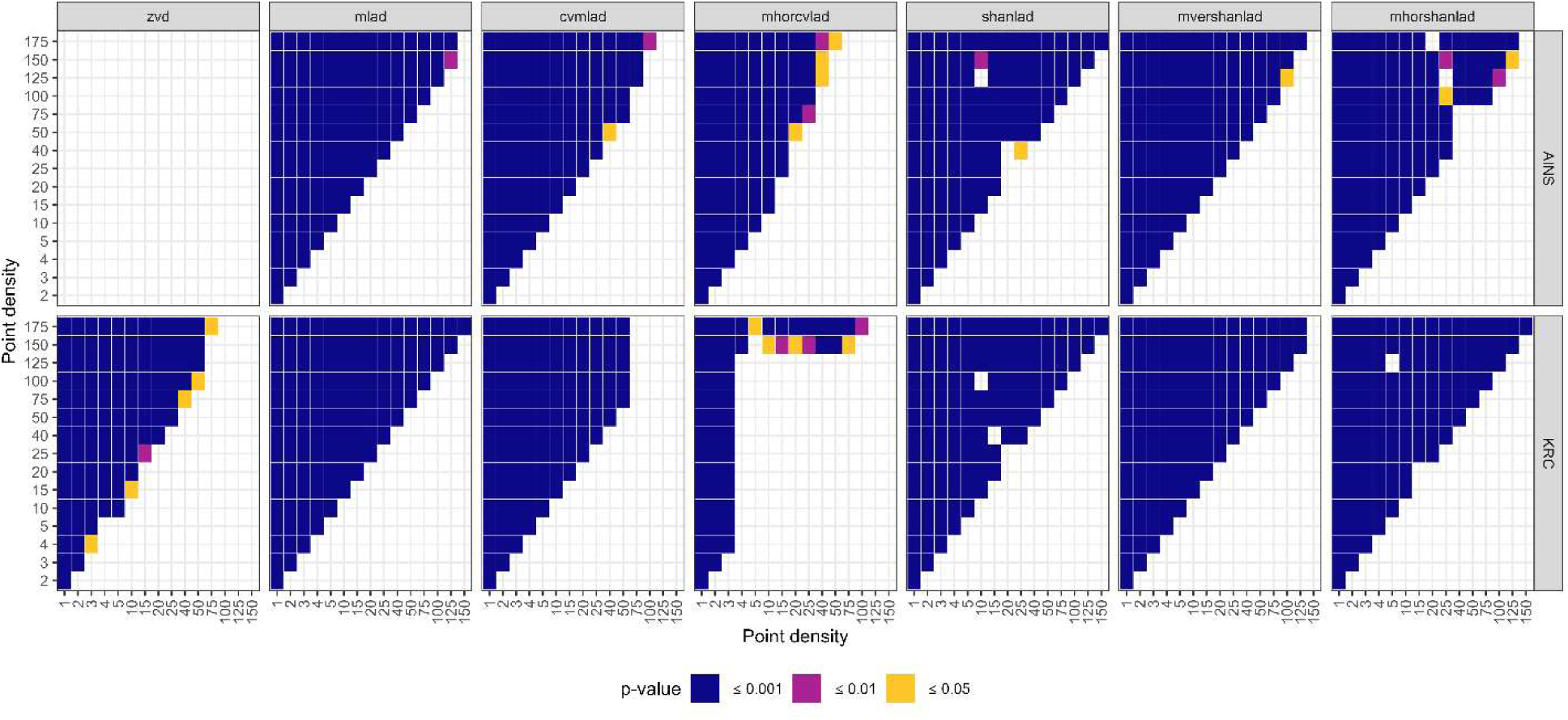
Results from Tukey’s HSD for metrics calculated from voxelized lidar. Colors indicate significance level. Missing values indicate metrics did not significantly differ between the paired point densities.

Shannon diversity of LAD (*shanlad*) differed significantly for most paired point density comparisons at both AINS and KRC. However, at AINS, *shanlad* was not significantly different when calculated from lidar with point densities of 10 points m^-2^ and 125 points m^-2^ and 20 points m^-2^ compared to 25 or 40 points m^-2^ (Figure 6). At KRC, there were no differences in *shanlad* when it was calculated from lidar point clouds with densities of 15 points m^-2^ and 40 or 100 points m^-2^ (Figure 6). At both sites, *mvershanlad* varied significantly for point densities lower than 150 points m^-2^ (Figure 6). Differences in *mhorshanlad* were dynamic at both AINS and KRC. At AINS, *mhorshanlad* varied significantly for point densities lower 150 points m^-2^, excluding 20 points m^-2^ compared to 175 points m^-2^ and 25 points m^-2^ when compared to 125 points m^-2^ (Figure 6). At KRC, there were significant differences in *mhorshanlad* for almost all point densities except between 5 and 125 points m^-2^ and between 15 and 20 points m^-2^ (Figure 6).

#### 3.2.3 Canopy height model metrics

*Rugosity* varied significantly based on point density at KRC but not at AINS. There was no significant difference in *rugosity* when derived from lidar with point densities of 1-10 points m^-2^ or 75-175 points m^-2^ (Figure 7). *Rugosity* calculated from lidar with point densities between 1-5 points m^-2^ significantly differed from that calculated from lidar with point densities of 15 points m^-2^ or more (Figure 7).

**Figure 7.**
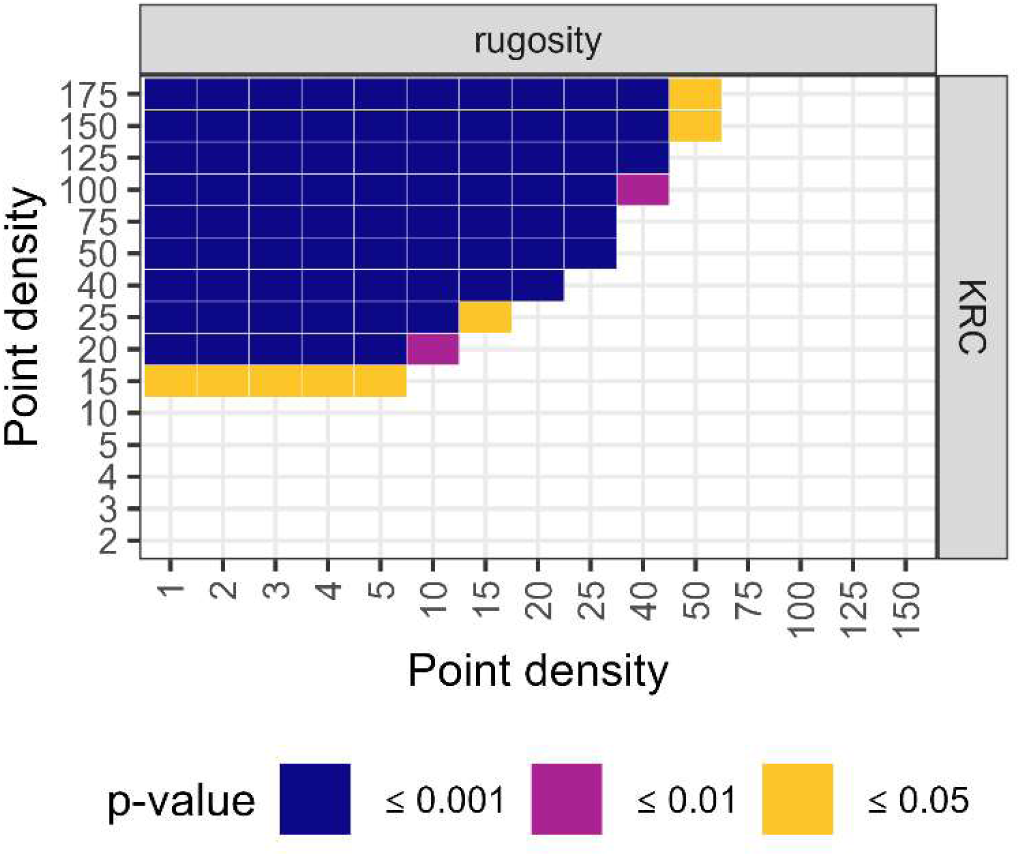
Results from Tukey’s HSD for metrics calculated from the canopy height model. Colors indicate significance level. Missing values indicate metrics did not significantly differ between the paired point densities.

### 3.3 Reliability Ratio

#### 3.3.1 Cloud metrics

Of the 18 metrics derived from the full point cloud, 10 had a reliability ratio ≥ 0.90 across all treatments , indicating the metrics were reliable independent of lidar point density and study site [45,46]. These metrics were *zmean*, *zsd*, *zcov*, *cover*, *cdens*, *lowveg*, *pground*, *zq95*, *zq99*, and *gini*. Seven of these 10 metrics (*zmean*, *zsd*, *zcov*, *cover*, *cdens*, *lowveg*, and *gini*) had a reliability ratio of one for all point densities at both sites. Reliability ratio for *pground* only deviated from one at 1 point m^-2^ at both AINS and KRC, but this deviation was small (< 0.025; Figure 8).

**Figure 8.**
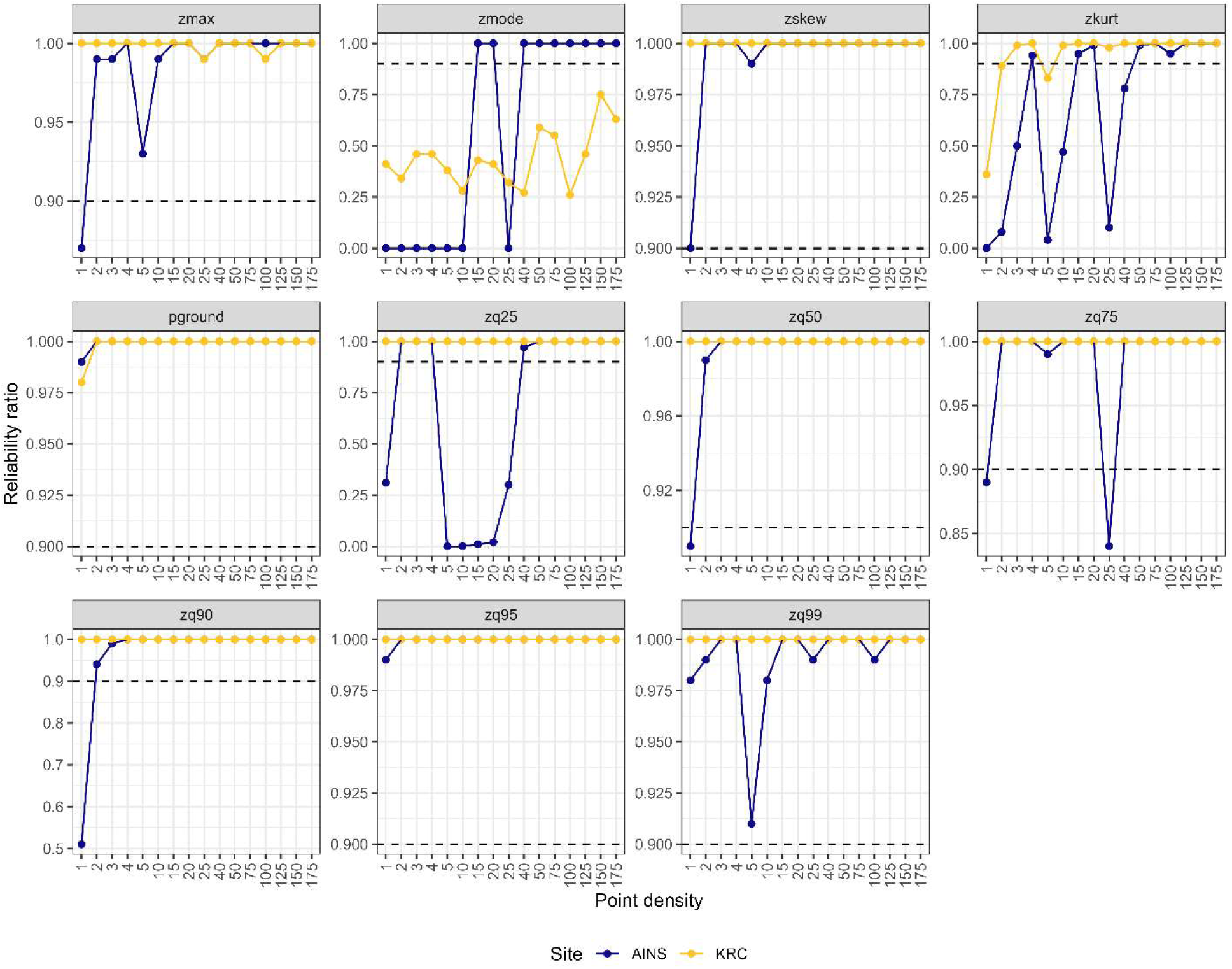
Reliability of metrics derived from the full lidar point cloud. Colors represent the two sites. The dashed horizontal line is realiability ratio = 0.90 (the cutoff for a metric to be considered reliable [45,46]).

Generally, the reliability ratio varied more at AINS than at KRC. For instance, while *zq95* and *zq99* were both considered reliable at both sites (rr > 0.90), at KRC, the ratio was one for both metrics at all point densities. At AINS, the reliability ratio for *zq95* was less than one at 1 point m^-2^ and the ratio for zq99 ranged from 0.91 to one without any clear pattern related to point density (Figure 8). The reliability ratio of *zmax*, *zskew*, *zq25*, zq50, *zq75*, and *zq90* was greater than 0.90 for all point densities at KRC but dropped below this threshold for at least some point densities at AINS. There, *zmax*, *zskew*, zq50, and *zq90* were reliable when calculated from lidar with point densities of at least 2 points m^-2^.

The reliability of both *zq25* and *zq75* was dynamic at AINS. For *zq25*, the reliability ratio was less than 0.90 for 1 point m^-2^ and again for point densities between 5 and 25 points m^-2^; *zq75* was similarly unreliable at 1 point m^-2^ and 25 points m^-2^ (Figure 8). Likewise, the reliability of *zkurt* varied considerably at AINS, reaching the 0.90 threshold at 4, 15, 20, and ≥ 50 points m^-2^. At KRC, the reliability ratio for *zkurt* stabilized for point densities ≥ 10 points m^-2^ (Figure 8).

Mode canopy height (*zmode*) had a reliability ratio of zero for 1-10 and 25 points m^-2^ and a reliability ratio of one for all other point densities at AINS. However, at KRC, reliability of *zmode* was below the 0.90 cutoff for all point densities (Figure 8).

#### 3.3.2 Voxel and canopy height metrics

The reliability ratio for *zvd*, *mlad*, and *mhorshanlad* was one for all point densities at both AINS and KRC. At KRC, *cvmlad*, shnlad, and *mvershanlad* also had a reliability ratio of one (Figure 9). The reliability ratio of *cvmlad* was dynamic at AINS, where it dropped below the 0.90 threshold for 25 and 100 points m^-2^ but was greater than 0.90 at all other point densities. At AINS, the reliability ratio of shanland and *mvershanlad* dipped slightly below one at 1 point m^-2^, though this change was negligible (< 0.025; Figure 9). While the reliability of *mhorcvlad* was greater than 0.90 across all point densities at AINS, there was considerable variation in the ratio that did not follow any specific pattern related to point density (Figure 9). At KRC, *mhorcvlad* became unreliable at 100 and 125 points m^-2^ (Figure 9). *Rugosity* had a reliability ratio of 1, independent of study site or point density.

**Figure 9.**
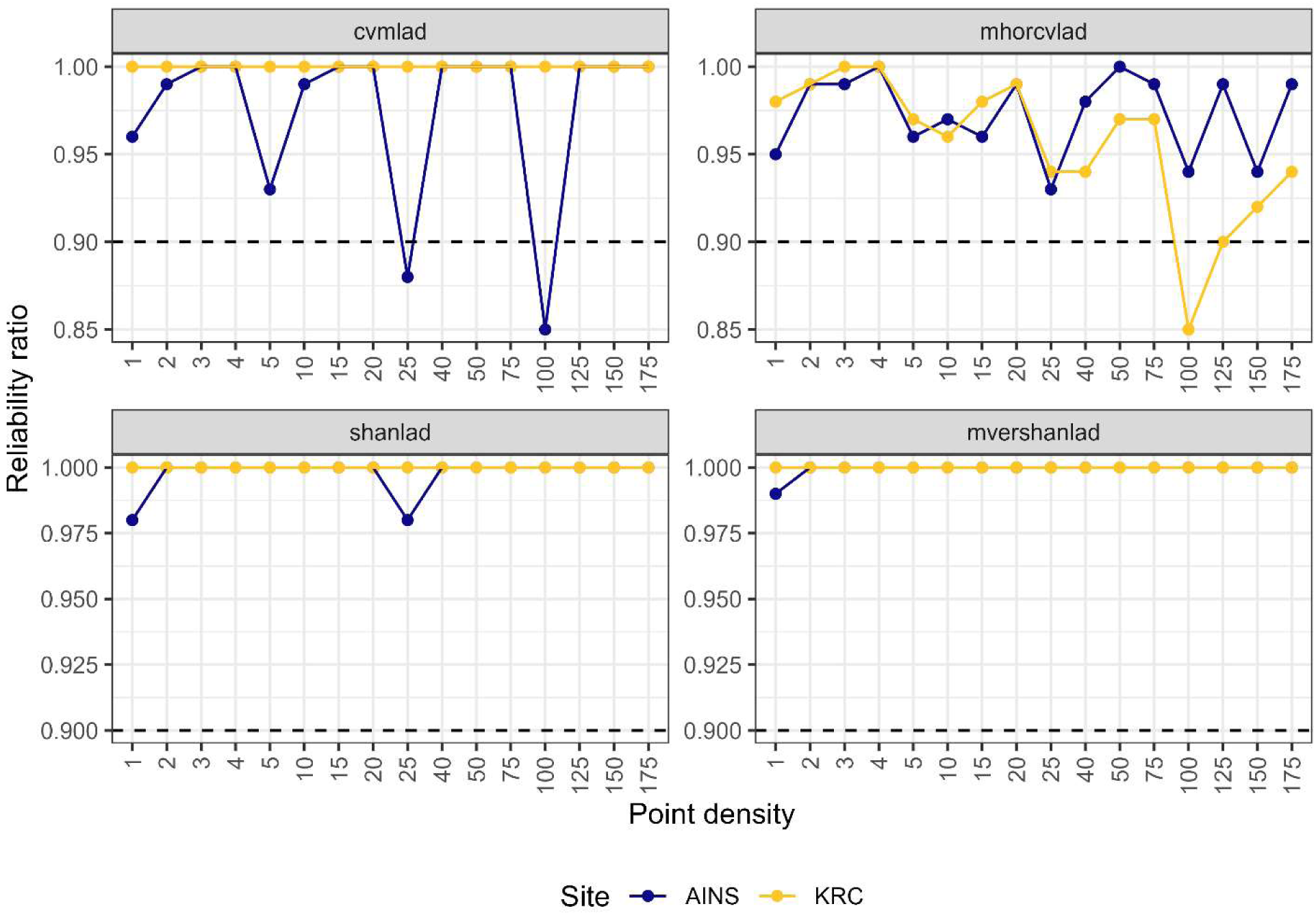
Reliability of metrics derived from the voxelized point cloud. Colors represent the two sites. The dashed horizontal line is realiability ratio = 0.90 (the cutoff for a metric to be considered reliable [45,46]).

### 3.4 Correlation Analysis

#### 3.4.1 Cloud metrics

There were no changes in direction of correlations between cloud metrics measured at different point densities (Appendix B, Figures B2-1 – B2-18). For some metrics, as point density increases, correlations with other metrics changed from strong (r ≥ 0.70) to weak (0.30 ≤ r < 0.70) or vice versa. However, in all cases, the changes in correlation strength between these metrics are quite small. The correlations between *zq75* and *zmax* and *zmode* changed from strongly positive to weakly positive when point density increased to 4 points m^-2^ (*zmax*, KRC) and 10 points m^-2^ (*zmax*, AINS; *zmode*, KRC; Appendix B, Figure B2-14). The correlation between *zkurt* and *cover* at AINS was weakly negative from 1-50 and ≥ 150 points m^-2^ and strongly negative for 75-125 points m^-2^ (Appendix B, Figure B2-5). At KRC, *zsd* and zq50 were weakly positively correlated at ≤ 3 points m^-2^ and strongly positively correlated at 4 or more points m^-2^ (Appendix B, Figure B2-6). Changes in correlation between cloud metrics and voxel metrics are described in Section 3.4.2.

#### 3.4.2 Voxel metrics

The largest variation in correlation across metrics occurs for *shanlad* at AINS (Figure 10). Correlations between *shanlad* and *zmax*, *zsd*, *cover*, *cdens*, *zq75*, *zq90*, *zq95*, *zq99*, *zvd*, *mvershanlad*, and *rugosity* all change from positive to negative at AINS as point density increases (Figure 10). For *shanlad* and *zmean* and *mhorshanlad* at AINS, correlations change from positive to negligible with increasing point density. At low point densities at AINS, *shanlad* in negatively correlated to *zskew*, *zkurt*, *zcov*, *pground*, and *cvmlad*. With increasing point density, these correlations either become negligible (*zcov*, *cvmlad*) or positive (*zskew*, *zkurt*, *pground*).

**Figure 10.**
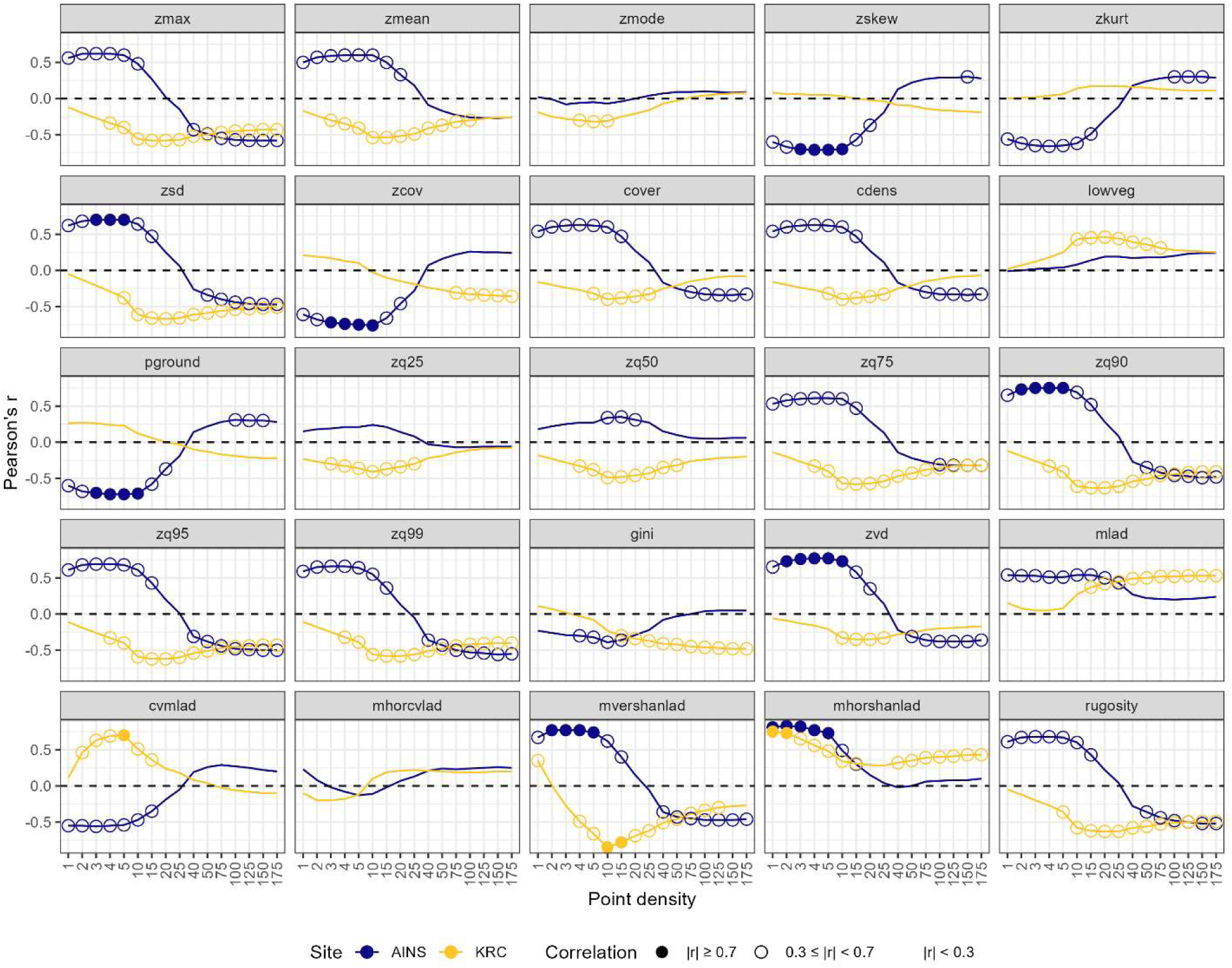
Pearson’s r for correlations between *shanlad* and the 25 other forest structure metrics calculated from lidar with differing point densities at AINS (blue) and KRC (yellow). A filled circle indicates a strong correlation (|r| ≥ 0.7), while an open circle indicates a weak correlation (0.3 ≤ |r| < 0.7) and no circle indicates a negligible correlation (|r| < 0.3).

At KRC, the only change in direction of correlation happens between *shanlad* and *mvershanlad* (weakly positive to weakly negative to negligible with increasing point density; Figure 10). However, many variables go from negligibly to weakly correlated to *shanlad* as point density increases (*zmax*, *zsd*, *zcov*, *zq75*, *zq90*, *zq95*, *zq99*, *gini*, *mlad*, *rugosity*; Figure 10). The correlations between *shanlad* and *zmean*, *zmode*, *cover*, *cdens*, *lowveg*, *zq25*, zq50, *zvd*, and *cvmlad* changes from weak to negligible at medium point densities before becoming negligible at the highest point densities (Figure 10). The breaking points for these relationships vary for each metric (Figure 10).

The correlation between *mhorshanlad* and *zq25*, which is weakly negative for 2 and 3 points m^-2^ and weakly positive for point densities ≥ 50 points m^-2^ at KRC (Figure B2-23). For other voxel-derived metrics, correlations change in strength, rather than in direction (Figures B2-19 – B2-23). For example, the correlations between *mlad* and 14 of the 25 metrics decrease in strength with increasing point density at AINS (Figure 11). These metrics are *zmax*, *zmean*, *zskew*, *zsd*, *zcov*, *cover*, *cdens*, *lowveg*, prground, *zq75*, *zq90*, *zq95*, *zq99*, and *rugosity*. At KRC, only the correlation between *mlad* and *zvd* weakens with increasing point density while the correlations between *shanlad* and *pground*, *cvmlad*, and *shanlad* all strengthen with increasing point density (Figure 11).

**Figure 11.**
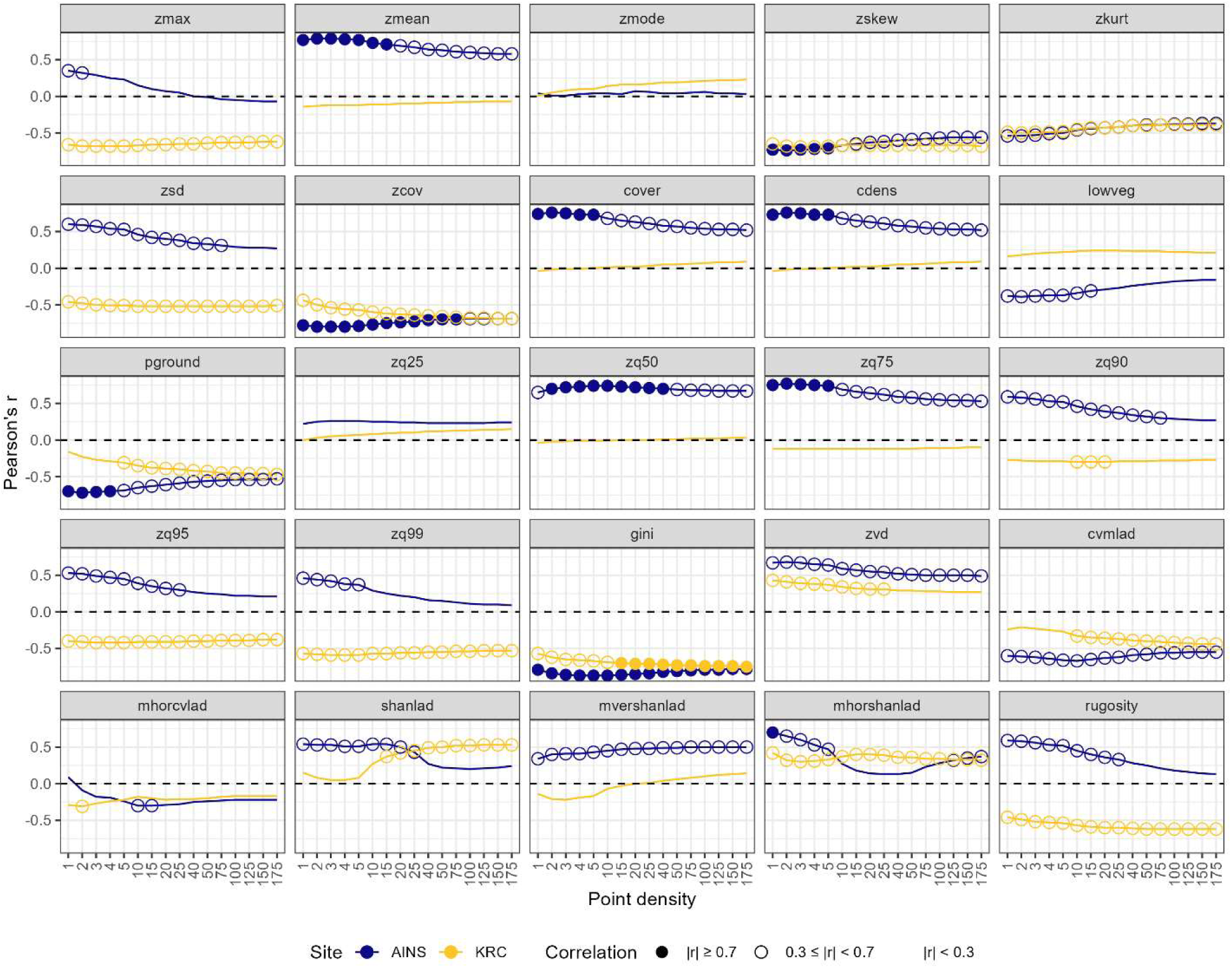
Pearson’s r for correlations between *mlad* and the 25 other forest structure metrics calculated from lidar with differing point densities at AINS (blue) and KRC (yellow). A filled circle indicates a strong correlation (|r| ≥ 0.7), while an open circle indicates a weak correlation (0.3 ≤ |r| < 0.7) and no circle indicates a negligible correlation (|r| < 0.3).

#### 3.4.3 Canopy height model metrics

Correlations between *rugosity* and *zmean*, *zmode*, and *zq75* decreased with increasing point density at KRC but not at AINS. *Rugosity* and *zmean* were strongly positively correlated (r ≥ 0.7) for point densities less than 15 points m^-2^ but weakly positively correlated (0.3 ≤ r < 0.7) for point densities of 15 points m^-2^ or higher (Figure 12). There was a similar pattern *zmode*, which was weakly correlated with *rugosity* at point densities of 10 points m^-2^ and lower but negligibly correlated to *rugosity* at 15 points m^-2^ (Figure 12). The strong positive correlation between *rugosity* and *zq75* weakens at point densities of 25 points m^-2^ and higher. At AINS, the strong negative correlation between *rugosity* and *zcov* becomes a weak negative correlation for point densities of 150 and 175 points m^-2^. However, the relationship between *rugosity* and *zcov* at KRC is negligible between 1 and 40 points m^-2^ and weak for point densities of 50 points m^-2^ or more (Figure 12).

**Figure 12.**
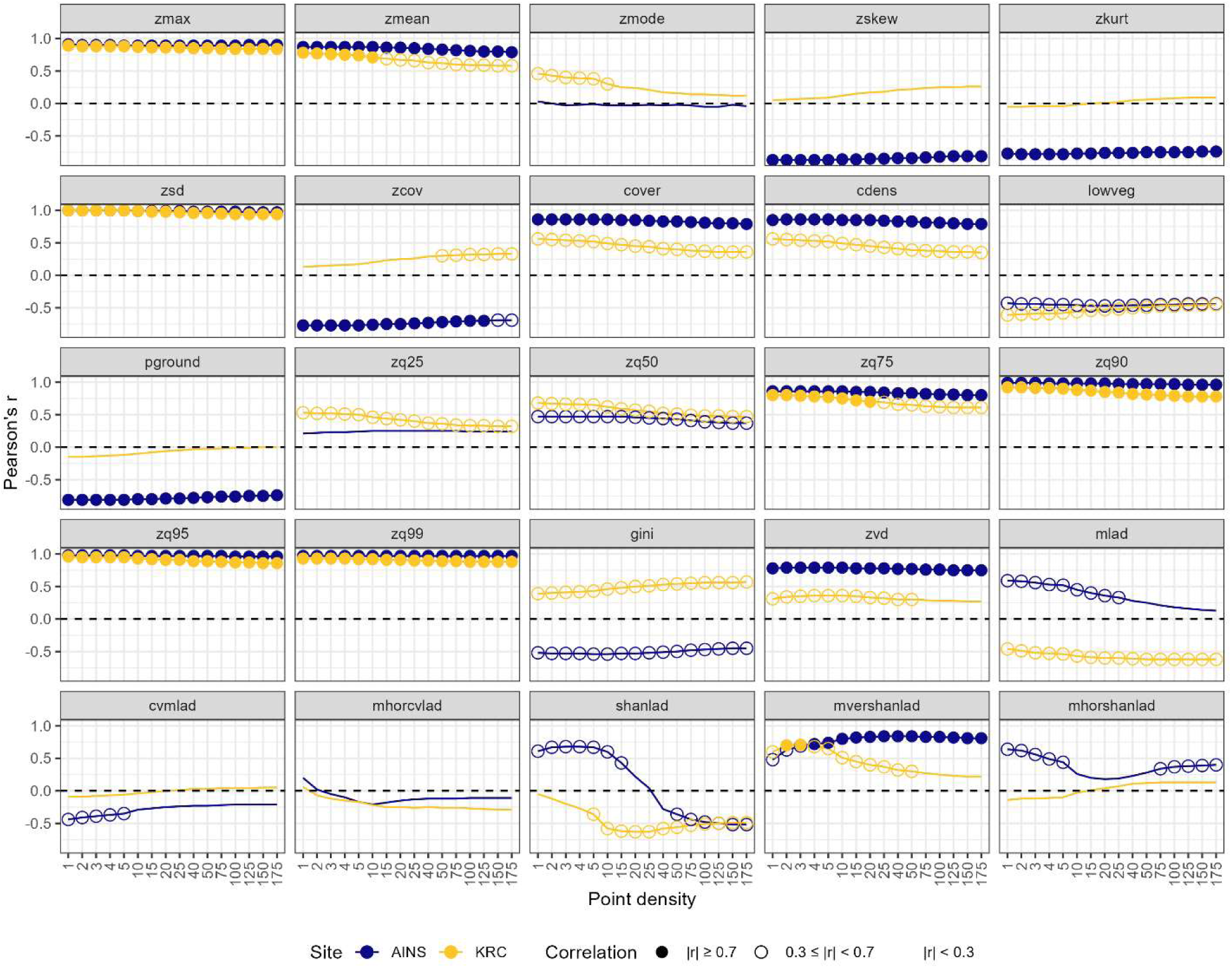
Pearson’s r for correlations between *rugosity* and the 25 other forest structure metrics calculated from lidar with differing point densities at AINS (blue) and KRC (yellow). A filled circle indicates a strong correlation (|r| ≥ 0.7), while an open circle indicates a weak correlation (0.3 ≤ |r| < 0.7) and no circle indicates a negligible correlation (|r| < 0.3).

## 4. Discussion

In this study, we examined the effects of lidar point density on forest structure parameters derived from the full lidar point cloud, three-dimensional voxels, and canopy height models.

Except for maximum and mode canopy height, lidar metrics derived from the full point cloud were not significantly different dependent on point density, which is supported by other studies [44,46,76]. However, all forest structure parameters derived from three-dimensional voxels had some level of significant difference when calculated from differing point densities. *Rugosity*, the only forest structure parameter we calculated from the canopy height model, differed significantly when calculated from different point densities at KRC but not at AINS.

Despite significant differences in forest structure metrics based on point density, most metrics had reliability ratios > 0.90 for point densities > 1 point m^-2^. The exceptions were *zmode*, *zkurt*, *zq25* (AINS), *zq75* (AINS), *cvmlad* (AINS), and *mhorcvlad* (KRC). This is similar to [45,46], who both find that reliability ratio was stable for height metrics derived from lidar with point densities of at least 2 points m^-2^ for metrics related to height. Correlations between forest structure metrics derived from the full lidar point cloud were also unchanged across point densities, while correlations between voxel-derived metrics and other forest structure metrics often either decreased in strength or changed in direction.

The differences in forest structure metrics derived from three-dimensional voxels may be due, in part, to the small voxel size we chose. As the horizontal (x,y) resolution of a voxel decreases, the number of voxels with no LAD data increases [49]. For instance, when [49] kept lidar point density constant at 10 points m^-2^ and decreased horizontal resolution from 10 m to 1 m, no-data voxels increased from 3% to 29%. However, at point densities of at least 15 points m^-2^, absolute LAD stabilized independent of horizontal voxel resolution [49]. Therefore, we would have expected differences in voxel-based measurements in our study to disappear with increasing density, even though our voxels were small.

The effects of point density on interpretation of lidar-derived forest structure metrics was often site-specific. For example, vertical diversity and rugosity were significantly different dependent on point density at KRC but not at AINS. This may be because KRC had higher vertical diversity than AINS. Our results contrast with [42], who showed that foliage height diversity, which is similar to our vertical diversity metric, stabilized at a higher point density in evergreen forest than in deciduous forests. However, [42] also compared forests that had higher average canopy cover (> 90%) than ours (24% - AINS; 42% - KRC) to each other, which may impacted these results. Despite having significant differences in more structural metrics and across a larger range of point densities, the reliability ratio for all forest structure metrics except *zmode* and *mhorcvlad* at KRC was either higher than or equivalent to the reliability ratio of forest structure metrics at AINS. Likewise, correlations between forest structure metrics did not change strength or direction at KRC while they did at AINS. Collectively, our results show that lidar-derived forest metrics may not be uniformly comparable across different forest structure types [42].

Though our lidar sensors had very similar specifications (Table 1), collecting lidar with different sensors at the two locations may have affected lidar-derived parameter estimation [76]. In general, increased footprint sizes can lead to overestimation of tree heights in mountainous terrain [77] or the opposite in terrain with less relief [47]. Both studies, however, had a difference in footprint size at least three orders of magnitude larger than the difference in footprint size between our lidars (114.4 mm^2^). Differences in scan angle can also impact estimation of forest structure variables. [76] showed that increasing a scan angle from 5° to 30° can increase height estimates of deciduous trees by 8% and coniferous trees by 19%. A larger range of scan angles at AINS compared to KRC may have led to higher estimates of maximum height at AINS. A follow-on analysis using these data sets can further explore this effect by eliminating lidar returns from more oblique angles.

Additionally, AINS had only approximately one third the number of plots that KRC had, which may help explain the greater variability in reliability ratio at AINS when compared to KRC. In a simulation study, [78] demonstrated that when fewer plots are used to model variables such as basal area and Lorey’s height, the probability of the 95% confidence interval capturing the true population mean decreases. However, despite having far fewer plots than KRC, the reliability ratio at AINS was ≥ 0.90 for most metrics when point densities increased to 2 or more points m^-2^. The lower reliability ratio at 1 point m^-2^ is likely related more to the forest structure at AINS (tall canopy trees that are widely spaced) than to the number of samples used, especially given that only eight of the 26 metrics had a reliability ratio less than 0.90 at any point density.

Our choice to use a single DTM derived from the full point density lidar from each may have resulted in under-detection of effects of point density on lidar-derived canopy metrics. For instance, [43] showed that in the complex terrain of Brazil’s Atlantic forest, decreasing point density from 20 points m^-2^ to 1 point m^-2^ resulted in an average underestimation of mean canopy height derived from the CHM of 5.26 m in a one hectare plot. Our study does not show an impact of reduced point density to estimate of mean canopy height, which we derived from the point cloud, not from the CHM. Maximum canopy height in our study decreased with decreasing point density in both closed and open canopy forests. It is possible that if we had used the thinned lidar point cloud to generate DTMs, the observed differences in maximum canopy height would have been even greater, particularly in the closed canopy forest at KRC where fewer points would penetrate the canopy to reach the ground. However, unlike the study sites in [43], which occurred across an elevational gradient, our study sites had low relief, and therefore differences in canopy height due to ground estimation would likely have a much lower impact in our study. Additionally, [46], did not show significant error in DTMs or canopy metrics generated from lidar point clouds with point densities of at least 1 point m^-2^, even in a topographically complex area with dense forest cover. Therefore, while our choice of DTM may have led us to underestimating effects of point density on lidar-derived forest structure variables, we expect that the effect would not have been great, particularly for variables derived directly from the point cloud.

Given the conic geometry of conifers, there should have been a greater difference in maximum canopy height calculated from lower versus higher point densities at AINS than at KRC [47]. However, we saw the opposite in this study, where differences in maximum canopy height at KRC were almost twice as large as at AINS. This is particularly unexpected because conifers at AINS were spaced far apart, meaning that as point density decreased, there would be a lower probability of sampling the true maximum height within any given plot. The effects of point density on estimation of maximum canopy height also stopped being significant at lower point densities at AINS than at KRC. One reason for this may be related to the larger range of scan angle at AINS compared to KRC, which could lead to an overestimation in heights [76].

This might be a relic of the smaller sample size or it may be reflective of some other pattern. To further investigate this pattern, researchers should collect lidar in an area that has both deciduous and evergreen forest on a single day with the same flight conditions and lidar collection parameters.

## 5. Conclusions

Except for maximum and mode canopy height, lidar-derived forest structure parameters derived from the full lidar point cloud are generally insensitive to point density for point densities > 1 point m^-2^. While forest structure metrics do show sensitivity to point density, this does not translate into reliability of these metrics for modeling other parameters. The relationship between lidar point density and forest structure metrics is likely to affected by forest structure type. Future research directions include examining the relationships between voxel size, point density, and lidar-derived metrics for various forest structure types.

## Author Contributions

ACS: Conceptualization, data curation, formal analysis, funding acquisition, investigation, methodology, project administration, software, validation, visualization, writing – original draft, writing – review and editing; TM: investigation, writing – review and editing; AA: funding acquisition, investigation, project administration, resources, supervision, writing – review and editing; RL: investigation; MV: investigation; WB: writing – review and editing; RRL: investigation

## Acknowledgments

We would like to thank E. Hansen for his explanation of the reliability ratio and for sharing the code he and his team developed for calculating it. R. Alger from KRC supported our data collection there and aided in identifying tree species within our collection site. Additionally, H. Littman gave valuable feedback on interpretability of several of our images.

## Appendix A. Supplementary equations for lidar metric calculations

Several of the equations used for the calculation of voxelated metrics require interim calculations. We include the equations for these interim variables within this appendix. For all equations, 𝑖, 𝑗, and ℎ represent a single voxel from the total voxels 𝐼, 𝐽, and 𝐻 in the x-, y-, and z-directions, respectively. For the Shannon diversity index equations, 𝑝_bij_ is the proportion of voxels in the grid cell, 𝑖𝑗, with LAD values that fall in the 𝑏^th^ even-sized LAD, and 𝑝_b_ is the proportion of all voxels in horizontal slice, ℎ, with LAD values in the 𝑏^th^bin; 𝐵 is the total number of occupied LAD bins.

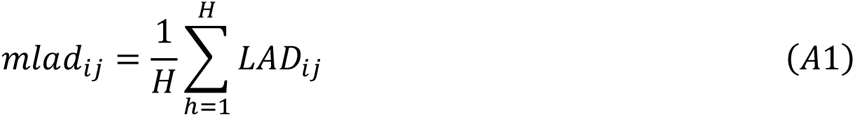

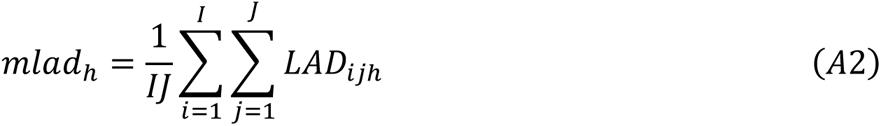

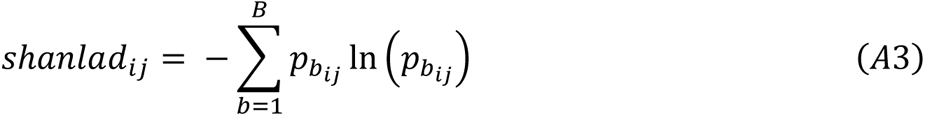

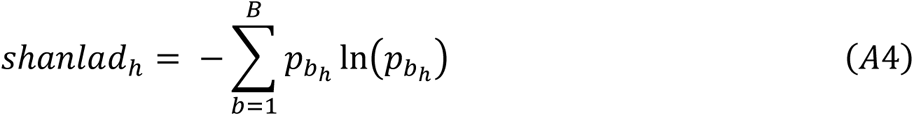

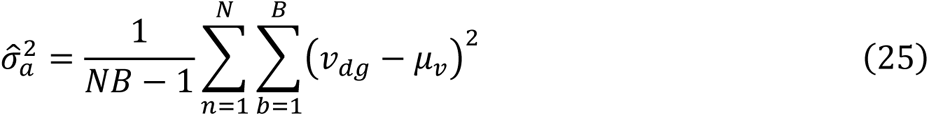

is the among plot variance and

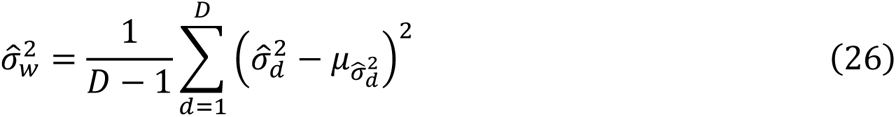

is the variance of the within plot variances across all buffers and the single buffer variance is expressed as:

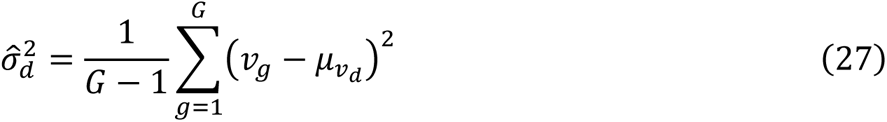

For the reliability ratio equations, 𝑑 represents a single plot from the set of all plots, 𝐷, while 𝑔 represents a single lidar decimation iteration from the set of all lidar decimation iterations, 𝐺. 𝑣 is a lidar variable and 𝜇_v_ is the mean of that variable across all plots and decimation iterations while 𝜇_vd_ is the mean for that variable for all iterations within a single plot.

## Appendix B. Full results

### B1. ANOVA results

**Table B1-1.**
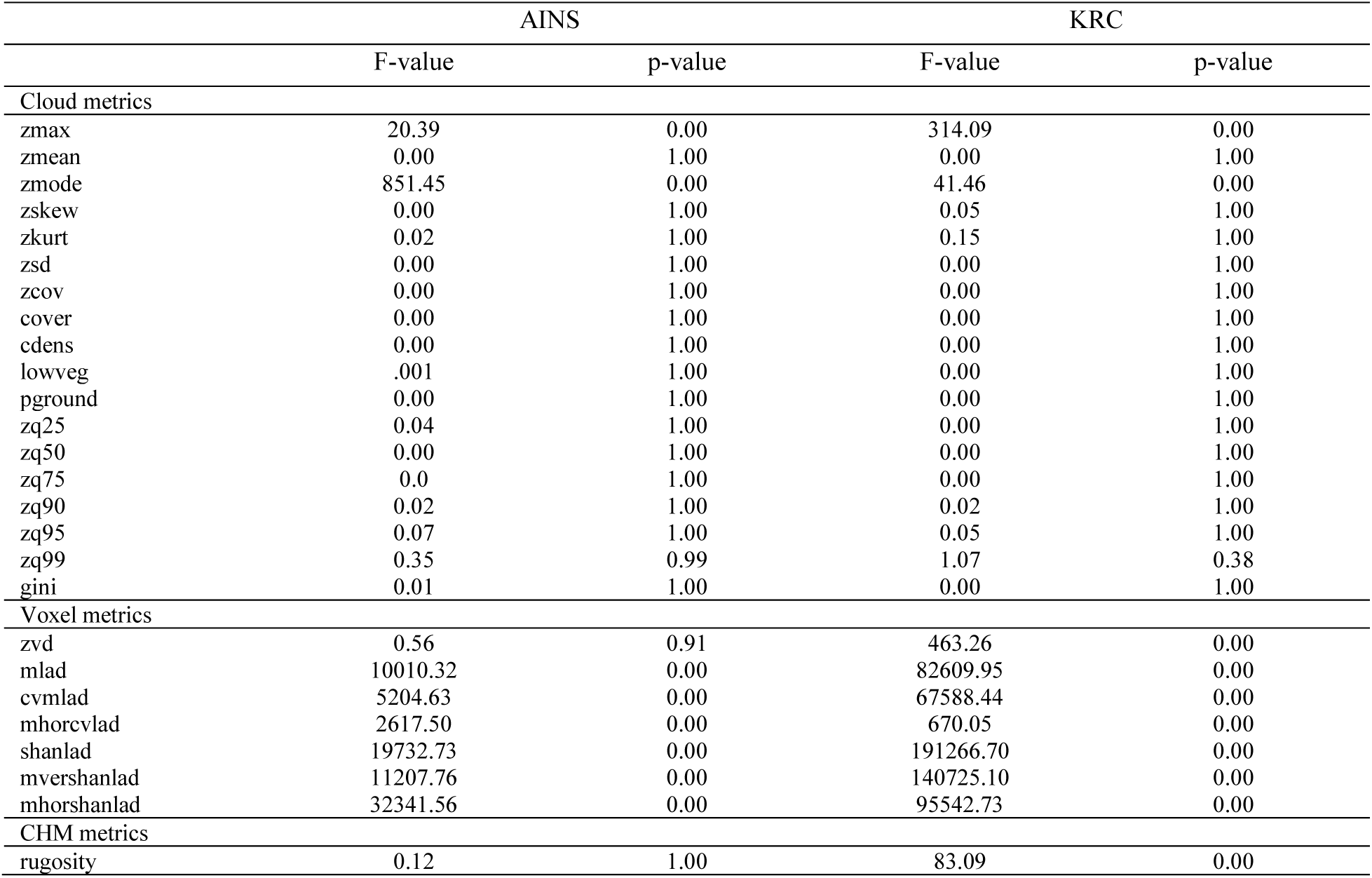
ANOVA results for differences in forest structure metrics derived from the full point cloud based on lidar point density.

**Figure B1-1.**
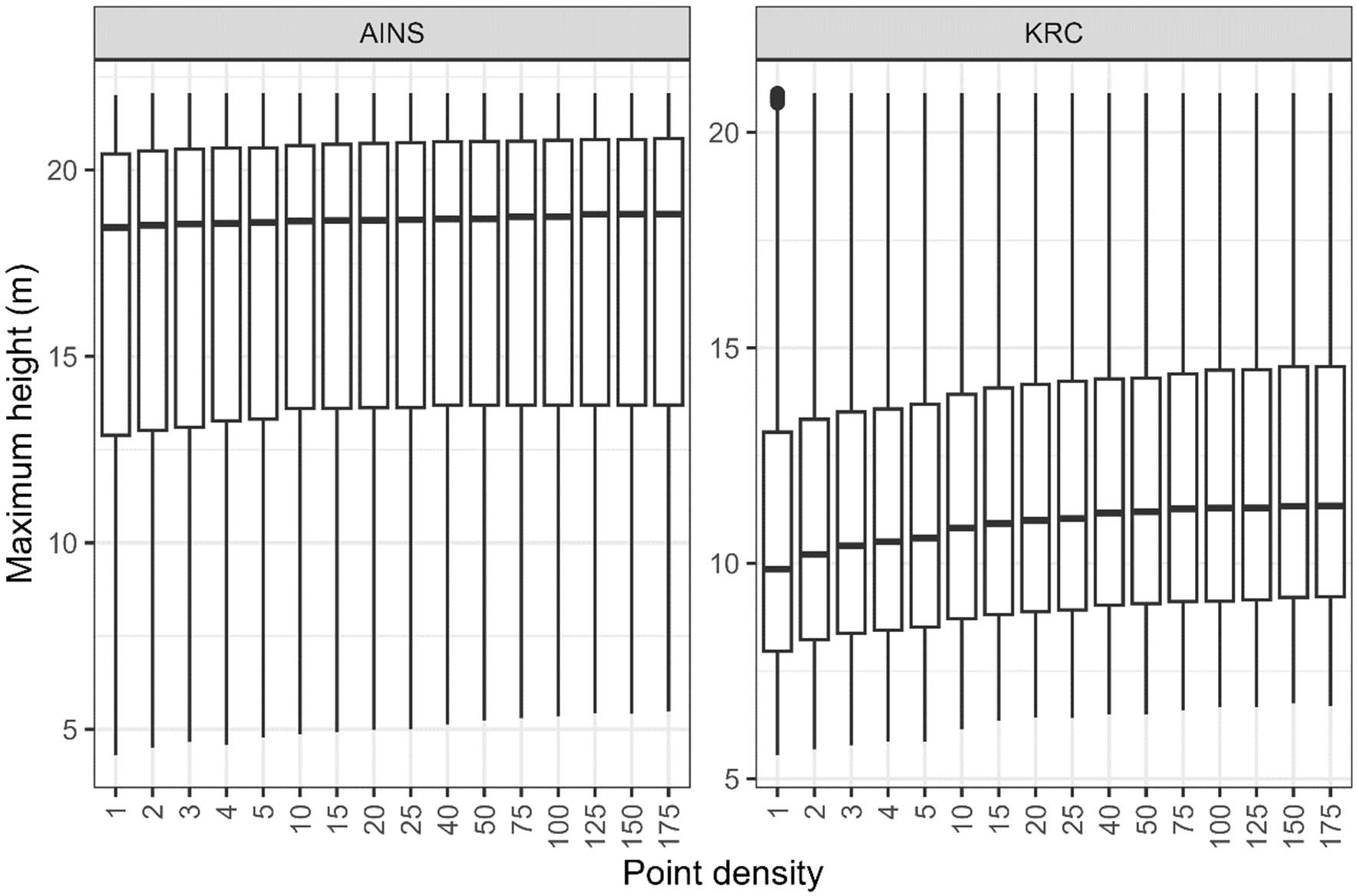
Boxplot of maximum canopy height at AINS and KRC. Bold line in the boxplot represent the median maximum height across all plots. Boxes are the inter-quartile range of maximum height and whiskers are the 95% confidence interval.

**Figure B1-2.**
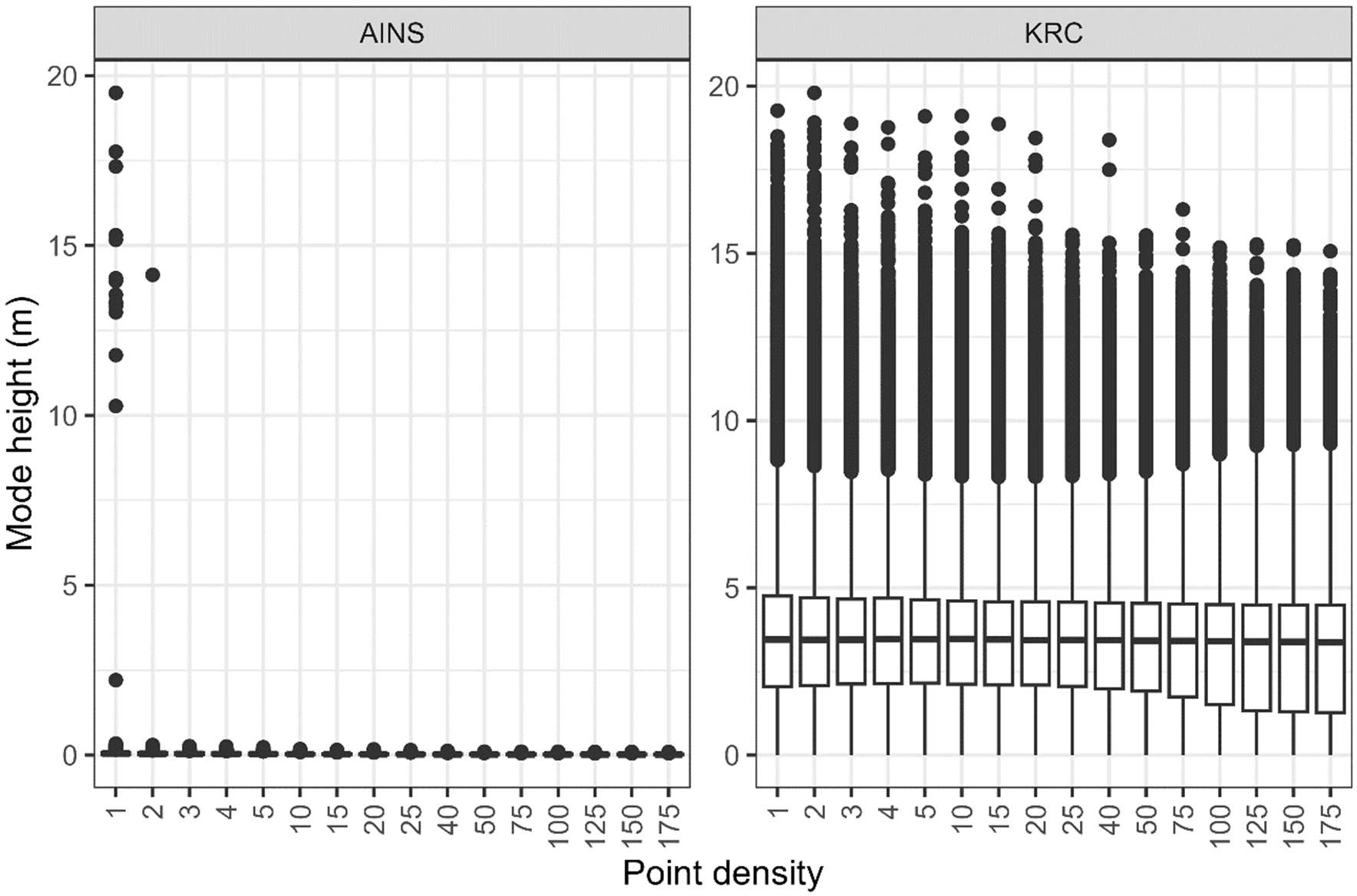
Boxplot of mode canopy height at AINS and KRC. Bold line in the boxplot represent the median mode height across all plots. Boxes are the inter-quartile range of maximum height and whiskers are the 95% confidence interval.

### B2. Correlation results

**Figure B2-1.**
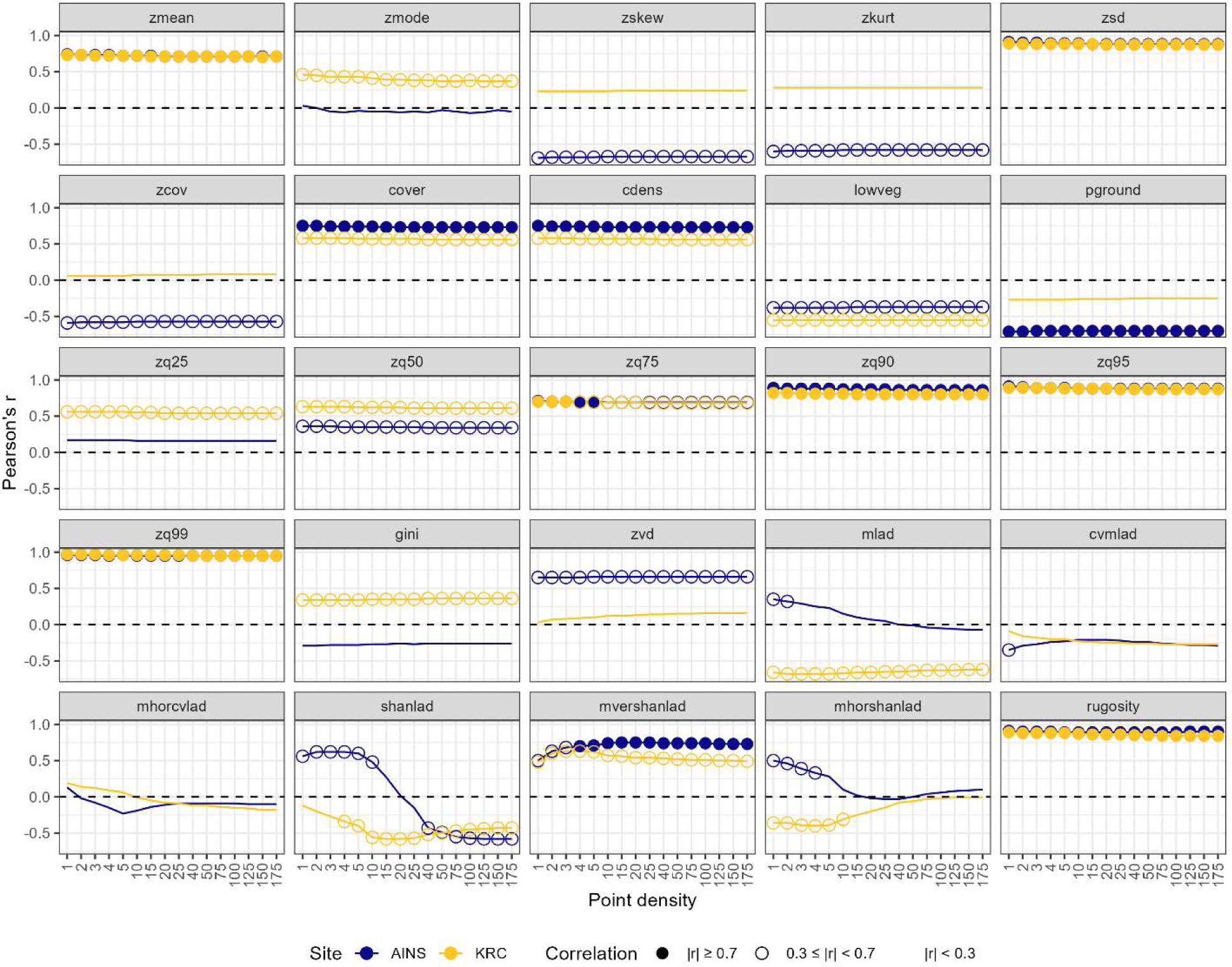
Pearson’s r for correlations between *zmax* and the 25 other forest structure metrics calculated from lidar with differing point densities at AINS (blue) and KRC (yellow). A filled circle indicates a strong correlation (|r| ≥ 0.7), while an open circle indicates a weak correlation (0.3 ≤ |r| < 0.7) and no circle indicates a negligible correlation (|r| < 0.3).

**Figure B2-2.**
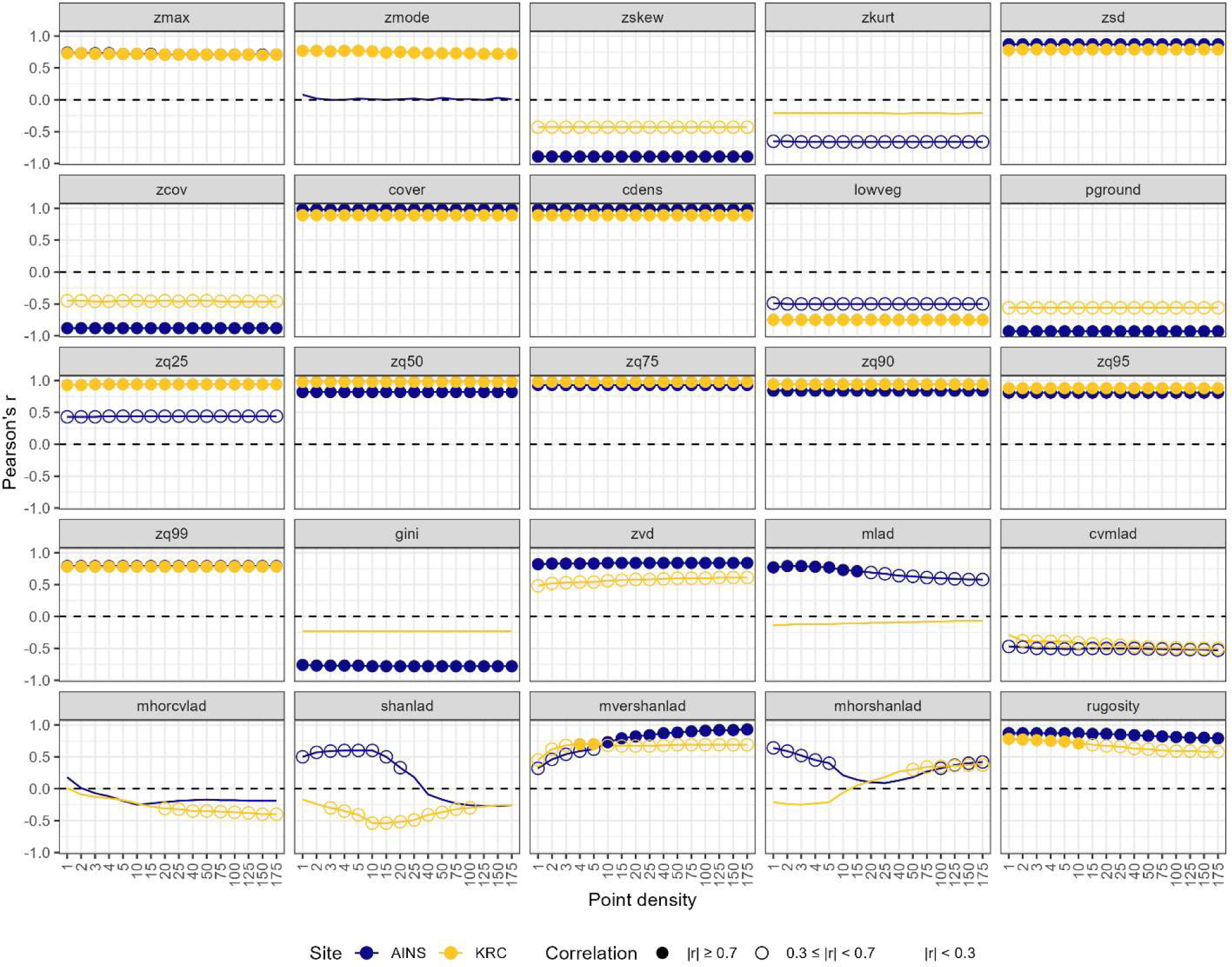
Pearson’s r for correlations between *zmean* and the 25 other forest structure metrics calculated from lidar with differing point densities at AINS (blue) and KRC (yellow). A filled circle indicates a strong correlation (|r| ≥ 0.7), while an open circle indicates a weak correlation (0.3 ≤ |r| < 0.7) and no circle indicates a negligible correlation (|r| < 0.3)..

**Figure B2-3.**
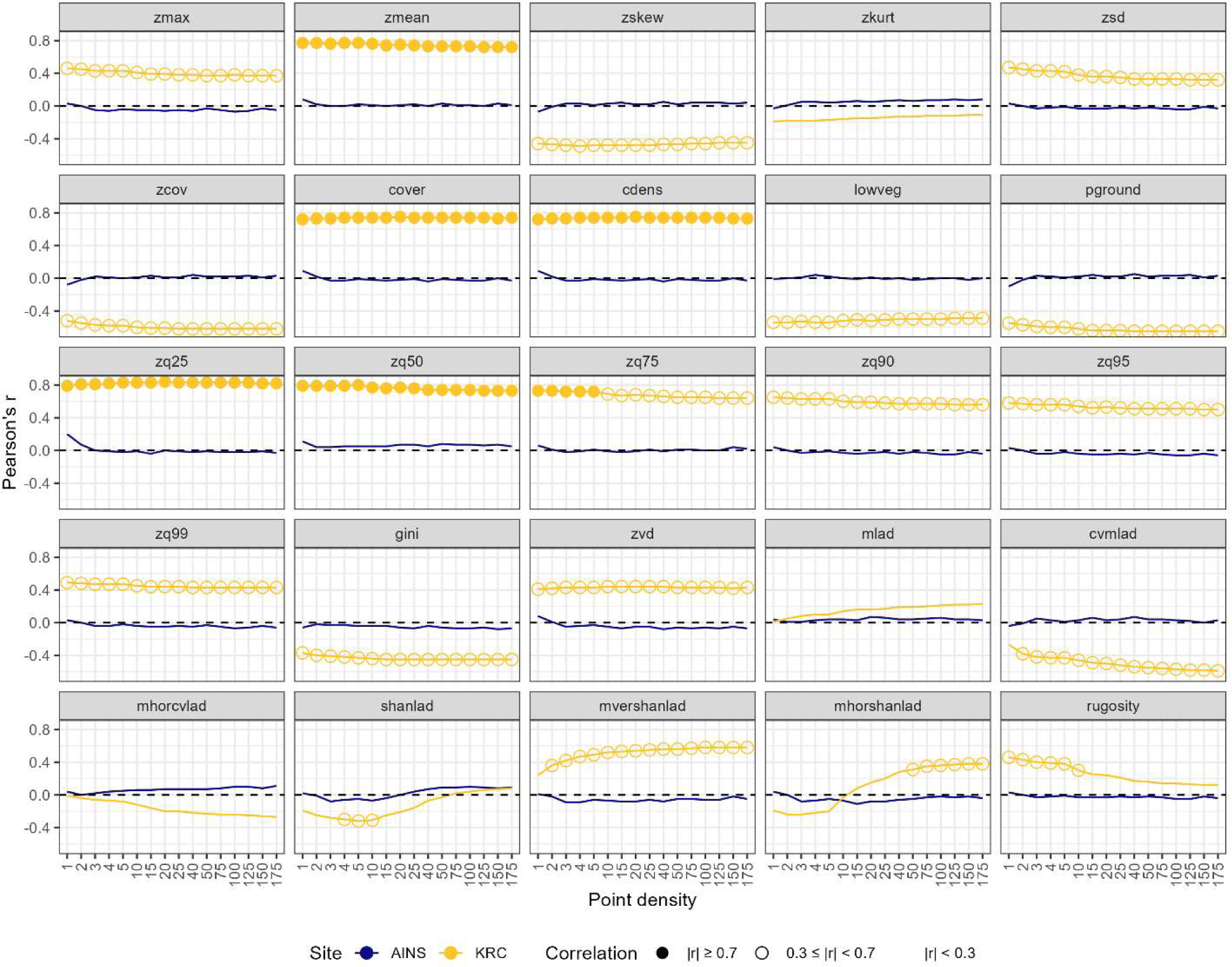
Pearson’s r for correlations between *zmode* and the 25 other forest structure metrics calculated from lidar with differing point densities at AINS (blue) and KRC (yellow). A filled circle indicates a strong correlation (|r| ≥ 0.7), while an open circle indicates a weak correlation (0.3 ≤ |r| < 0.7) and no circle indicates a negligible correlation (|r| < 0.3).

**Figure B2-4.**
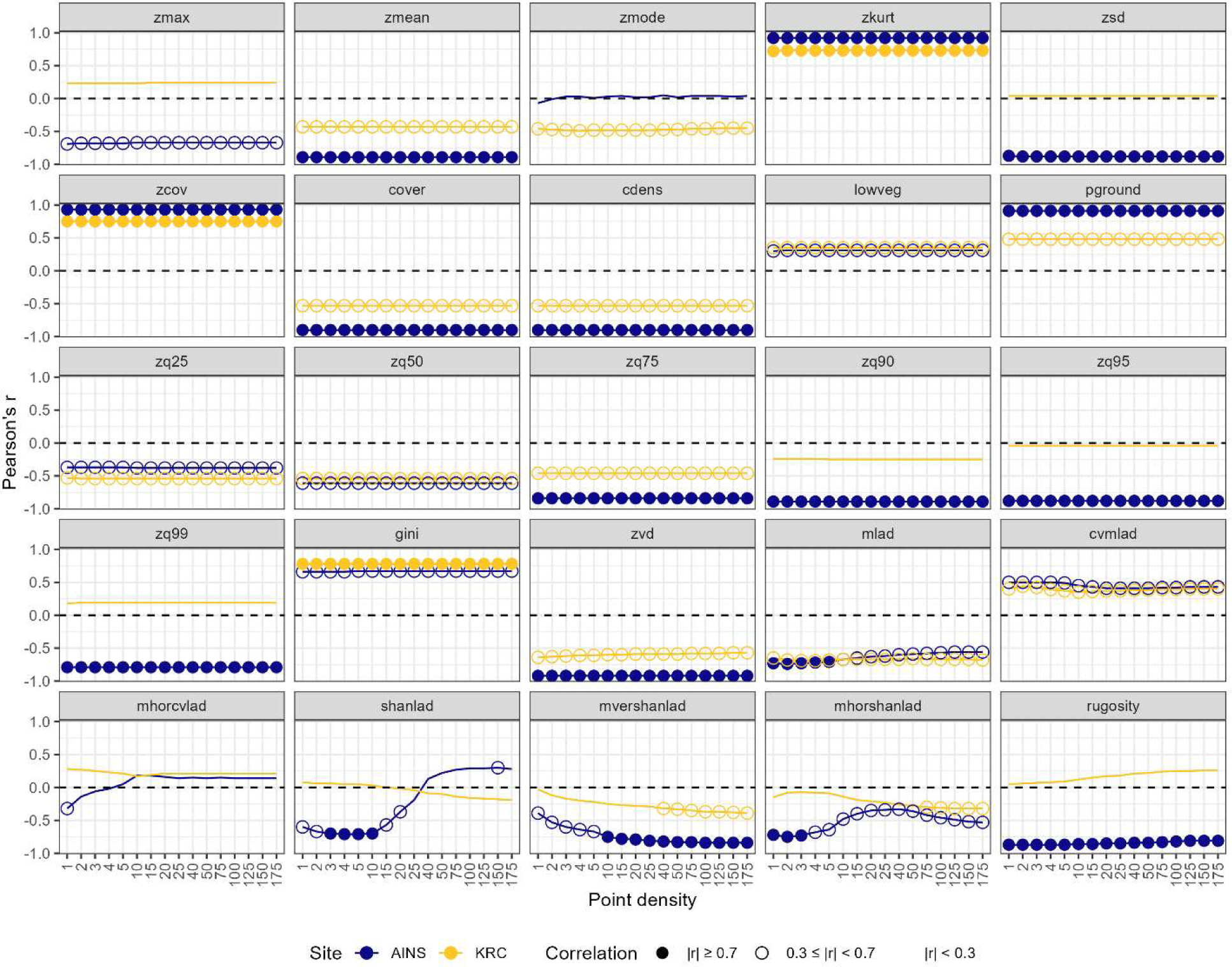
Pearson’s r for correlations between *zskew* and the 25 other forest structure metrics calculated from lidar with differing point densities at AINS (blue) and KRC (yellow). A filled circle indicates a strong correlation (|r| ≥ 0.7), while an open circle indicates a weak correlation (0.3 ≤ |r| < 0.7) and no circle indicates a negligible correlation (|r| < 0.3).

**Figure B2-5.**
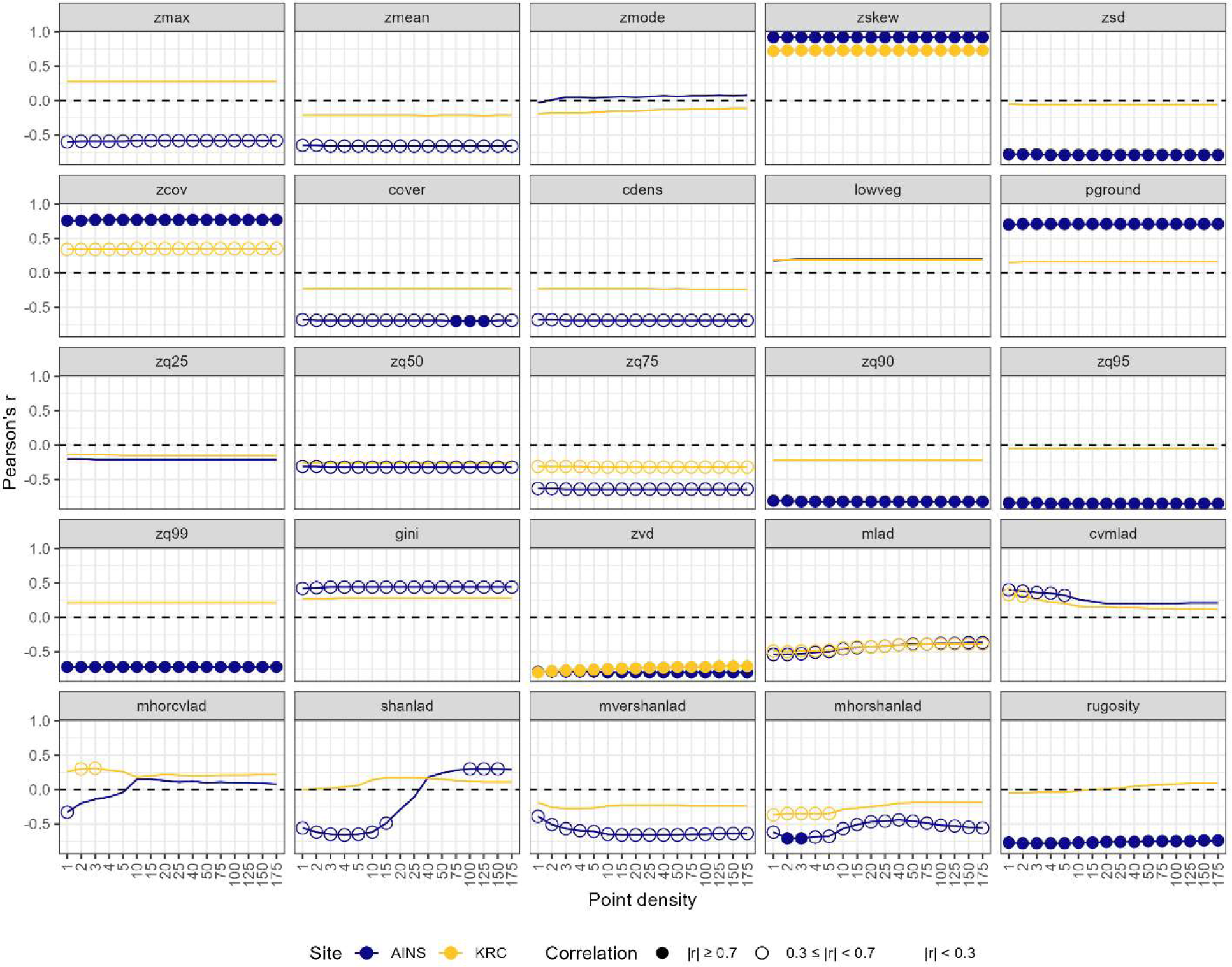
Pearson’s r for correlations between *zkurt* and the 25 other forest structure metrics calculated from lidar with differing point densities at AINS (blue) and KRC (yellow). A filled circle indicates a strong correlation (|r| ≥ 0.7), while an open circle indicates a weak correlation (0.3 ≤ |r| < 0.7) and no circle indicates a negligible correlation (|r| < 0.3).

**Figure B2-6.**
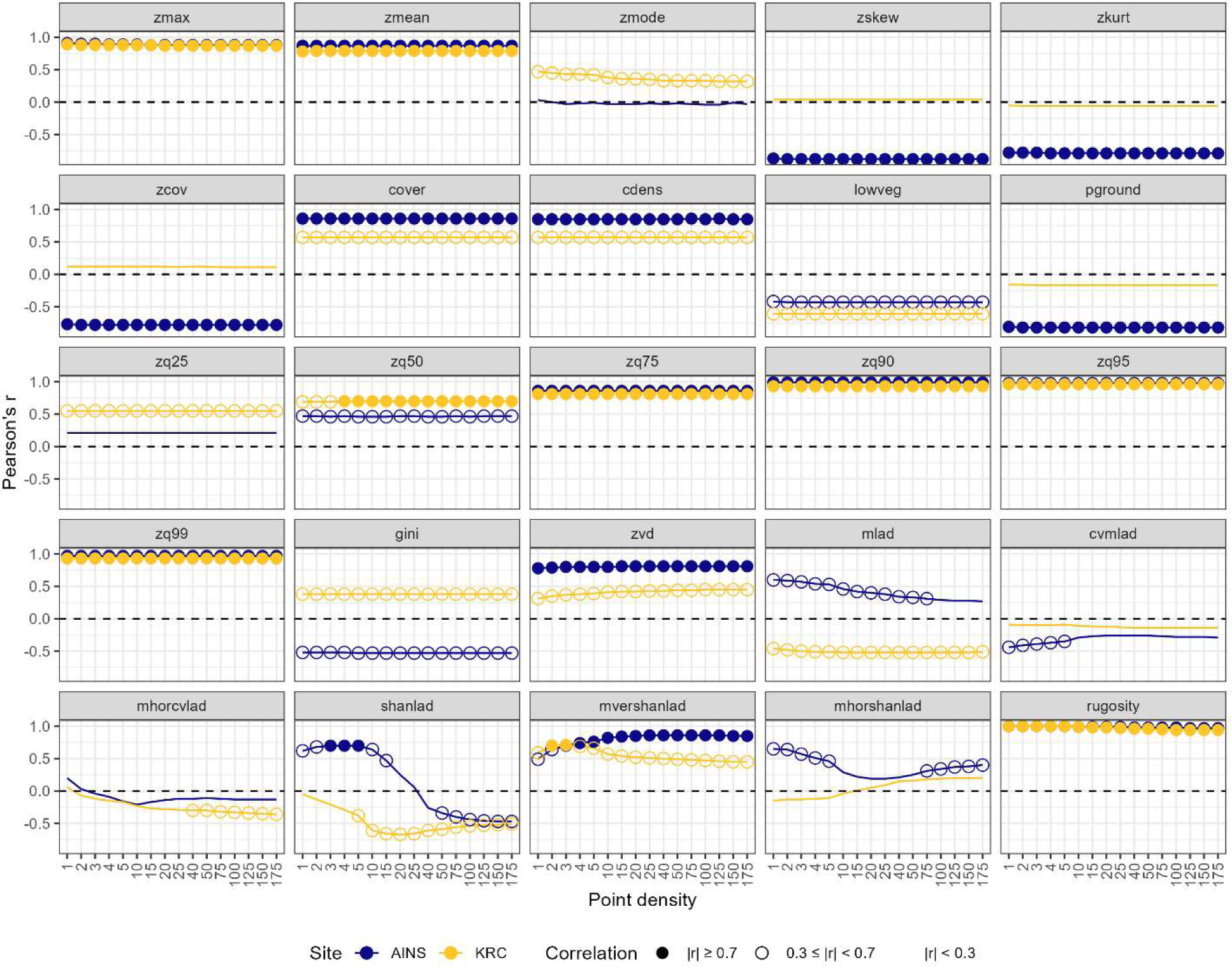
Pearson’s r for correlations between *zsd* and the 25 other forest structure metrics calculated from lidar with differing point densities at AINS (blue) and KRC (yellow). A filled circle indicates a strong correlation (|r| ≥ 0.7), while an open circle indicates a weak correlation (0.3 ≤ |r| < 0.7) and no circle indicates a negligible correlation (|r| < 0.3).

**Figure B2-7.**
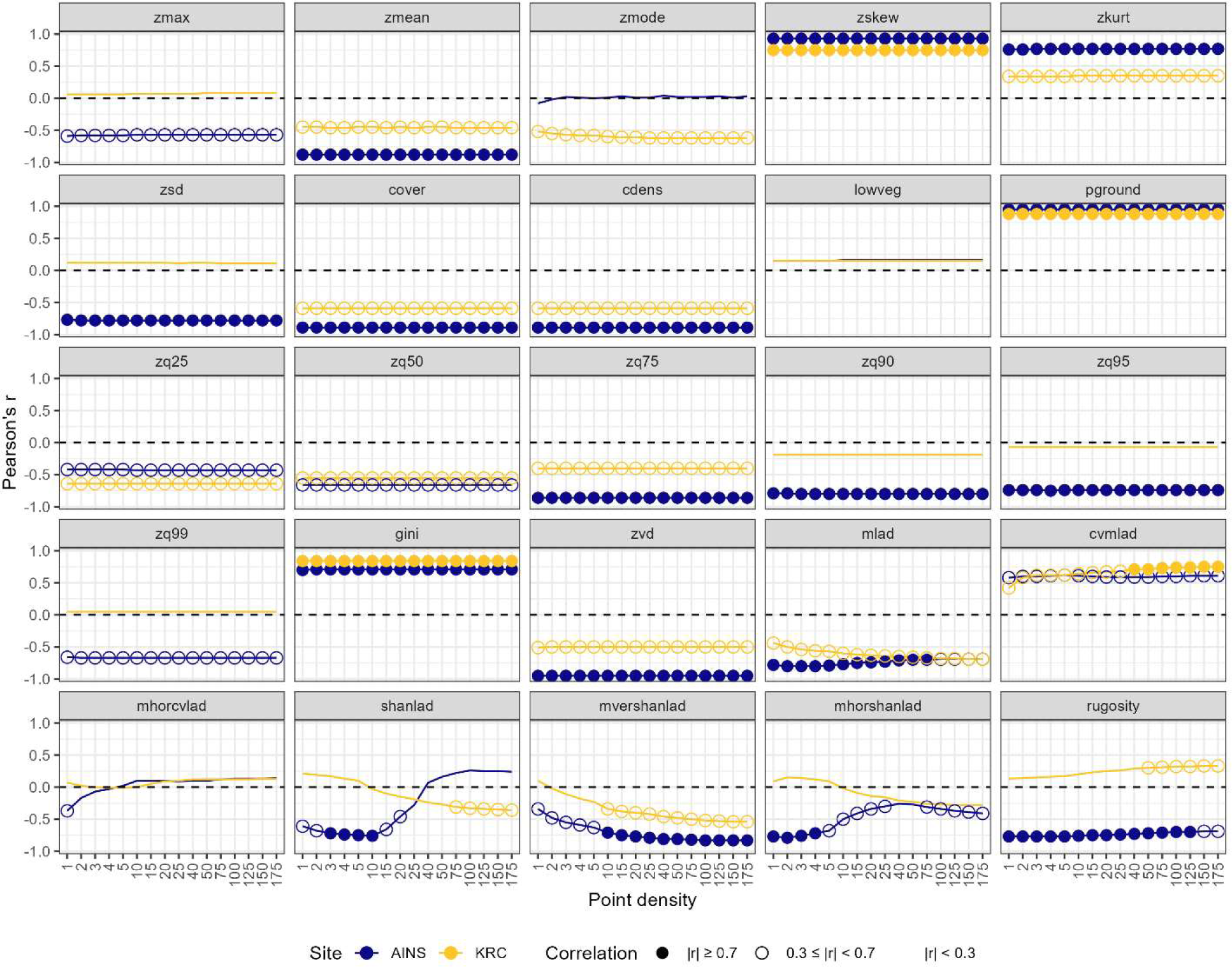
Pearson’s r for correlations between *zcov* and the 25 other forest structure metrics calculated from lidar with differing point densities at AINS (blue) and KRC (yellow). A filled circle indicates a strong correlation (|r| ≥ 0.7), while an open circle indicates a weak correlation (0.3 ≤ |r| < 0.7) and no circle indicates a negligible correlation (|r| < 0.3).

**Figure B2-8.**
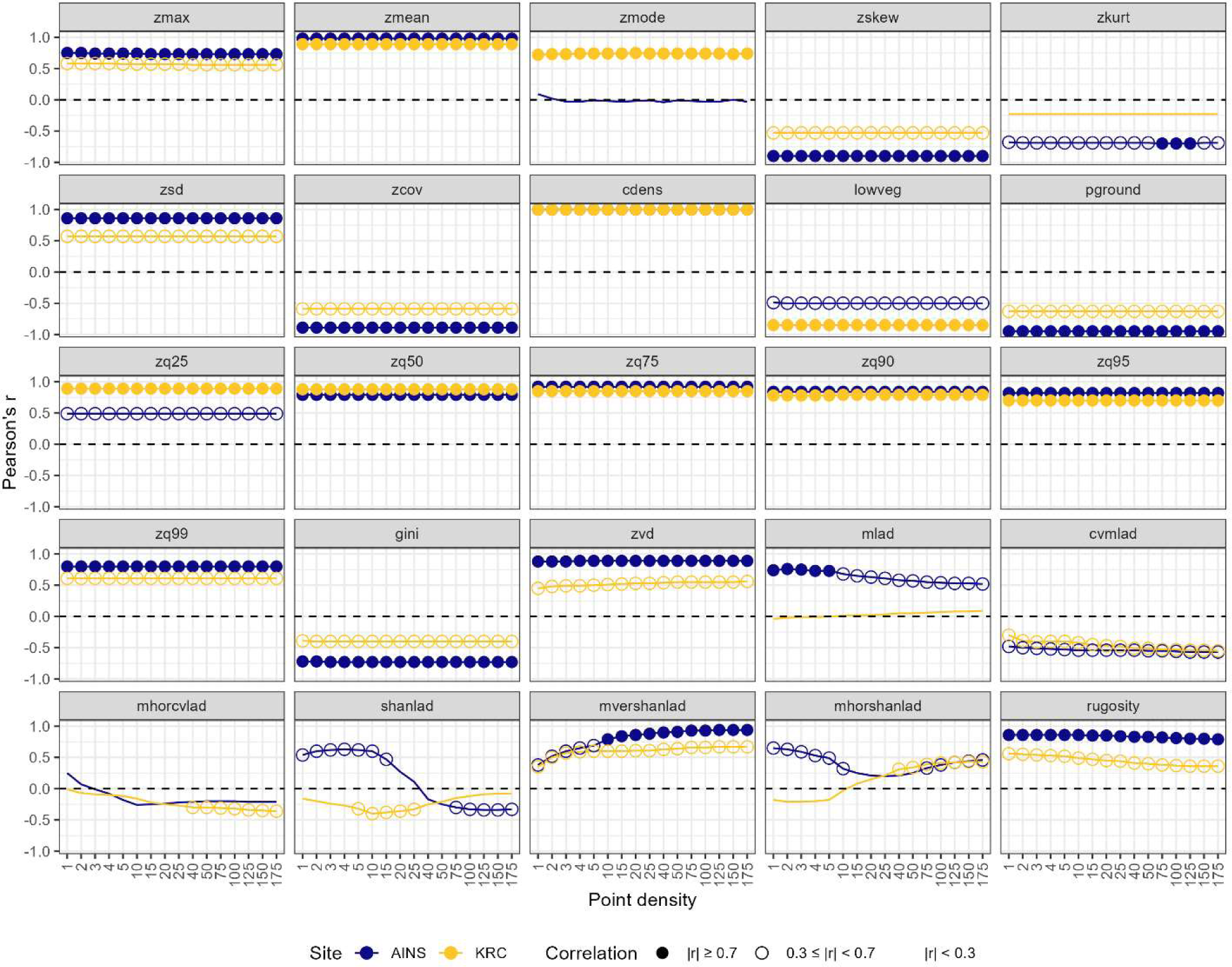
Pearson’s r for correlations between *cover* and the 25 other forest structure metrics calculated from lidar with differing point densities at AINS (blue) and KRC (yellow). A filled circle indicates a strong correlation (|r| ≥ 0.7), while an open circle indicates a weak correlation (0.3 ≤ |r| < 0.7) and no circle indicates a negligible correlation (|r| < 0.3).

**Figure B2-9.**
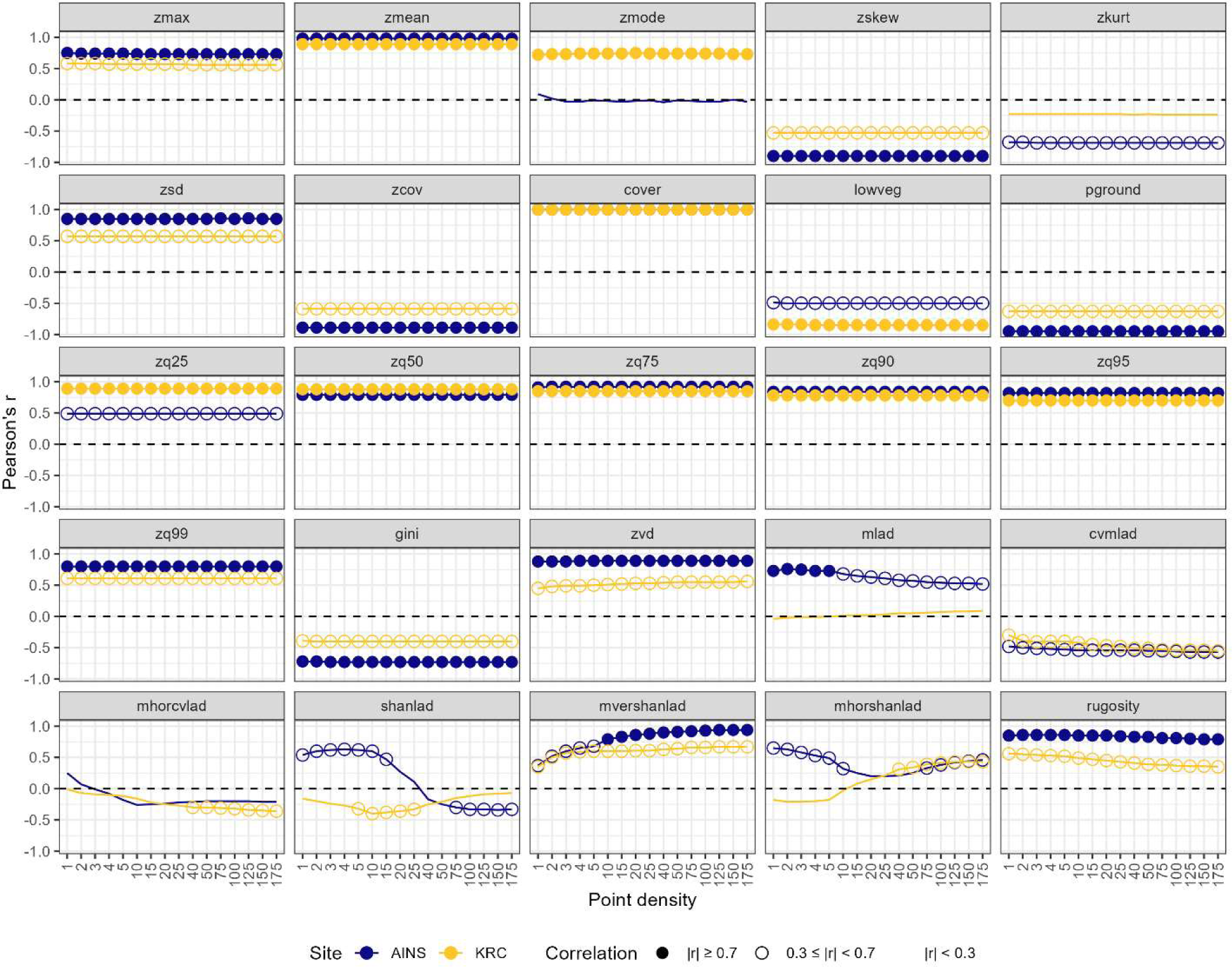
Pearson’s r for correlations between *cdens* and the 25 other forest structure metrics calculated from lidar with differing point densities at AINS (blue) and KRC (yellow). A filled circle indicates a strong correlation (|r| ≥ 0.7), while an open circle indicates a weak correlation (0.3 ≤ |r| < 0.7) and no circle indicates a negligible correlation (|r| < 0.3).

**Figure B2-10.**
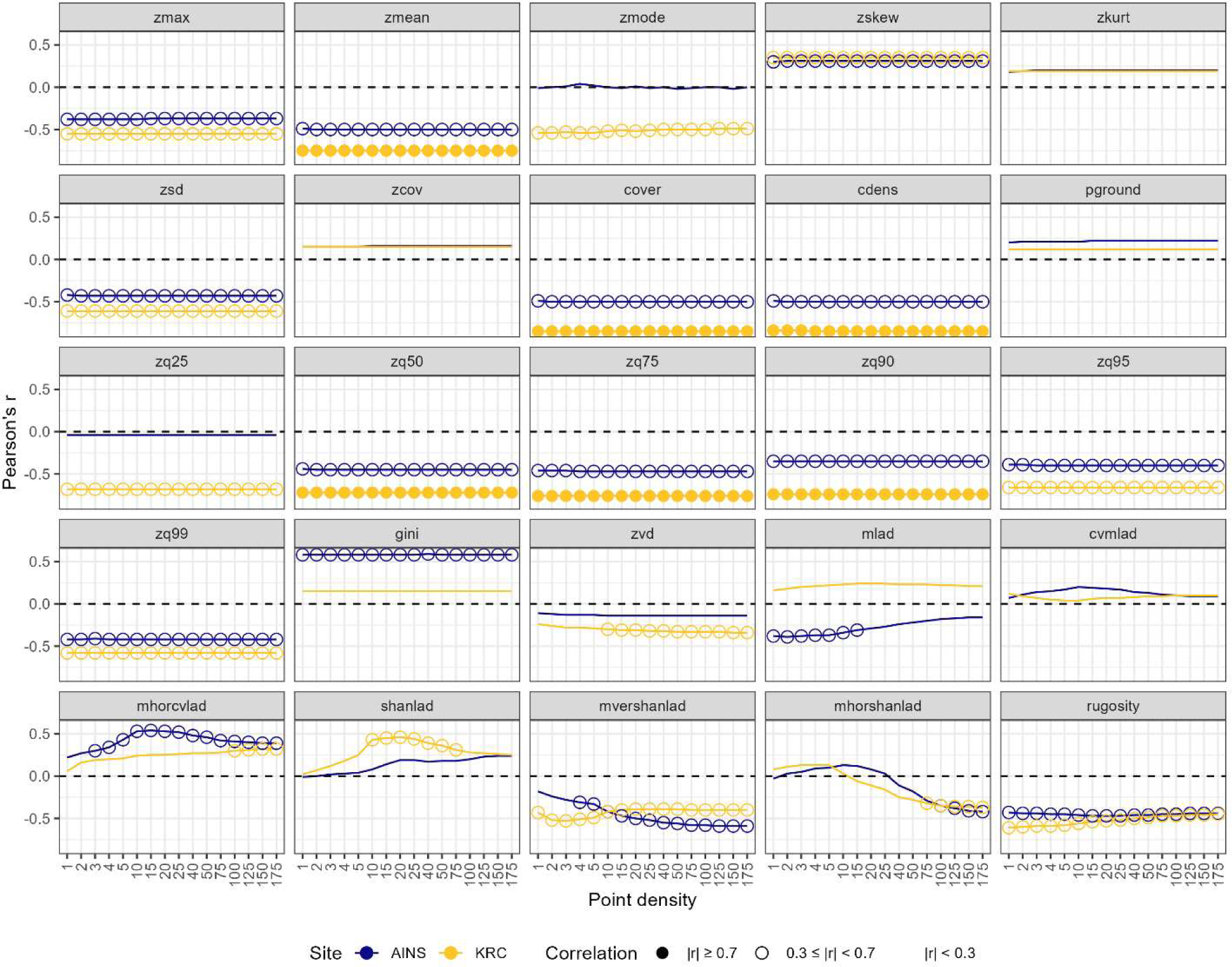
Pearson’s r for correlations between *lowveg* and the 25 other forest structure metrics calculated from lidar with differing point densities at AINS (blue) and KRC (yellow). A filled circle indicates a strong correlation (|r| ≥ 0.7), while an open circle indicates a weak correlation (0.3 ≤ |r| < 0.7) and no circle indicates a negligible correlation (|r| < 0.3).

**Figure B2-11.**
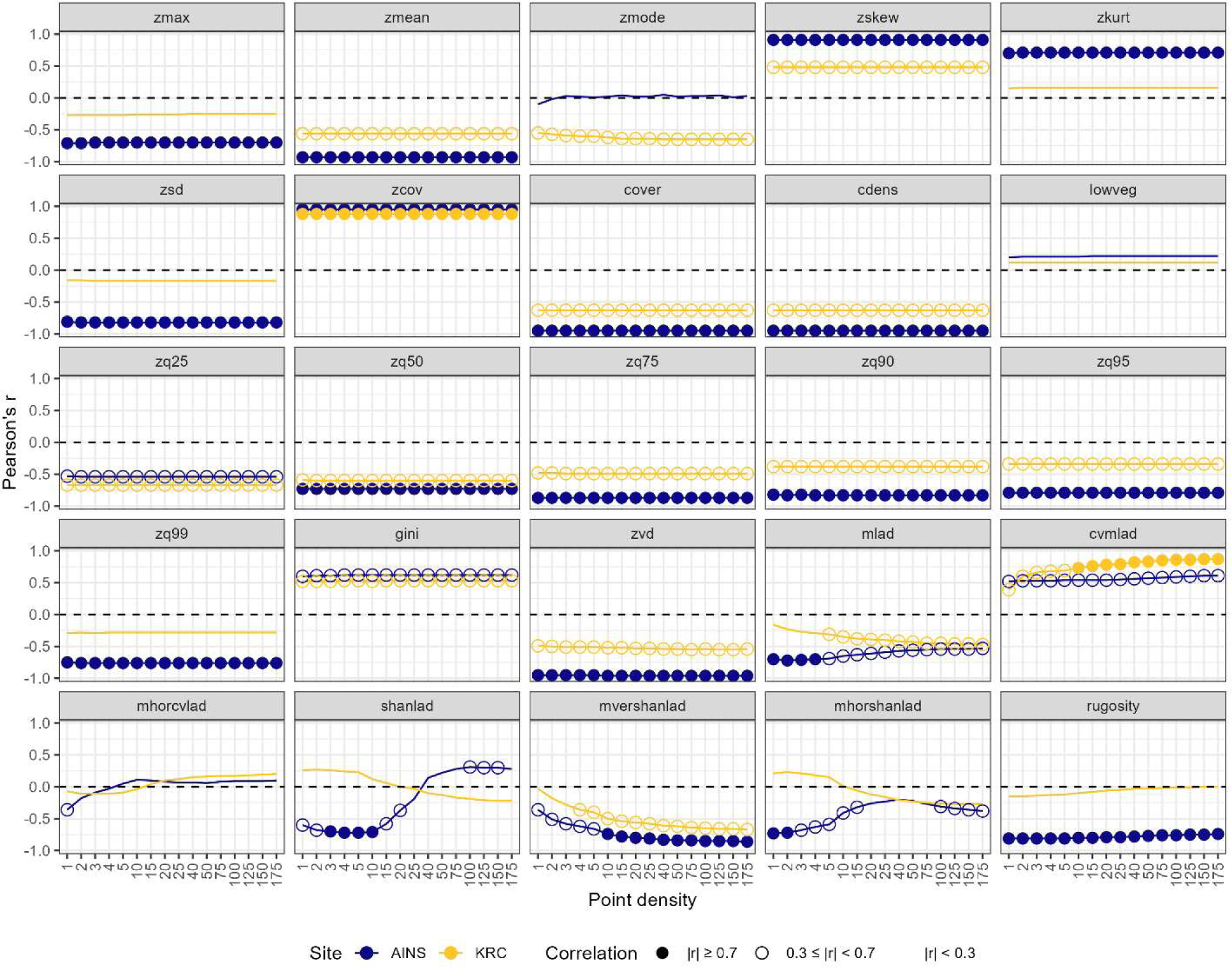
Pearson’s r for correlations between *pground* and the 25 other forest structure metrics calculated from lidar with differing point densities at AINS (blue) and KRC (yellow). A filled circle indicates a strong correlation (|r| ≥ 0.7), while an open circle indicates a weak correlation (0.3 ≤ |r| < 0.7) and no circle indicates a negligible correlation (|r| < 0.3).

**Figure B2-12.**
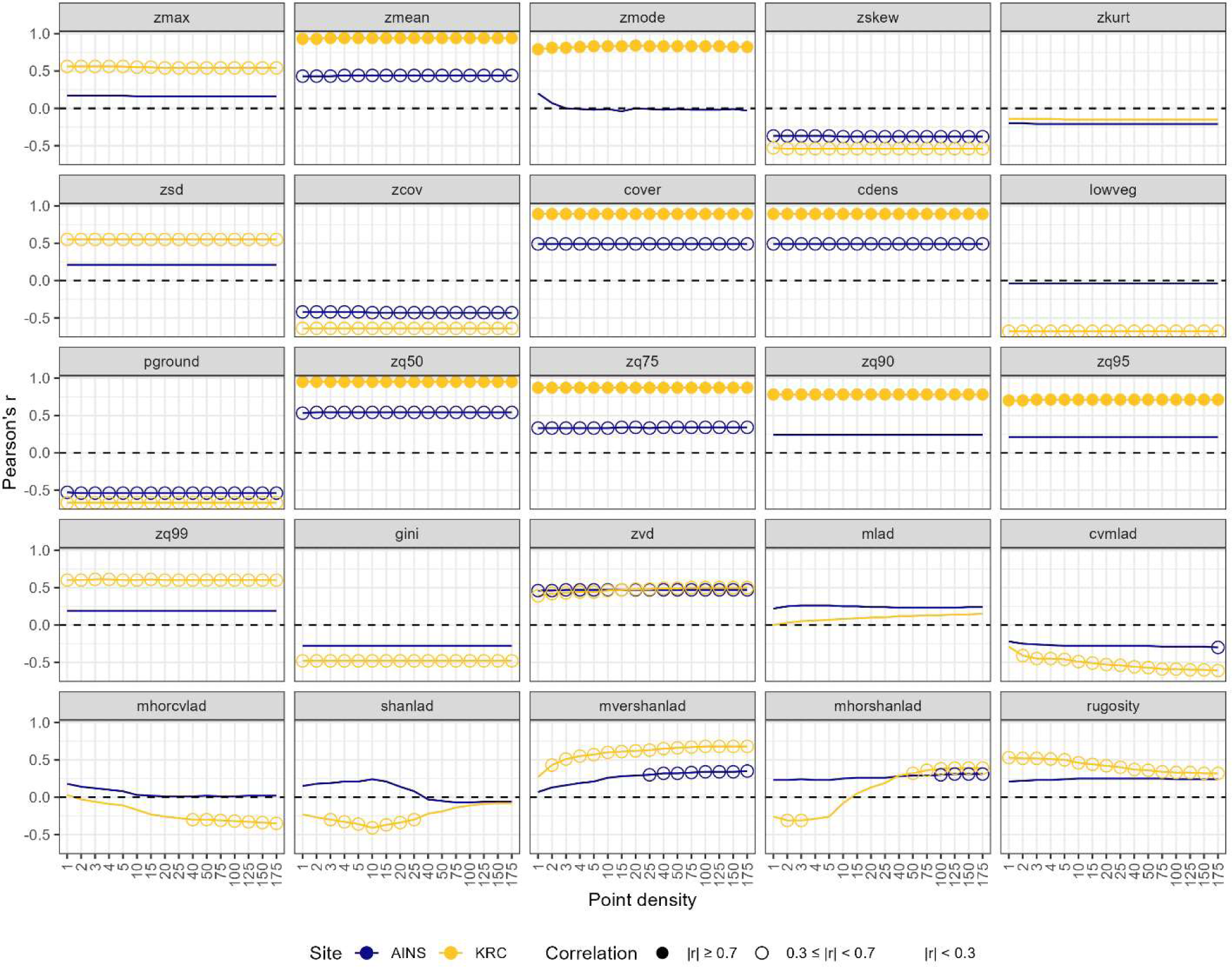
Pearson’s r for correlations between *zq25* and the 25 other forest structure metrics calculated from lidar with differing point densities at AINS (blue) and KRC (yellow). A filled circle indicates a strong correlation (|r| ≥ 0.7), while an open circle indicates a weak correlation (0.3 ≤ |r| < 0.7) and no circle indicates a negligible correlation (|r| < 0.3).

**Figure B2-13.**
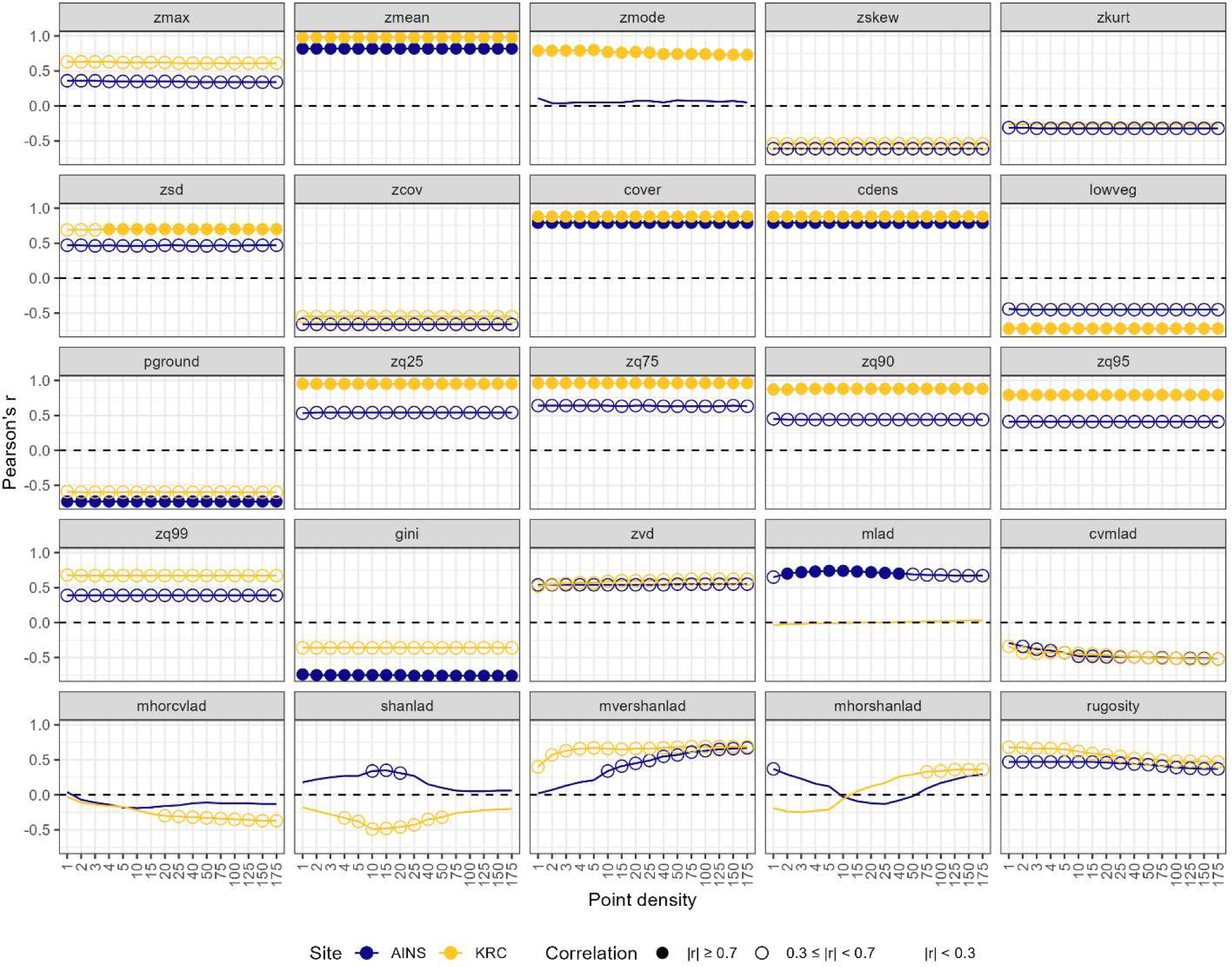
Pearson’s r for correlations between *zq50* and the 25 other forest structure metrics calculated from lidar with differing point densities at AINS (blue) and KRC (yellow). A filled circle indicates a strong correlation (|r| ≥ 0.7), while an open circle indicates a weak correlation (0.3 ≤ |r| < 0.7) and no circle indicates a negligible correlation (|r| < 0.3).

**Figure B2-14.**
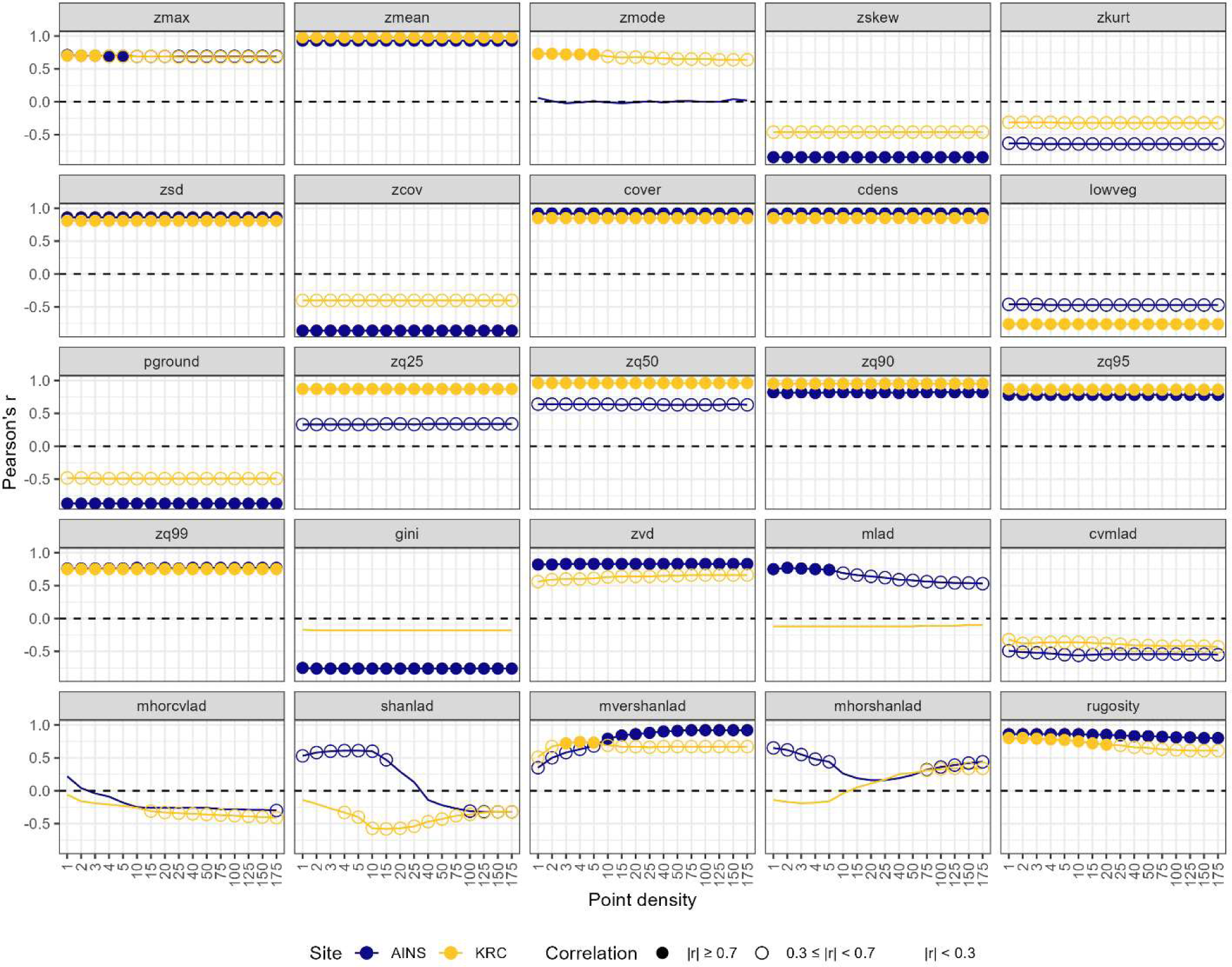
Pearson’s r for correlations between *zq75* and the 25 other forest structure metrics calculated from lidar with differing point densities at AINS (blue) and KRC (yellow). A filled circle indicates a strong correlation (|r| ≥ 0.7), while an open circle indicates a weak correlation (0.3 ≤ |r| < 0.7) and no circle indicates a negligible correlation (|r| < 0.3).

**Figure B2-15.**
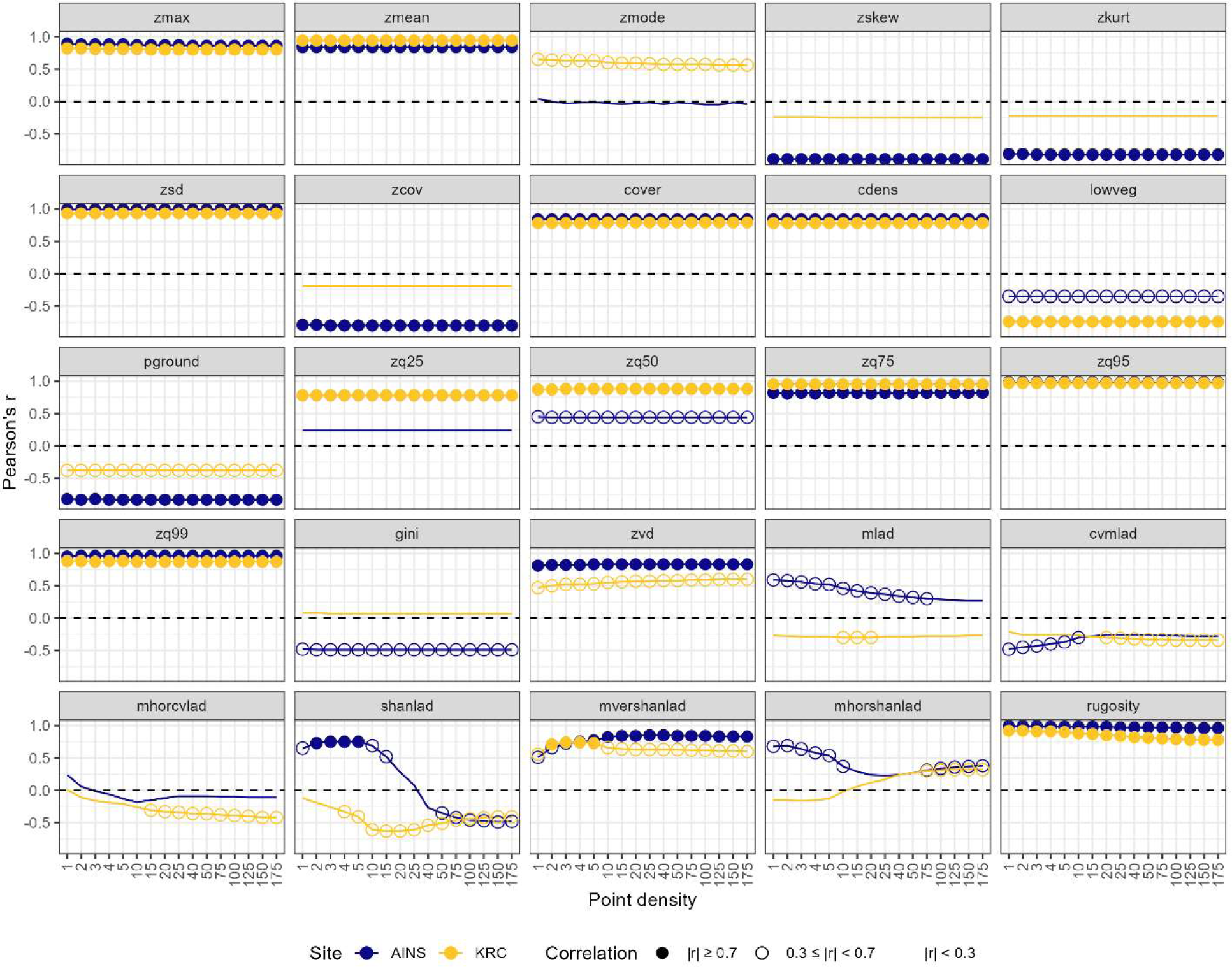
Pearson’s r for correlations between *zq90* and the 25 other forest structure metrics calculated from lidar with differing point densities at AINS (blue) and KRC (yellow). A filled circle indicates a strong correlation (|r| ≥ 0.7), while an open circle indicates a weak correlation (0.3 ≤ |r| < 0.7) and no circle indicates a negligible correlation (|r| < 0.3).

**Figure B2-16.**
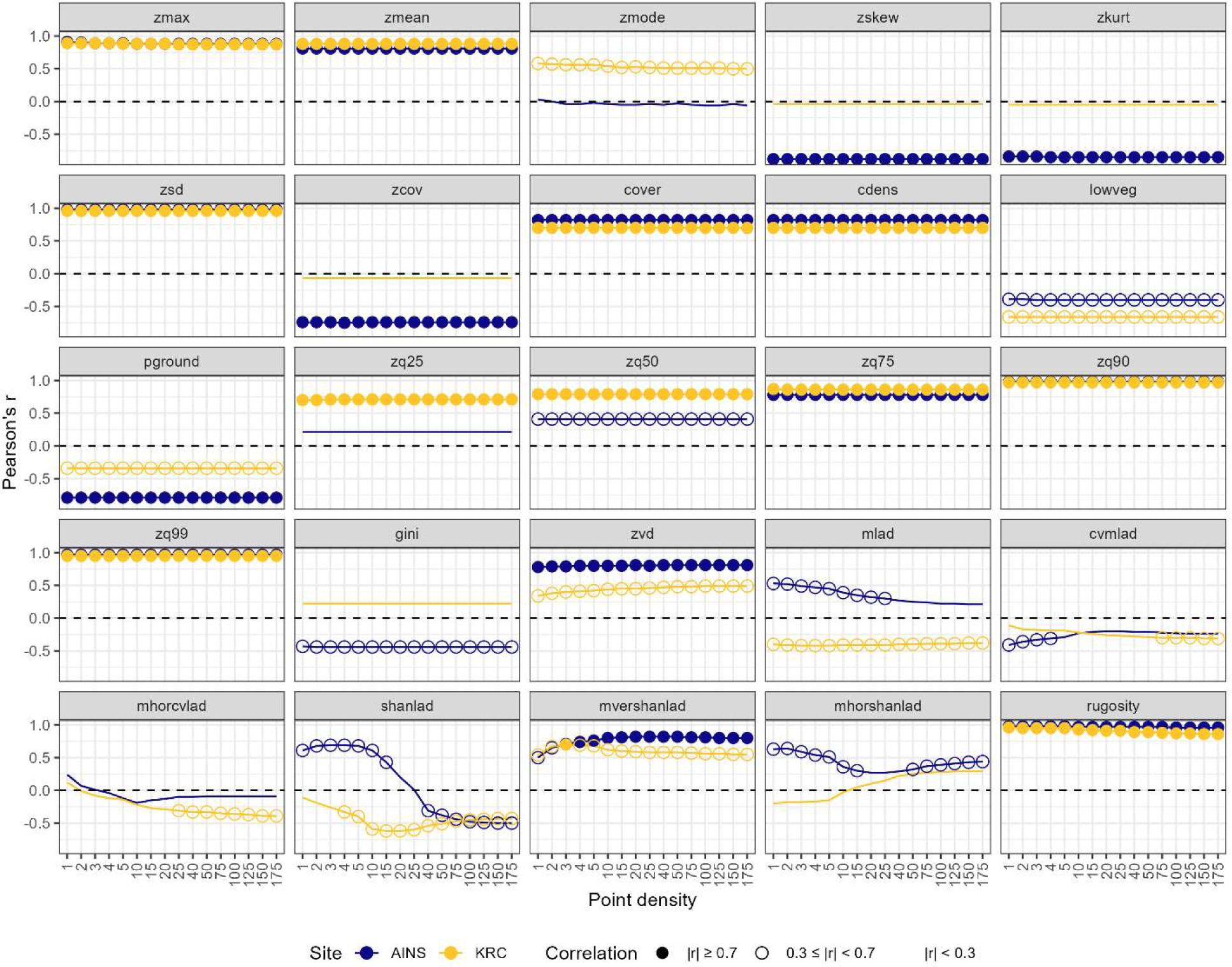
Pearson’s r for correlations between *zq95* and the 25 other forest structure metrics calculated from lidar with differing point densities at AINS (blue) and KRC (yellow). A filled circle indicates a strong correlation (|r| ≥ 0.7), while an open circle indicates a weak correlation (0.3 ≤ |r| < 0.7) and no circle indicates a negligible correlation (|r| < 0.3).

**Figure B2-17.**
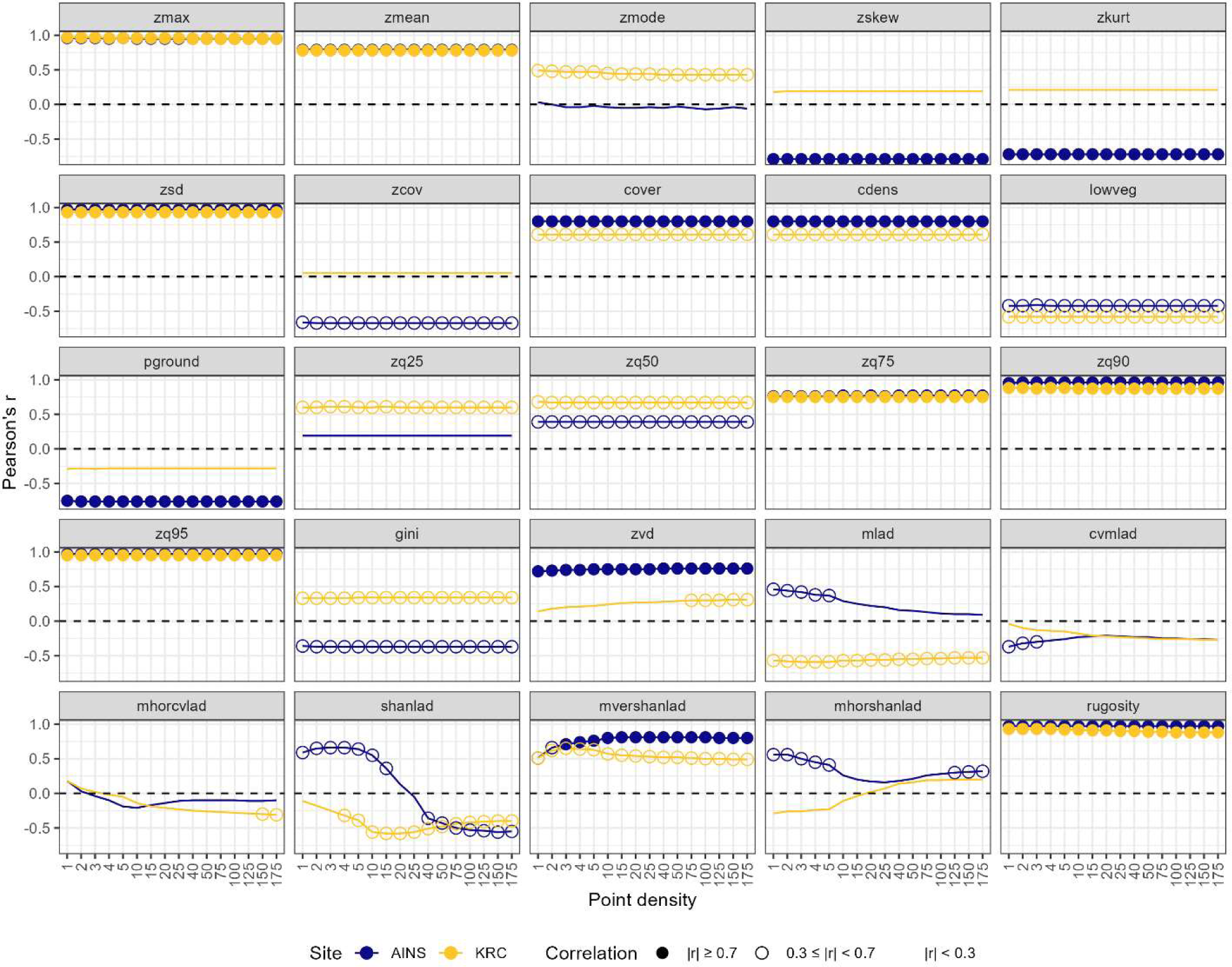
Pearson’s r for correlations between *zq99* and the 25 other forest structure metrics calculated from lidar with differing point densities at AINS (blue) and KRC (yellow). A filled circle indicates a strong correlation (|r| ≥ 0.7), while an open circle indicates a weak correlation (0.3 ≤ |r| < 0.7) and no circle indicates a negligible correlation (|r| < 0.3).

**Figure B2-18.**
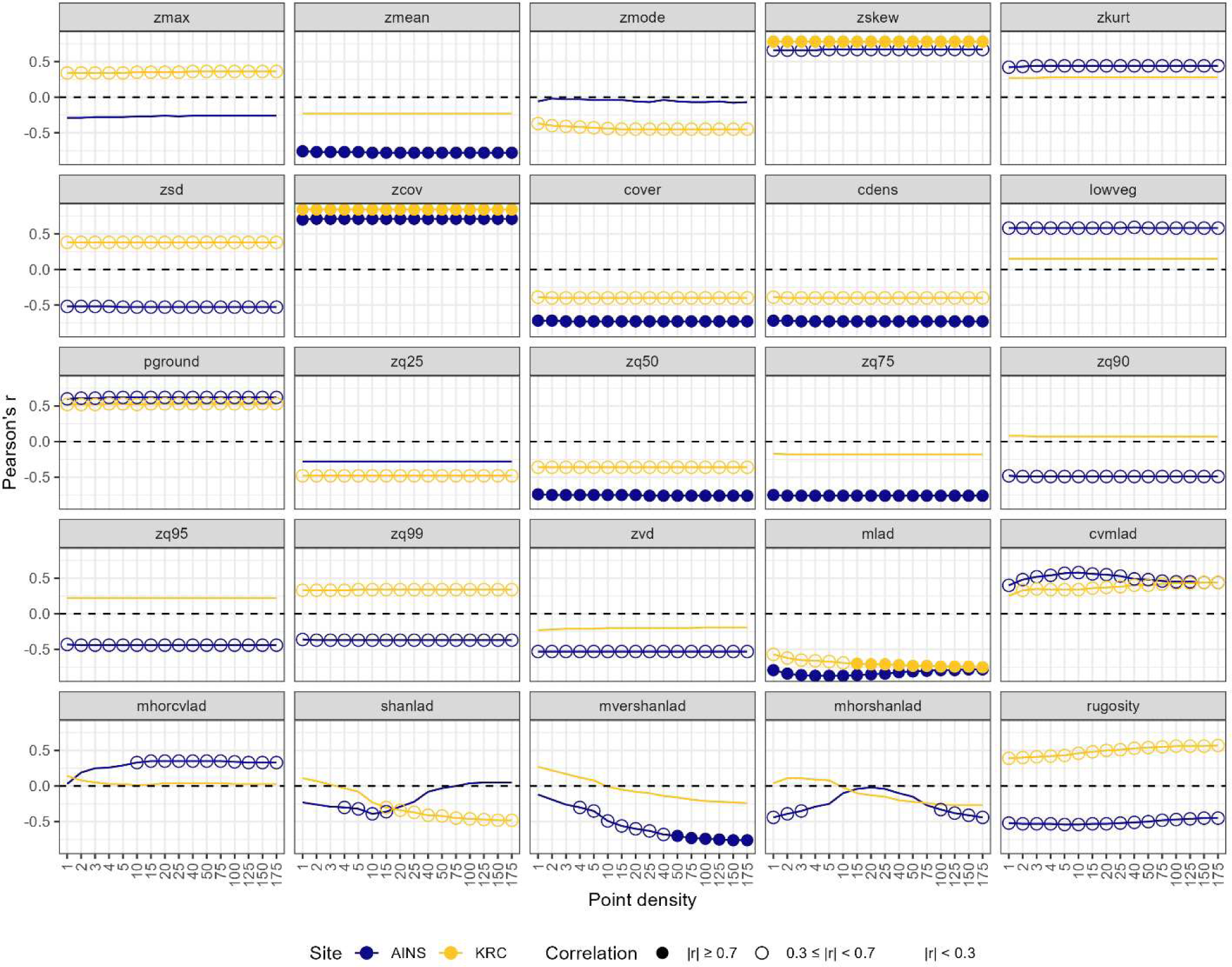
Pearson’s r for correlations between *gini* and the 25 other forest structure metrics calculated from lidar with differing point densities at AINS (blue) and KRC (yellow). A filled circle indicates a strong correlation (|r| ≥ 0.7), while an open circle indicates a weak correlation (0.3 ≤ |r| < 0.7) and no circle indicates a negligible correlation (|r| < 0.3).

**Figure B2-19.**
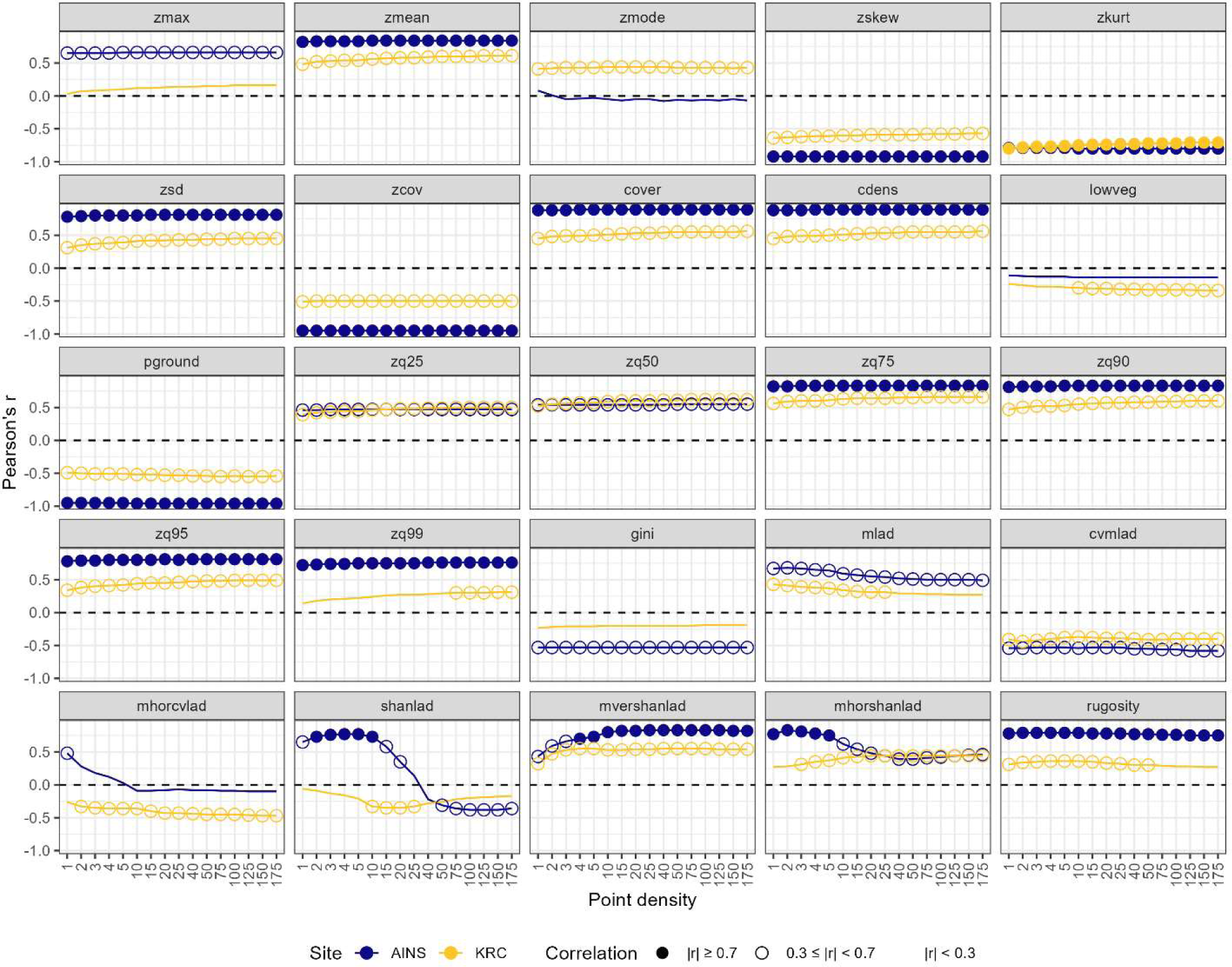
Pearson’s r for correlations between *zvd* and the 25 other forest structure metrics calculated from lidar with differing point densities at AINS (blue) and KRC (yellow). A filled circle indicates a strong correlation (|r| ≥ 0.7), while an open circle indicates a weak correlation (0.3 ≤ |r| < 0.7) and no circle indicates a negligible correlation (|r| < 0.3).

**Figure B2-20.**
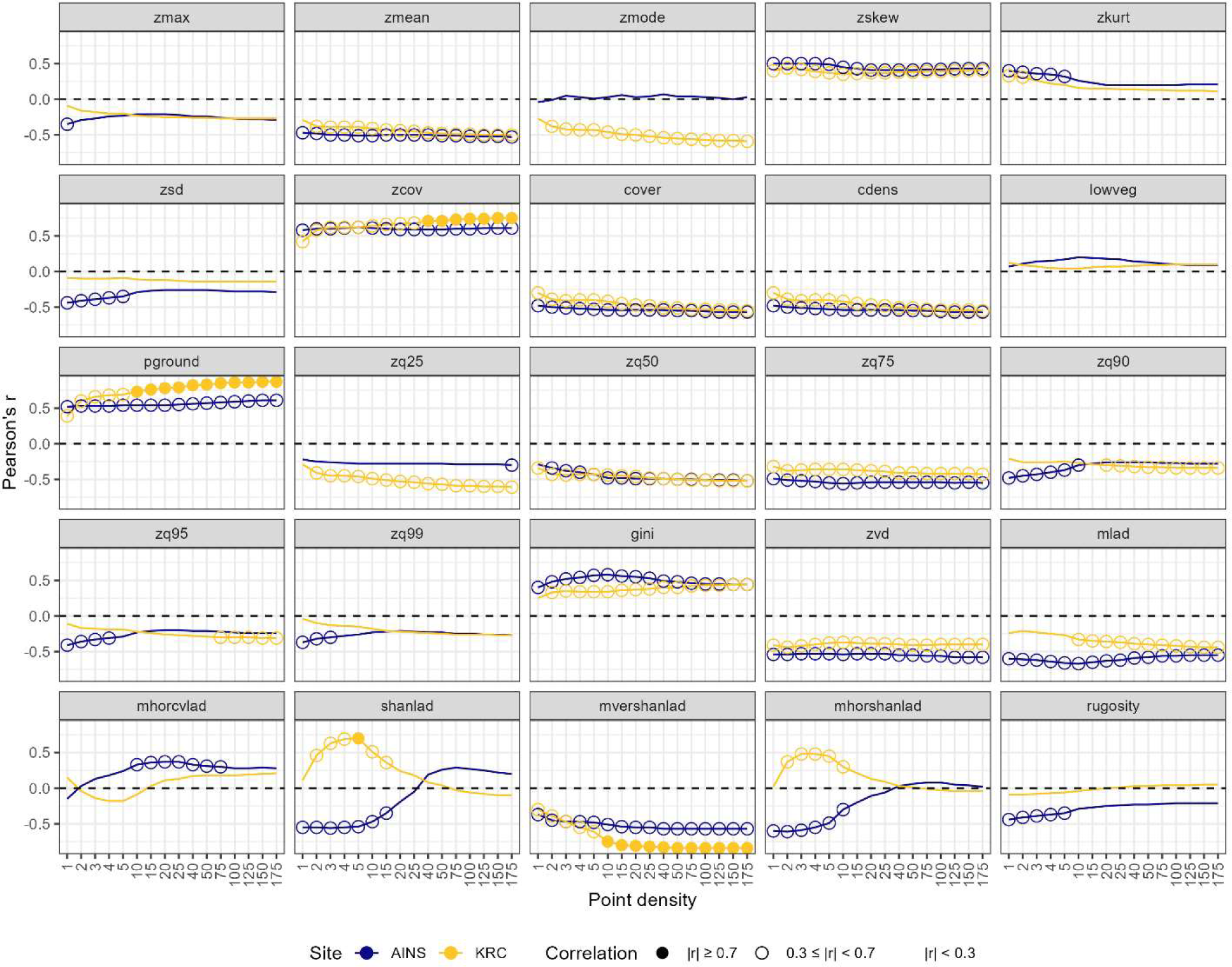
Pearson’s r for correlations between *cvmlad* and the 25 other forest structure metrics calculated from lidar with differing point densities at AINS (blue) and KRC (yellow). A filled circle indicates a strong correlation (|r| ≥ 0.7), while an open circle indicates a weak correlation (0.3 ≤ |r| < 0.7) and no circle indicates a negligible correlation (|r| < 0.3).

**Figure B2-21.**
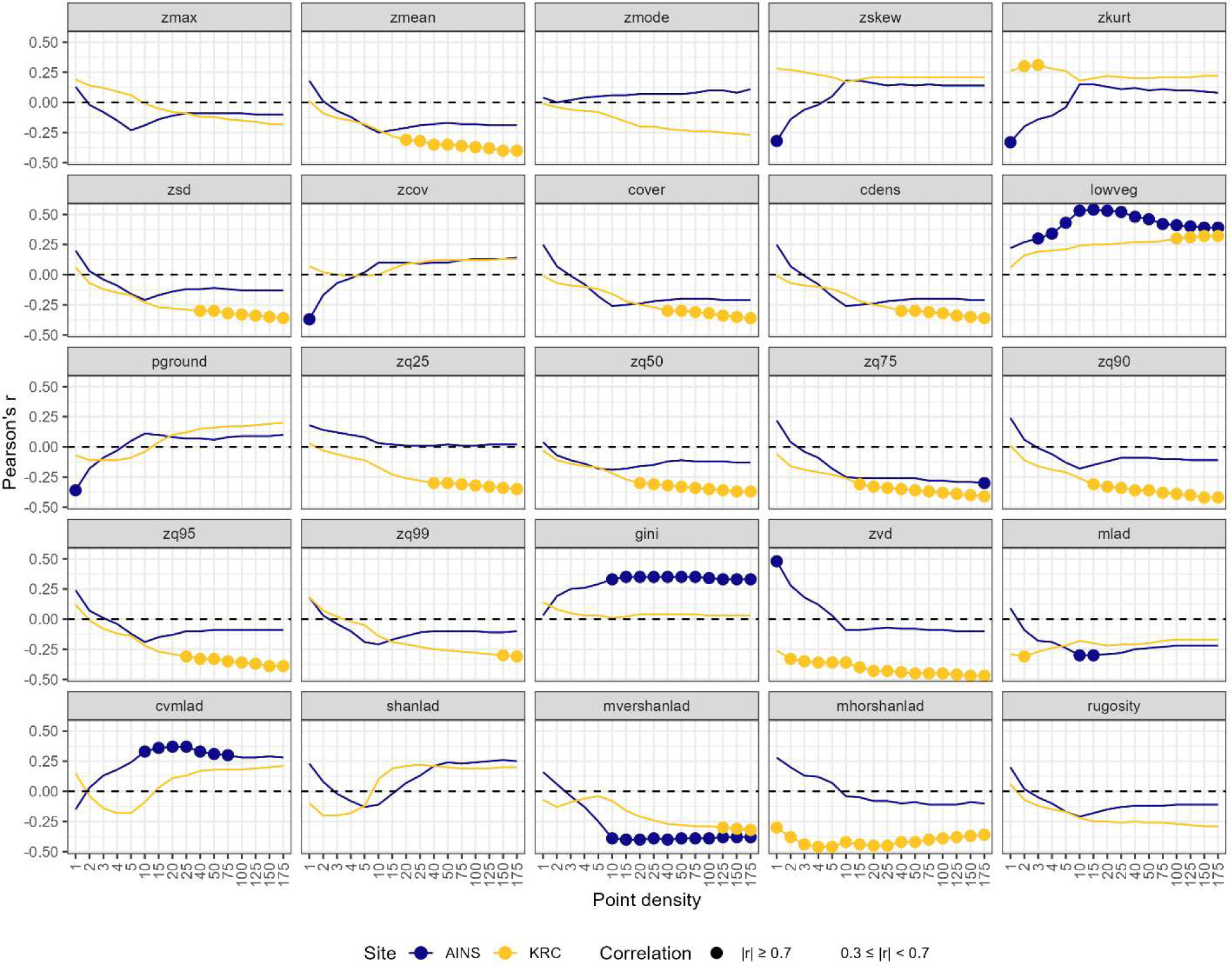
Pearson’s r for correlations between *mhorcvlad* and the 25 other forest structure metrics calculated from lidar with differing point densities at AINS (blue) and KRC (yellow). A filled circle indicates a strong correlation (|r| ≥ 0.7), while an open circle indicates a weak correlation (0.3 ≤ |r| < 0.7) and no circle indicates a negligible correlation (|r| < 0.3).

**Figure B2-22.**
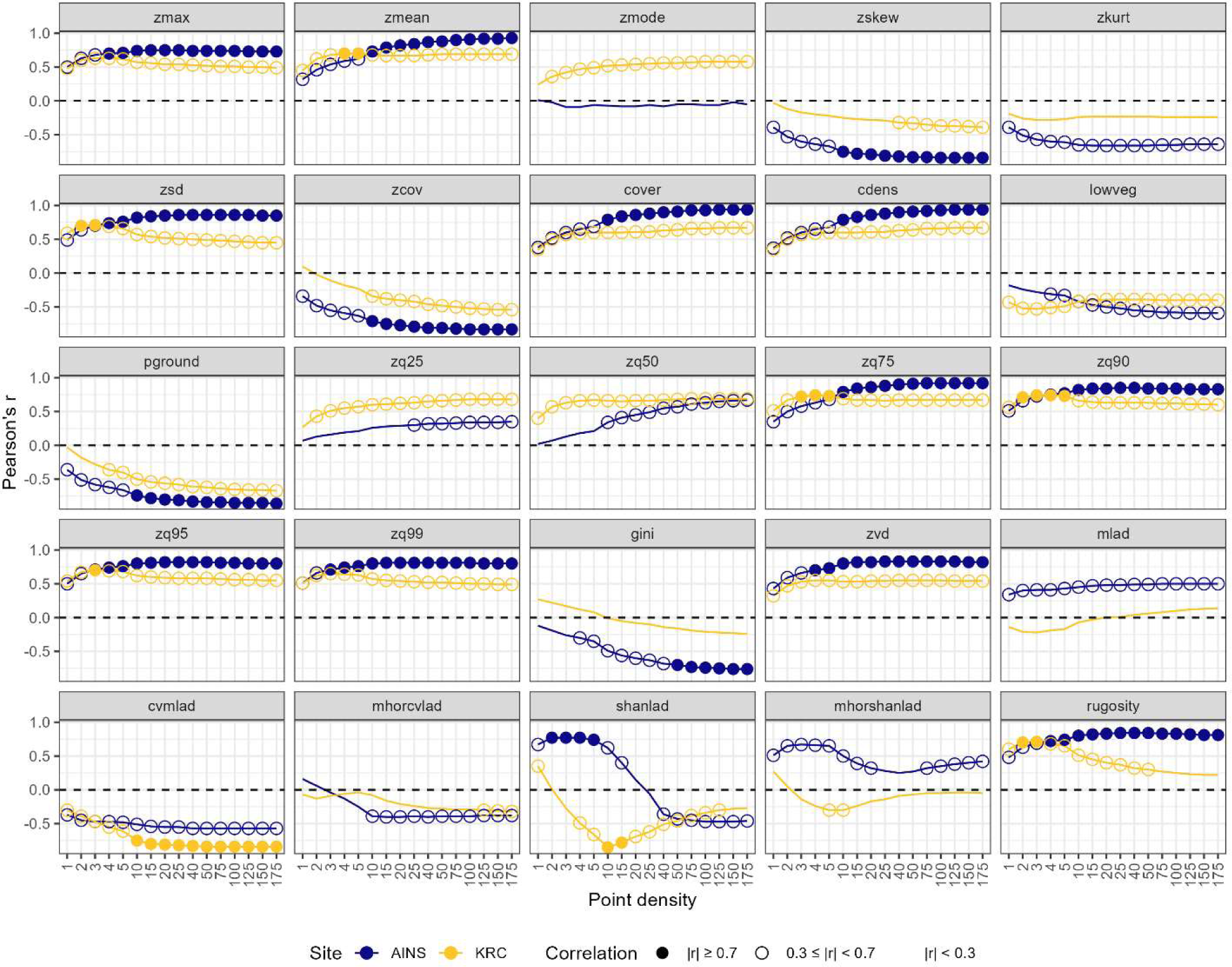
Pearson’s r for correlations between *mvershanlad* and the 25 other forest structure metrics calculated from lidar with differing point densities at AINS (blue) and KRC (yellow). A filled circle indicates a strong correlation (|r| ≥ 0.7), while an open circle indicates a weak correlation (0.3 ≤ |r| < 0.7) and no circle indicates a negligible correlation (|r| < 0.3).

**Figure B2-23.**
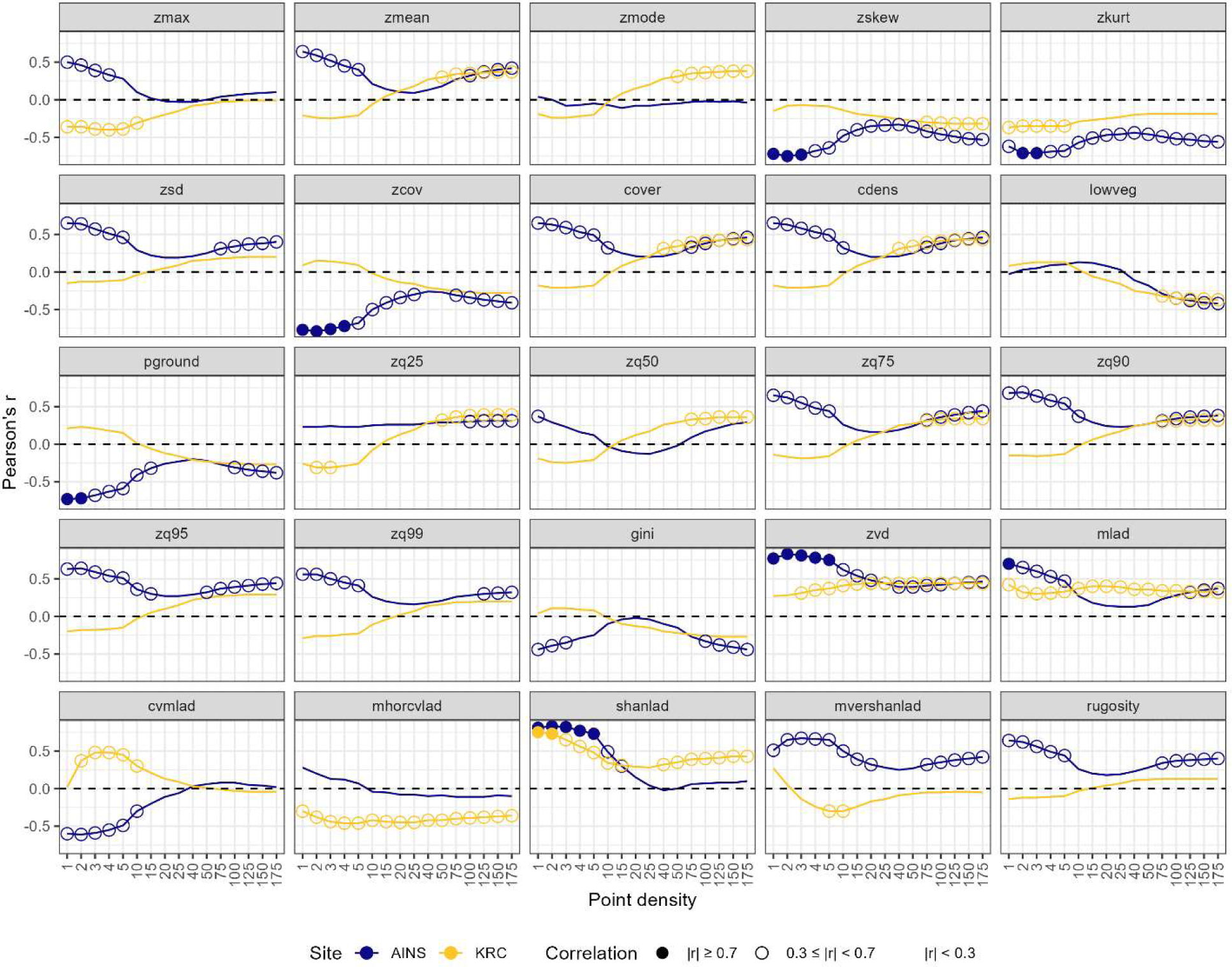
Pearson’s r for correlations between *mhorshanlad* and the 25 other forest structure metrics calculated from lidar with differing point densities at AINS (blue) and KRC (yellow). A filled circle indicates a strong correlation (|r| ≥ 0.7), while an open circle indicates a weak correlation (0.3 ≤ |r| < 0.7) and no circle indicates a negligible correlation (|r| < 0.3).

## References

1. Vashum, K.T.; Jayakumar, S. Methods to Estimate Above-Ground Biomass and Carbon Stock in Natural Forests-a Review. Journal of Ecosystem & Ecography 2012, 2, 1–7.

2. Fan, J.; Zhong, H.; Harris, W.; Yu, G.; Wang, S.; Hu, Z.; Yue, Y. Carbon Storage in the Grasslands of China Based on Field Measurements of Above- and below-Ground Biomass. Climatic Change 2008, 86, 375–396, doi:10.1007/s10584-007-9316-6.

3. Návar, J. Measurement and Assessment Methods of Forest Aboveground Biomass: A Literature Review and the Challenges Ahead. Biomass 2010, 27–64.

4. Cruz, M.G.; Alexander, M.E.; Wakimoto, R.H. Assessing Canopy Fuel Stratum Characteristics in Crown Fire Prone Fuel Types of Western North America. International Journal of Wildland Fire 2003, 12, 39–50.

5. Forbes, B.; Reilly, S.; Clark, M.; Ferrell, R.; Kelly, A.; Krause, P.; Matley, C.; O’Neil, M.; Villasenor, M.; Disney, M.;, et al. Comparing Remote Sensing and Field-Based Approaches to Estimate Ladder Fuels and Predict Wildfire Burn Severity. Frontiers in Forests and Global Change 2022, 5.

6. Lack, D. The Numbers of Bird Species on Islands. Bird Study 1969, 16, 193–209, doi:10.1080/00063656909476244.

7. MacArthur, R.H.; Wilson, E.O. The Theory of Island Biogeography; Princeton university press, 2001; Vol. 1; ISBN 0-691-08836-5.

8. MacArthur, R.H.; MacArthur, J.W. On Bird Species Diversity. Ecology 1961, 42, 594–598, doi:10.2307/1932254.

9. Bersier, L.-F.; Meyer, D.R. Bird Assemblages in Mosaic Forests: The Relative Importance of Vegetation Structure and Floristic Composition along the Successional Gradient. Acta Oecologica 1994, 15, 561–576.

10. Schwarzkopf, L.; Rylands, A.B. Primate Species Richness in Relation to Habitat Structure in Amazonian Rainforest Fragments. Biological Conservation 1989, 48, 1–12, doi:10.1016/0006-3207(89)90055-4.

11. Southwell, C.J.; Cairns, S.C.; Pople, A.R.; Delaney, R. Gradient Analysis of Macropod Distribution in Open Forest and Woodland of Eastern Australia. Australian Journal of Ecology 1999, 24, 132–143, doi:10.1046/j.1442-9993.1999.241954.x.

12. Vallan, D. Effects of Anthropogenic Environmental Changes on Amphibian Diversity in the Rain Forests of Eastern Madagascar. Journal of tropical ecology 2002, 18, 725–742.

13. Docherty, M.; Leather, S.R. Structure and Abundance of Arachnid Communities in Scots and Lodgepole Pine Plantations. Forest Ecology and Management 1997, 95, 197–207, doi:10.1016/S0378-1127(97)00024-8.

14. Halaj, J.; Ross, D.W.; Moldenke, A.R. Importance of Habitat Structure to the Arthropod Food-web in Douglas-fir Canopies. Oikos 2000, 90, 139–152.

15. Lang, N.; Kalischek, N.; Armston, J.; Schindler, K.; Dubayah, R.; Wegner, J.D. Global Canopy Height Regression and Uncertainty Estimation from GEDI LIDAR Waveforms with Deep Ensembles. Remote Sensing of Environment 2022, 268, 112760, doi:10.1016/j.rse.2021.112760.

16. Potapov, P.; Li, X.; Hernandez-Serna, A.; Tyukavina, A.; Hansen, M.C.; Kommareddy, A.; Pickens, A.; Turubanova, S.; Tang, H.; Silva, C.E.;, et al. Mapping Global Forest Canopy Height through Integration of GEDI and Landsat Data. Remote Sensing of Environment 2021, 253, 112165, doi:10.1016/j.rse.2020.112165.

17. Calders, K.; Verbeeck, H.; Burt, A.; Origo, N.; Nightingale, J.; Malhi, Y.; Wilkes, P.; Raumonen, P.; Bunce, R.G.H.; Disney, M. Laser Scanning Reveals Potential Underestimation of Biomass Carbon in Temperate Forest. Ecological Solutions and Evidence 2022, 3, e12197, doi:10.1002/2688-8319.12197.

18. Dubayah, R.O.; Sheldon, S.L.; Clark, D.B.; Hofton, M.A.; Blair, J.B.; Hurtt, G.C.; Chazdon, R.L. Estimation of Tropical Forest Height and Biomass Dynamics Using Lidar Remote Sensing at La Selva, Costa Rica. Journal of Geophysical Research: Biogeosciences 2010, 115, doi:10.1029/2009JG000933.

19. Oliveira, C.P. de; Ferreira, R.L.C.; da Silva, J.A.A.; Lima, R.B. de; Silva, E.A.; Silva, A.F. da; Lucena, J.D.S. de; dos Santos, N.A.T.; Lopes, I.J.C.; Pessoa, M.M. de L.;, et al. Modeling and Spatialization of Biomass and Carbon Stock Using LiDAR Metrics in Tropical Dry Forest, Brazil. Forests 2021, 12, 473, doi:10.3390/f12040473.

20. Asner, G.P.; Mascaro, J.; Anderson, C.; Knapp, D.E.; Martin, R.E.; Kennedy-Bowdoin, T.; van Breugel, M.; Davies, S.; Hall, J.S.; Muller-Landau, H.C.;, et al. High-Fidelity National Carbon Mapping for Resource Management and REDD+. Carbon Balance Manage 2013, 8, 7, doi:10.1186/1750-0680-8-7.

21. Erdody, T.L.; Moskal, L.M. Fusion of LiDAR and Imagery for Estimating Forest Canopy Fuels. Remote Sensing of Environment 2010, 114, 725–737, doi:10.1016/j.rse.2009.11.002.

22. Skowronski, N.S.; Clark, K.L.; Duveneck, M.; Hom, J. Three-Dimensional Canopy Fuel Loading Predicted Using Upward and Downward Sensing LiDAR Systems. Remote Sensing of Environment 2011, 115, 703–714, doi:10.1016/j.rse.2010.10.012.

23. Andersen, H.-E.; McGaughey, R.J.; Reutebuch, S.E. Estimating Forest Canopy Fuel Parameters Using LIDAR Data. Remote Sensing of Environment 2005, 94, 441–449, doi:10.1016/j.rse.2004.10.013.

24. Kramer, H.A.; Collins, B.M.; Kelly, M.; Stephens, S.L. Quantifying Ladder Fuels: A New Approach Using LiDAR. Forests 2014, 5, 1432–1453, doi:10.3390/f5061432.

25. Kramer, H.A.; Collins, B.M.; Lake, F.K.; Jakubowski, M.K.; Stephens, S.L.; Kelly, M. Estimating Ladder Fuels: A New Approach Combining Field Photography with LiDAR. Remote Sensing 2016, 8, 766, doi:10.3390/rs8090766.

26. Listopad, C.M.C.S.; Masters, R.E.; Drake, J.; Weishampel, J.; Branquinho, C. Structural Diversity Indices Based on Airborne LiDAR as Ecological Indicators for Managing Highly Dynamic Landscapes. Ecological Indicators 2015, 57, 268–279, doi:10.1016/j.ecolind.2015.04.017.

27. Davies, A.B.; Asner, G.P. Advances in Animal Ecology from 3D-LiDAR Ecosystem Mapping. Trends in Ecology & Evolution 2014, 29, 681–691, doi:10.1016/j.tree.2014.10.005.

28. Vogeler, J.C.; Hudak, A.T.; Vierling, L.A.; Evans, J.; Green, P.; Vierling, K.T. Terrain and Vegetation Structural Influences on Local Avian Species Richness in Two Mixed-Conifer Forests. Remote Sensing of Environment 2014, 147, 13–22, doi:10.1016/j.rse.2014.02.006.

29. Carrasco, L.; Giam, X.; Papeş, M.; Sheldon, K.S. Metrics of Lidar-Derived 3D Vegetation Structure Reveal Contrasting Effects of Horizontal and Vertical Forest Heterogeneity on Bird Species Richness. Remote Sensing 2019, 11, 743, doi:10.3390/rs11070743.

30. Froidevaux, J.S.P.; Zellweger, F.; Bollmann, K.; Jones, G.; Obrist, M.K. From Field Surveys to LiDAR: Shining a Light on How Bats Respond to Forest Structure. Remote Sensing of Environment 2016, 175, 242–250, doi:10.1016/j.rse.2015.12.038.

31. Jung, K.; Kaiser, S.; Böhm, S.; Nieschulze, J.; Kalko, E.K.V. Moving in Three Dimensions: Effects of Structural Complexity on Occurrence and Activity of Insectivorous Bats in Managed Forest Stands. Journal of Applied Ecology 2012, 49, 523–531, doi:10.1111/j.1365-2664.2012.02116.x.

32. Palminteri, S.; Powell, G.V.N.; Asner, G.P.; Peres, C.A. LiDAR Measurements of Canopy Structure Predict Spatial Distribution of a Tropical Mature Forest Primate. Remote Sensing of Environment 2012, 127, 98–105, doi:10.1016/j.rse.2012.08.014.

33. Vierling, K.T.; Bässler, C.; Brandl, R.; Vierling, L.A.; Weiß, I.; Müller, J. Spinning a Laser Web: Predicting Spider Distributions Using LiDAR. Ecological Applications 2011, 21, 577– 588, doi:10.1890/09-2155.1.

34. Müller, J.; Brandl, R. Assessing Biodiversity by Remote Sensing in Mountainous Terrain: The Potential of LiDAR to Predict Forest Beetle Assemblages. Journal of Applied Ecology 2009, 46, 897–905, doi:10.1111/j.1365-2664.2009.01677.x.

35. Müller, J.; Bae, S.; Röder, J.; Chao, A.; Didham, R.K. Airborne LiDAR Reveals Context Dependence in the Effects of Canopy Architecture on Arthropod Diversity. Forest Ecology and Management 2014, 312, 129–137, doi:10.1016/j.foreco.2013.10.014.

36. Lefsky, M.A.; Cohen, W.B.; Parker, G.G.; Harding, D.J. Lidar Remote Sensing for Ecosystem Studies: Lidar, an Emerging Remote Sensing Technology That Directly Measures the Three-Dimensional Distribution of Plant Canopies, Can Accurately Estimate Vegetation Structural Attributes and Should Be of Particular Interest to Forest, Landscape, and Global Ecologists. BioScience 2002, 52, 19–30, doi:10.1641/0006-3568(2002)052[0019:LRSFES]2.0.CO;2.

37. Asner, G.P.; Anderson, C.B.; Martin, R.E.; Knapp, D.E.; Tupayachi, R.; Sinca, F.; Malhi, Y. Landscape-Scale Changes in Forest Structure and Functional Traits along an Andes-to-Amazon Elevation Gradient. Biogeosciences 2014, 11, 843–856, doi:10.5194/bg-11-843-2014.

38. Pearse, G.D.; Watt, M.S.; Dash, J.P.; Stone, C.; Caccamo, G. Comparison of Models Describing Forest Inventory Attributes Using Standard and Voxel-Based Lidar Predictors across a Range of Pulse Densities. International Journal of Applied Earth Observation and Geoinformation 2019, 78, 341–351, doi:10.1016/j.jag.2018.10.008.

39. Kim, E.; Lee, W.-K.; Yoon, M.; Lee, J.-Y.; Son, Y.; Abu Salim, K. Estimation of Voxel-Based Above-Ground Biomass Using Airborne LiDAR Data in an Intact Tropical Rain Forest, Brunei. Forests 2016, 7, 259, doi:10.3390/f7110259.

40. Gobakken, T.; Næsset, E. Assessing Effects of Laser Point Density, Ground Sampling Intensity, and Field Sample Plot Size on Biophysical Stand Properties Derived from Airborne Laser Scanner Data. Can. J. For. Res. 2008, 38, 1095–1109, doi:10.1139/X07-219.

41. Rombouts, J.; Ferguson, I.S.; Leech, J.W. Campaign and Site Effects in LiDAR Prediction Models for Site-Quality Assessment of Radiata Pine Plantations in South Australia. International Journal of Remote Sensing 2010, 31, 1155–1173, doi:10.1080/01431160903380573.

42. LaRue, E.A.; Fahey, R.; Fuson, T.L.; Foster, J.R.; Matthes, J.H.; Krause, K.; Hardiman, B.S. Evaluating the Sensitivity of Forest Structural Diversity Characterization to LiDAR Point Density. Ecosphere 2022, 13, e4209, doi:10.1002/ecs2.4209.

43. Leitold, V.; Keller, M.; Morton, D.C.; Cook, B.D.; Shimabukuro, Y.E. Airborne Lidar-Based Estimates of Tropical Forest Structure in Complex Terrain: Opportunities and Trade-Offs for REDD+. Carbon Balance Manage 2015, 10, 3, doi:10.1186/s13021-015-0013-x.

44. Ruiz, L.A.; Hermosilla, T.; Mauro, F.; Godino, M. Analysis of the Influence of Plot Size and LiDAR Density on Forest Structure Attribute Estimates. Forests 2014, 5, 936–951, doi:10.3390/f5050936.

45. Magnussen, S.; Næsset, E.; Gobakken, T. Reliability of LiDAR Derived Predictors of Forest Inventory Attributes: A Case Study with Norway Spruce. Remote Sensing of Environment 2010, 114, 700–712, doi:10.1016/j.rse.2009.11.007.

46. Hansen, E.H.; Gobakken, T.; Næsset, E. Effects of Pulse Density on Digital Terrain Models and Canopy Metrics Using Airborne Laser Scanning in a Tropical Rainforest. Remote Sensing 2015, 7, 8453–8468, doi:10.3390/rs70708453.

47. Roussel, J.-R.; Caspersen, J.; Béland, M.; Thomas, S.; Achim, A. Removing Bias from LiDAR-Based Estimates of Canopy Height: Accounting for the Effects of Pulse Density and Footprint Size. Remote Sensing of Environment 2017, 198, 1–16, doi:10.1016/j.rse.2017.05.032.

48. Crespo-Peremarch, P.; Ruiz, L.Á.; Balaguer-Beser, Á.; Estornell, J. Analyzing the Role of Pulse Density and Voxelization Parameters on Full-Waveform LiDAR-Derived Metrics. ISPRS Journal of Photogrammetry and Remote Sensing 2018, 146, 453–464, doi:10.1016/j.isprsjprs.2018.10.012.

49. Almeida, D.R.A. de; Stark, S.C.; Shao, G.; Schietti, J.; Nelson, B.W.; Silva, C.A.; Gorgens, E.B.; Valbuena, R.; Papa, D. de A.; Brancalion, P.H.S. Optimizing the Remote Detection of Tropical Rainforest Structure with Airborne Lidar: Leaf Area Profile Sensitivity to Pulse Density and Spatial Sampling. Remote Sensing 2019, 11, 92, doi:10.3390/rs11010092.

50. Peng, X.; Zhao, A.; Chen, Y.; Chen, Q.; Liu, H. Tree Height Measurements in Degraded Tropical Forests Based on UAV-LiDAR Data of Different Point Cloud Densities: A Case Study on Dacrydium Pierrei in China. Forests 2021, 12, 328, doi:10.3390/f12030328.

51. Prata, G.A.; Broadbent, E.N.; de Almeida, D.R.A.; St. Peter, J.; Drake, J.; Medley, P.; Corte, A.P.D.; Vogel, J.; Sharma, A.; Silva, C.A.;, et al. Single-Pass UAV-Borne GatorEye LiDAR Sampling as a Rapid Assessment Method for Surveying Forest Structure. Remote Sensing 2020, 12, 4111, doi:10.3390/rs12244111.

52. National Park Service Assateague Island National Seashore: History and Culture 2021.

53. R Core Team R: A Language and Environment for Statistical Computing. 2021.

54. Roussel, J.-R.; Auty, D.; Coops, N.C.; Tompalski, P.; Goodbody, T.R.H.; Meador, A.S.; Bourdon, J.-F.; de Boissieu, F.; Achim, A. lidR: An R Package for Analysis of Airborne Laser Scanning (ALS) Data. Remote Sensing of Environment 2020, 251, 112061, doi:10.1016/j.rse.2020.112061.

55. Zhang, W.; Qi, J.; Wan, P.; Wang, H.; Xie, D.; Wang, X.; Yan, G. An Easy-to-Use Airborne LiDAR Data Filtering Method Based on Cloth Simulation. Remote Sensing 2016, 8, 501, doi:10.3390/rs8060501.

56. Adnan, S.; Valbuena, R.; Kauranne, T.; Gopalakrishnan, R.; Maltamo, M. Optimizing the Airborne Laser Scanning Estimation of Basal Area Larger than Mean (BALM): An Indicator of Cohort Balance in Forests. Ecological Indicators 2022, 142, 109162, doi:10.1016/j.ecolind.2022.109162.

57. de Almeida, D.; Stark, S.; Silva, C.; Hamamura, C.; Valbuena, R. leafR: Calculates the Leaf Area Index (LAD) and Other Related Functions. CRAN. https://cran.r-project.org/web/packages/leafR/index.html 2019.

58. Hagar, J.C.; Yost, A.; Haggerty, P.K. Incorporating LiDAR Metrics into a Structure-Based Habitat Model for a Canopy-Dwelling Species. Remote Sensing of Environment 2020, 236, 111499, doi:10.1016/j.rse.2019.111499.

59. Zhang, L.; Shao, Z.; Liu, J.; Cheng, Q. Deep Learning Based Retrieval of Forest Aboveground Biomass from Combined LiDAR and Landsat 8 Data. Remote Sensing 2019, 11, 1459, doi:10.3390/rs11121459.

60. d’Oliveira, M.V.N.; Broadbent, E.N.; Oliveira, L.C.; Almeida, D.R.A.; Papa, D.A.; Ferreira, M.E.; Zambrano, A.M.A.; Silva, C.A.; Avino, F.S.; Prata, G.A.;, et al. Aboveground Biomass Estimation in Amazonian Tropical Forests: A Comparison of Aircraft- and GatorEye UAV-Borne LiDAR Data in the Chico Mendes Extractive Reserve in Acre, Brazil. Remote Sensing 2020, 12, 1754, doi:10.3390/rs12111754.

61. van Aardt, J.A.N.; Wynne, R.H.; Oderwald, R.G. Forest Volume and Biomass Estimation Using Small-Footprint Lidar-Distributional Parameters on a Per-Segment Basis. Forest Science 2006, 52, 636–649, doi:10.1093/forestscience/52.6.636.

62. Knapp, N.; Fischer, R.; Huth, A. Linking Lidar and Forest Modeling to Assess Biomass Estimation across Scales and Disturbance States. Remote Sensing of Environment 2018, 205, 199–209, doi:10.1016/j.rse.2017.11.018.

63. Adnan, S.; Maltamo, M.; Mehtätalo, L.; Ammaturo, R.N.L.; Packalen, P.; Valbuena, R. Determining Maximum Entropy in 3D Remote Sensing Height Distributions and Using It to Improve Aboveground Biomass Modelling via Stratification. Remote Sensing of Environment 2021, 260, 112464, doi:10.1016/j.rse.2021.112464.

64. Knapp, N.; Fischer, R.; Cazcarra-Bes, V.; Huth, A. Structure Metrics to Generalize Biomass Estimation from Lidar across Forest Types from Different Continents. Remote Sensing of Environment 2020, 237, 111597, doi:10.1016/j.rse.2019.111597.

65. Renner, S.C.; Suarez-Rubio, M.; Kaiser, S.; Nieschulze, J.; Kalko, E.K.V.; Tschapka, M.; Jung, K. Divergent Response to Forest Structure of Two Mobile Vertebrate Groups. Forest Ecology and Management 2018, 415*–*416, 129–138, doi:10.1016/j.foreco.2018.02.028.

66. Cooper, W.J.; McShea, W.J.; Forrester, T.; Luther, D.A. The Value of Local Habitat Heterogeneity and Productivity When Estimating Avian Species Richness and Species of Concern. Ecosphere 2020, 11, e03107, doi:10.1002/ecs2.3107.

67. Adnan, S.; Maltamo, M.; Coomes, D.A.; Valbuena, R. Effects of Plot Size, Stand Density, and Scan Density on the Relationship between Airborne Laser Scanning Metrics and the Gini Coefficient of Tree Size Inequality. Can. J. For. Res. 2017, 47, 1590–1602, doi:10.1139/cjfr-2017-0084.

68. Van Ewijk, K.Y.; Treitz, P.M.; Scott, N.A. Characterizing Forest Succession in Central Ontario Using Lidar-Derived Indices. photogramm eng remote sensing 2011, 77, 261–269, doi:10.14358/PERS.77.3.261.

69. MacArthur, R.H.; Horn, H.S. Foliage Profile by Vertical Measurements. Ecology 1969, 50, 802–804.

70. Kamoske, A.G.; Dahlin, K.M.; Stark, S.C.; Serbin, S.P. Leaf Area Density from Airborne LiDAR: Comparing Sensors and Resolutions in a Temperate Broadleaf Forest Ecosystem. Forest Ecology and Management 2019, 433, 364–375, doi:10.1016/j.foreco.2018.11.017.

71. Gartside, P.S. A Study of Methods for Comparing Several Variances. Journal of the American Statistical Association 1972, 67, 342–346, doi:10.1080/01621459.1972.10482385.

72. Fox, J. Applied Regression Analysis and Generalized Linear Models; Sage Publications, 2015; ISBN 1-4833-2131-2.

73. Fox, J.; Weisberg, S.; Adler, D.; Bates, D.; Baud-Bovy, G.; Ellison, S.; Firth, D.; Friendly, M.; Gorjanc, G.; Graves, S. Package ‘Car.’ Vienna: R Foundation for Statistical Computing 2012, 16.

74. Abdi, H.; Williams, L.J. Tukey’s Honestly Significant Difference (HSD) Test. Encyclopedia of research design 2010, 3, 1–5.

75. Højsgaard, S.; Halekoh, U.; Robison-Cox, J.; Wright, K.; Leidi, A.A. doBy: Groupwise Statistics, LSmeans, Linear Contrasts, Utilities. R package version 2016, 4, 15.

76. Disney, M.I.; Kalogirou, V.; Lewis, P.; Prieto-Blanco, A.; Hancock, S.; Pfeifer, M. Simulating the Impact of Discrete-Return Lidar System and Survey Characteristics over Young Conifer and Broadleaf Forests. Remote Sensing of Environment 2010, 114, 1546– 1560, doi:10.1016/j.rse.2010.02.009.

77. Hirata, Y. The Effects of Footprint Size and Sampling Density in Airborne Laser Scanning to Extract Individual Trees in Mountainous Terrain.

78. Strunk, J.; Temesgen, H.; Andersen, H.-E.; Flewelling, J.P.; Madsen, L. Effects of Lidar Pulse Density and Sample Size on a Model-Assisted Approach to Estimate Forest Inventory Variables. Canadian Journal of Remote Sensing 2012, 38, 644–654, doi:10.5589/m12-052.

